# STEP: Spatial Transcriptomics Embedding Procedure for Multi-scale Biological Heterogeneities Revelation in Multiple Samples

**DOI:** 10.1101/2024.04.15.589470

**Authors:** Lounan Li, Zhong Li, Xiao-ming Yin, Xiaojiang Xu

## Abstract

In the realm of spatially resolved transcriptomics (SRT) and single-cell RNA sequencing (scRNA-seq), addressing the intricacies of complex tissues, integration across non-contiguous sections, and scalability to diverse data resolutions remain paramount challenges. We introduce STEP (Spatial Transcriptomics Embedding Procedure), a novel foundation AI architecture for SRT data, elucidating the nuanced correspondence between biological heterogeneity and data characteristics. STEP’s innovation lies in its modular architecture, combining a Transformer and *β*-VAE based backbone model for capturing transcriptional variations, a novel batch-effect model for correcting inter-sample variations, and a graph convolutional network (GCN)-based spatial model for incorporating spatial context—all tailored to reveal biological heterogeneities with unprecedented fidelity. Notably, STEP effectively scales the newly proposed 10x Visium HD technology for both cell type and spatial domain identifications. STEP also significantly improves the demarcation of liver zones, outstripping existing methodologies in accuracy and biological relevance. Validated against leading benchmark datasets, STEP redefines computational strategies in SRT and scRNA-seq analysis, presenting a scalable and versatile framework to the dissection of complex biological systems.

## Introduction

Developments in spatially resolved transcriptomics (SRT) and single-cell RNA sequencing (scRNA-seq) have revolutionized our comprehension of gene expression patterns. SRT, a fusion of histology and transcriptomics, offers precise transcript mapping within tissues, unveiling unprecedented details about tissue structure and gene expression profiles. Technologies like Visium ^1^ and Slide-seq V2 ^2^ have facilitated the collection of data at a non-single-cell resolution, capturing gene expression in ‘spots’ comprising multiple cells. Analyzing such data presents significant challenges in distinguishing various biological heterogeneities within these multicellular mixtures. In contrast, technologies such as Visium HD ^3^, MERFISH ^4, 5^ and STARmap ^6^ provide single-cell resolution in spatial transcriptomics, offering cell-specific gene expression data. These approaches play a crucial role in uncovering individual cellular identities and interactions, thereby enhancing our understanding of complex biological systems.

Accordingly, various computational tools are developed to solve these categories of problems in SRT data analyses: 1. Cell clustering, aiming to differentiate cells by cell types ^7-10^; 2. Spatial domain identification for discovery of biologically functional regions with a certain degree of spatial continuity in tissues ^11-21^. The significant improvements can be observed in solving these two tasks in the single-section scenario for a specific data resolution. Conversely, these tools are often not adaptive to the diverse data resolutions resulting from different SRT technologies. More importantly, there is a substantial challenge facing extant analytical methods due to the intricacies of the complex tissues whose regions are always unstructured and show a low level of spatial continuity. This can be particularly highlighted by liver zonation. Addressing this biological conundrum necessitates a methodology adept at not just capturing the full complexity of transcript information but also proficiently harnessing spatial context. Furthermore, the integration of data from multiple sections is required for understanding the complex mechanisms and characterizing the tissue environment. These sections are derived from different patients, covering various regions and conditions, such as distinctive treatments for the same ailment and genetic modifications like gene knock-outs. This reconciliation calls for computational methods capable of integrating spatial transcriptomic data from such disparate and non-contiguous slices. Regrettably, the current suite of tools often falls short of meeting these multifaceted demands.

These limitations facing the current methods arise from their excessive focus on specific tasks or data characteristics, that is, whether the data have single-cell resolution (“resolution orientation”) or are spatially resolved (“modality orientation”). These tendencies ignore the correspondence between the target underlying biological heterogeneities and source data resolutions and modalities. This vagueness further leads to the excessive focus on spatial variations but the underestimation of essential difference at transcriptional level. For example, some methods emphasize the correlation between spatial distances and the similarity of cell types as the core assumption of their cell clustering methods ^8, 9^. Not only are these assumptions incompletely correct, but they are also actually unnecessary because cell types can be well distinguished at the level of gene expression regardless of spatial locations. Also, for the integration of multiple sections, current methods often require the spatial contiguity between sections, whether vertically or horizontally ^14, 16, 18, 22-26^.Though, in concept, spatial domains demonstrate the spatial variations and the lower resolution compared to the cell types, the most fundamental variations that segment the regions remain transcriptional. Although one Bayesian model ^13^ and a cell-type annotation based method ^24^ are applicable to process both the multi-scale or multi-section (non-contiguous) SRT dataset, there is another limitation to which these methods are subject, that is, the inability to provide the embeddings that are consistent with their identified spatial domains for the downstream analyses, like pseudotime analysis, where the PCA and other batch-effect removal algorithms specific to scRNA-seq data ^27, 28^ are conducted, instead. In fact, such compromise leaves the spatial context aside the further biological explanations of the results. In summary, these causes can be attributed to the inability of the existing methods to balance the transcriptional, batch and spatial variations on different scales of biological heterogeneities. And to resolve this imbalance, a universal architecture is required to model the SRT data.

Considering this, the design of our solution starts with the clarification of the correspondences between target biological heterogeneities, source data resolution and source data modalities. Specifically, the scales of biological heterogeneities can be categorized into the cell type level and spatial domain level. Also, these two heterogeneities can be bound to data resolution at single-cell level and data modality of the spatially resolved, respectively. Moreover, the relationships between biological heterogeneities and variations can also well-established: with the transcriptional variation as the most essential one, the revelation of the cell type level heterogeneity requires the conservation of transcriptional variations but the elimination of batch variations, while of the spatial domain level one requires an extra incorporation of spatial variations.

Based on these foundational insights, we introduce STEP (Spatial Transcriptomics Embedding Procedure), a pioneering foundation deep learning architecture for SRT data that adopts a “decoupled but combinable” (modular) strategy to effectively address the aforementioned limitations. Our solution consists of three specialized models: 1. A Transformer ^29^ and *β*-VAE based backbone model (BBM), the core of STEP, designed for the precise extraction of transcriptional variations into low-dimensional embeddings, ensuring the scalability across diverse data resolutions; 2. A batch-embedding based paired transformation model (BEM), introducing an unparalleled batch-correction technique that operates independently from BBM and indicates a new direction in dealing with batch variations; and 3. A spatial model (SpM) leveraging the graph convolutional network (GCN) ^30^ distinctively applied to low-dimensional embeddings as a spatial filter or smoother rather than direct gene expression data as a feature extractor ^12, 14, 16, 18, 19, 25, 26.^ This innovative use of GCN significantly reduces noise, effectively capturing spatial contexts and variations with enhanced biological relevance. This gives full play to the power of GCN for its naturality as a spatial filter or smoother ^31^, allowing the major attention paid to the transcriptional features rather than being biased seriously by spatial variations. Ensuring a focused extraction of transcriptional variations, our decoupled strategy achieves a balanced attention across transcriptional, batch, and spatial variations. Meanwhile, the combinability of STEP allows for tailored analysis across diverse data resolutions and modalities according to the target heterogeneities to reveal, thus delineating multi-scale biological heterogeneities into consistent and comparative embeddings. Based on this unprecedented consistency and comparability, we also design an embedding-based cell type deconvolution algorithm with a spatial-domain-wise regularization, targeting the cell type level heterogeneity in non-single-cell resolution SRT data.

To demonstrate the versatility of STEP, its performance is evaluated across five distinct tasks with the combinations of three models based on target heterogeneities: 1. The integration of multiple batches to identify unified cell types or states. STEP is applied to both scRNA-seq data and single-cell resolution SRT data from multiple batches or platforms in this task, and adeptly handles clustering and batch effect removal in both embeddings and gene expressions. STEP achieves superior performance by employing the combo of BBM and BEM, outperforming the existing methods. 2. The analysis of single-batch non-single-cell resolution SRT data. In this scenario, STEP, with the combo of BBM and SpM, not only identifies the zones or regions within a tissue accurately, but also exhibits highly enhanced spatial continuity, smoothness and developmental trajectory to current methodologies. 3. The integration of multi-batch non-single-cell resolution SRT data. For the SRT dataset of this resolution, STEP makes use of the combo of all three models to efficiently align both contiguous and non-contiguous sections into a shared latent space. Also, the developmental trajectories are outlined for unified domains. This alignment enables comparative studies across tissues, which facilitates the identification and exploration of consistent biological features among different spatial domains under different conditions. 4. The integration of multi-batch single-cell resolution SRT data. For single-cell resolution SRT data, STEP can be applied not only to identify cell-types but also to aggregate the cell embeddings for identification of the spatial domains under the given spatial context, which reveals consistent biological heterogeneity on multiple scales. 5. By merging scRNA-seq and SRT data, STEP achieves efficient co-embedding and cell-type deconvolution. This approach amalgamates the strength of both data modalities, thus improving our understanding of complex biological structures. STEP is implemented based on the workflow SCANPY ^32^ in an extendable way, and is open-sourced with comprehensive tutorials available. Considering the architectural design of STEP, it has a massive potential in scaling to a Large Transcriptomic Model (LTM).

## Results

### STEP’s architecture and workflows

The STEP’s backbone model (BBM) is implemented based on Transformer and conceptualized on two fundamental biological principles: 1. Genes seldom operate in isolation but instead function as part of a group or module, such as a pathway. These gene modules comprise sets of genes collaborating in a coordinated manner to execute specific biological functions ^33, 34^. 2. Their behavior is intricately regulated through interactions with other genes within the same module and also with other gene modules. These regulatory interactions among gene modules involve diverse and complex mechanisms ^35-38^. BBM extends these principles by two corresponding key steps. First, it refers to multi view learning methods to map gene expression data into gene module sequences ^39, 40^. Second, it captures the interactions of gene modules with self-attention mechanisms, the core of Transformer. By this means, BBM ensures itself the capability to extract complex transcriptional variations from a regulatory perspective. During its forward propagation process, BBM first maps each z-scored gene expression data (a cell/spot) to a sequence of gene modules through learnable linear transformations, and then processes gene module sequences with several Transformer encoders. The outputs are extracted from BBM’s bottleneck layer as the low-dimensional embedding of each gene expression dataset. To further process these representations, we utilize an architecture resembling *β*-VAE ^41^ and incorporating the Zero Inflated Negative Binomial (ZINB) distribution to fit the raw gene expression data, wherein ZINB distribution has been corroborated to be highly effective in modeling gene-expression data. BBM subsequently fits the low-dimensional representations in standard normal distribution according to specialized standardization, i.e., reparameterization trick, and reconstructs them in the original data space via the decoder (Fig. 1a).

**Fig. 1.**
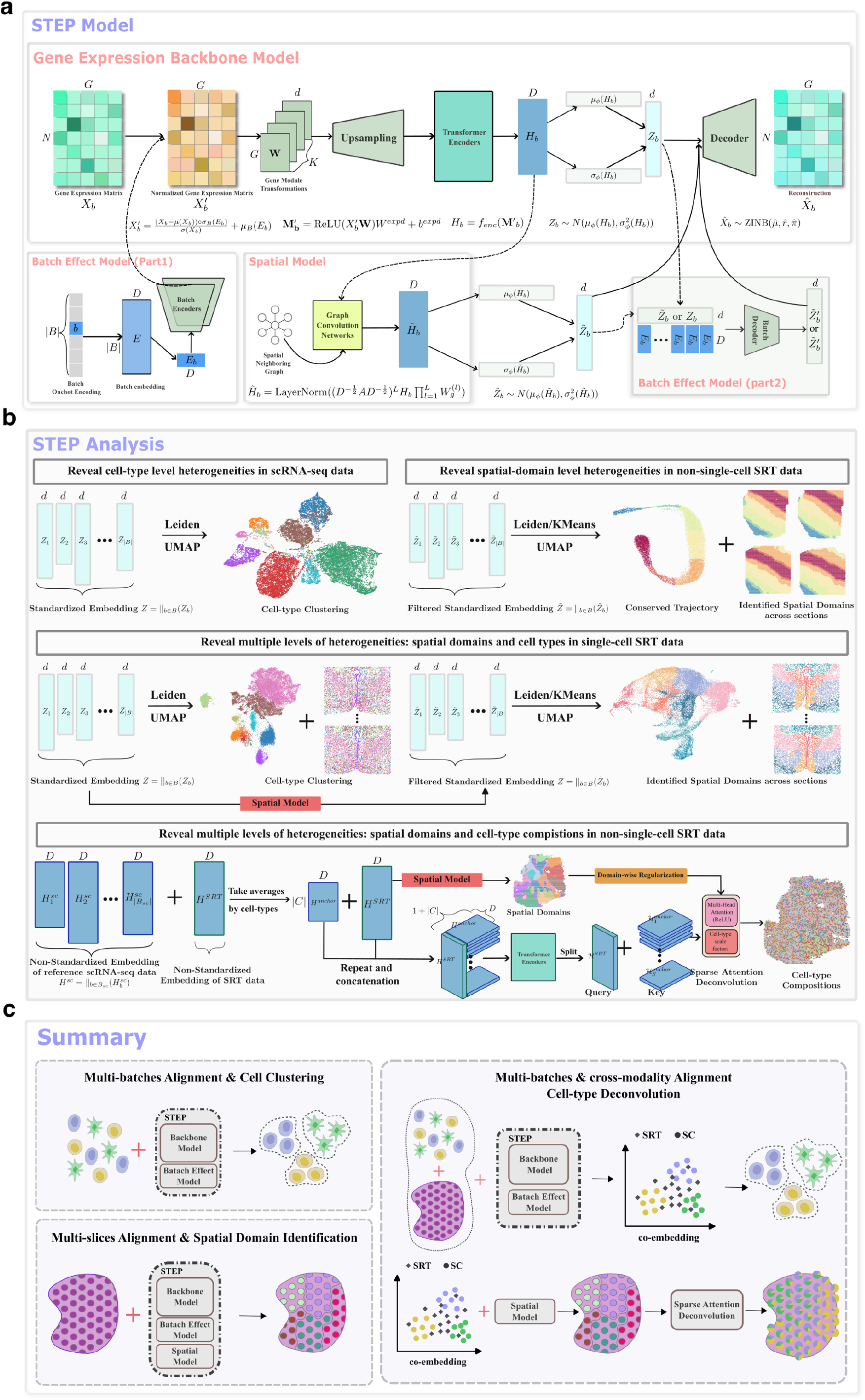
Overview of STEP’s architecture and workflows. **a**. The STEP framework consists of 3 main parts: backbone model, batch-effect model and spatial model and each part is tailored for different facts and issues encountered in the analysis of scRNA-seq data and SRT data; **b**. The compatible analyses with STEP’s embeddings of different data modalities and resolutions; First, STEP can integrate multiple batches of scRNA-seq data integration and correctly uncover the different cell types; Next, for non-single SRT data, STEP can align multiple sections and identify spatial domains across sections; Also, when it comes to the single-cell resolution SRT data, STEP can reveal both cell types and spatial domains across the tissues. The last is the integration across modality that co-embeds the multi-batch SC data and multi-batch SRT data and achieve cell-type deconvolution for the non-single-cell resolution SRT data; **c**. The corresponding model combinations and settings for different analyses.

Our newly proposed paired batch-effect model (BEM), drawing insights from current Pearson residual based normalization methods ^42, 43^ and the layer normalization ^44^ in deep learning, offers a new way to handle batch effects. By pairing such normalization with an “inverted” operation before the fitting of raw gene-expression data, BEM comprises two core components: 1. Batch Encoder with “BatchAwareLayerNorm”: the initial part addresses batch variations by modeling batch-dependent shifts and scales on observed gene expression data; 2. Batch Decoder with “BatchAwareScale”: analogously, an element-wise scale transformation in the latent space is applied before the decoding process (Fig. 1a). This BEM independently extends the BBM with a learnable batch-embedding concept, akin to essential word-embedding techniques widely used in natural language processing (NLP) methods. With this batch-embedding concept, the batch covariate can be effectively incorporated or regressed out for different purposes in a way akin to conditional generation (Supplementary Notes 1).

For the Batch Encoder, two MLPs use batch-embedding to estimate the batch-dependent shifts and scales of cells in each batch and perform element-wise addition and multiplication on the cell-wise z-scored gene expression data. The described process above is termed as “BatchAwareLayerNorm”. The Batch Decoder performs a batch-dependent element-wise scaling in the latent space, referred to “BatchAwareScale”, and it tries to retrieve the batch-variations before the decoding process to fit the raw counts after decoding. The specific workflow of BatchAwareScale is as follows: initially, a feedforward network is applied to the low-dimensional embedding and its corresponding batch embedding to produce a scaling vector the embedding. Then, the embedding is scaled by this scale vector and directed to the decoder. This modeling strategy delegates the extraction of transcriptional variation to the BBM, by placing the extraction of batch variations in the optimal timing or position: before and after the extraction of transcriptional ones, effectively decouples these two variations in the multi-batch/multi-section scenario.

We now introduce the forward propagation in the combination of BBM and BEM to show the co-work of these two models. Technically, for a gene expression dataset 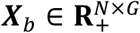 encompassing *N* cells, *G* genes sequenced, and belonging to batch *b* in the batch indicator set *B*, we initialize a set of linear transformations **W** ∈ **R**^*G*×*d*×*K*^ = {***W***_1,_***W***_2,_…,***W***_*K*_} where *d* is the dimension of the final standardized embedding (default *d* = 30). This set **W** is randomly initialized with a user-defined number of modules *K* (default *K* = 32) and a batch embedding ***E***_*B*_ ∈ **R**^|*B*|×*D*^ where *D* is the dimension of non-standardized embedding. For any cell/spot **x**_*B,n*_ ∈ ***X***_*B*_ belonging to batch *b*, we perform the following transformation to map it to a sequence of gene modules: ***M***_*B,n*_ = ReLU(BatchAwareLayerNorm (**x**_*B,n*_ onehot(*b*)***E***_*B*_)**W**).

Subsequently, we concatenate a randomly initialized “cls-token” ***z***^*cls*^ ∈ **R**^*d*^ to the sequence of gene modules to extract biological information and generate the final representation for each cell/spot. This concatenated sequence is then added by positional encodings ***E***_*pos*_ and fed into 3 Transformer encoders. Afterwards, an extra feedforward layer named “Readout module” is used to derive the non-standardized low-dimensional representation, denoted as **h**_*B,n*_ ∈ **R**^*D*^. This representation is then transformed to ***z***_*B,n*_ ∈ **R**^*d*^ using the reparameterization trick.

Before decoding ***z***_*B,n*_ back to original data space, the batch variations are retrieved for the alignment of reconstructions and raw gene expressions that indeed have batch-effects. Here, BatchAwareScale transforms the batch-independent embedding ***z***_*B,n*_ to a batch-dependent vector 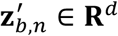.

As we model the gene expression data using a ZINB distribution, a non-linear decoder is employed to decode 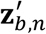 and to infer the parameters of the ZINB distribution for sampling the reconstructed data and calculating the negative log-likelihood (reconstruction error). The model training is based on the summation of the reconstruction error and the Kullback-Leibler (KL) divergence between the distribution of ***z***_*B,n*_ and standard normal, scaled by a small coefficient *β*, thereby forming a *β*-VAE (Fig. 1a). The representation 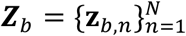 is the final representation of ***X***_*B*_, called standard-ized embedding.

Referring to our earlier introduction, GCN in our spatial model (SpM) can be termed as a specialized filter. It operates on a graph which is constructed by using spatial locations and BBM-extracted embedding. The SpM emphasizes principal signals and minimizes non-dominant cell signals throughout the model training process. By training the model in this manner, wherein data reconstruction occurs from neighbor-aggregated embedding (Fig. 1a), a form of spatial regularization is achieved. Consequently, the resulting SRT embedding produced by STEP is denoted as 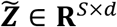 (we use tilde to represent the spatially filtered or smoothed embedding), where *S* locations are sequenced. This embedding demonstrates excellent performance in discerning spatial domains within tissues.

SpM is initially inserted between the calculations of Transformer encoders and reparameterization trick and is trained by tuning a pre-trained BBM (and BEM for multi-section case). There are two stages during the training process: 1. training a BMM (and BEM); 2. training the SpM together with the tuning of the BBM (and BEM). However, it is not necessary for non-single-cell resolution SRT data to capture the sole transcriptional variations. Due to the above reason, in our pursuit of optimizing model performance, we implement a contrastive learning strategy akin to the SimCSE ^45^. This strategic approach (see Methods), termed as “fast mode”, enhances the training process as an end-to-end style, accelerating the targeted extraction of spatial domain level heterogeneity.

After the training of STEP, we conduct the KMeans or Leiden algorithm on the embedding ***Z*** = ||_*B*∈*B*_ ***Z***_*B*_ for the clustering of cell types; and spatially refined 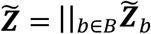, when the dataset is of SRT data, for the identification of spatial domains (Fig. 1b). In addition, for peeking cell type level heterogeneity in non-single-cell resolution SRT data, we integrate the workflows of two modalities by combining the annotated scRNA-seq data with the non-single-cell SRT data. We use just one BBM combined with BEM to map the data from both modalities into a shared space, termed as coembedding. Subsequently, we train a decoder specific for the SRT to remap it back to the original data space. Besides, a SpM is also trained for understanding the spatial context of SRT data. This integration enables us to comprehensively merge the data from both modalities while accomplishing tasks specific to each modality. Our target of this integrative analysis is to infer the cell type composition in each spot, referred to cell-type deconvolution, in a more biologically plausible way: spots in the same spatial domain share the similar cell-type compositions. With this biological observation and the comparative representation obtained from the co-embedding stage, we build our embedding-based deconvolution algorithm with domain-wise and gene-expression-wise regularizations (Fig. 1b and Methods).

### STEP consistently reveals multi-scale biological heterogeneities in single-cell resolution SRT data and integrates multiple batches of data

We first evaluate STEP’s BBM in two data, Human Colorectal Cancer (HCRC) and Mouse Small Intestine (MSI), from the newly proposed spatial transcriptomics technology called Visium HD by 10x Genomics ^46, 47^. This high-resolution spatial technology can resolve data at single-cell scale that consists of a grid with 2×2 µm barcoded areas, and the data is binned to 8×8 µm for analysis. The primary cell types and cellular components/structures are well identified by STEP’s BBM with the inspection of the corresponding markers. For the HCRC data (bin size: 16 µm) STEP’s BBM successfully deciphers cancer cells (cluster 0, 1, 6), cancer-associated fibroblasts (CAF) (cluster 3) and fibroblast cells (cluster 4) where the CAF cells surround and support the cancer cells as boundaries; goblet cells (cluster 2) surrounded by plasma cells (cluster 9) where they are co-locating in the intestine region and being walled by myofibroblasts cells (cluster 14); smooth muscle cells (cluster 7); endothelial cells (cluster 11) and epithelial cells (cluster 10) (Fig. 2a and Supplementary Fig. 1). For the MSI data (bin size: 8 µm), BBM also manages to illustrate the sophisticated structure of intestine. Firstly, two cell clusters form an organizational protruding structure where absorption and immune functions are closely juxtaposed: nutrient-absorbing villus tip enterocytes (cluster 1, 2) envelop immune-active plasma cells (cluster 5). Notably, two kinds of villus tip enterocytes (cluster 1 and 2) are differentiated by the gradient expression profile of Reg3b (Supplementary Fig. 2) along with the inner or outer segment of the intestine. Beneath, the intestine layer structure is also clearly revealed: the outermost layer, termed as the “protective epithelial barrier layer” (cluster 4); the second layer, composed of alternating clusters of “mucosal epithelial cells” (cluster 6) and “mixed immune-structural interface cells” (cluster 7); the middle layer, formed by “intestinal crypt secretory cells” (cluster 3); and the most innermost layer, termed as “muscularis propria” and composed of smooth muscle cells (cluster 0). (Fig. 2b and Supplementary Fig. 2). Another notable structure is lymphoid aggregates, including two big ones and a small one (cluster 8) where they are embedding between layers of intestinal crypt secretory cells and muscularis propria. Furthermore, the spatial domains are derived by applying SpM on the embeddings from each data. These spatial domains are biologically consistent with the original cell clustering results, which preserves the overall tissue structure and consolidates the corresponding cells into 8 primary regions (Fig. 2a, b). Conclusively, the results of these two data of ultra-high resolution and throughput significantly indicate the robustness and effectiveness of BBM in capturing the transcriptional variations and deciphering the complex cellular structures in tissue microenvironment with the new sophisticated SRT technology.

**Fig. 2.**
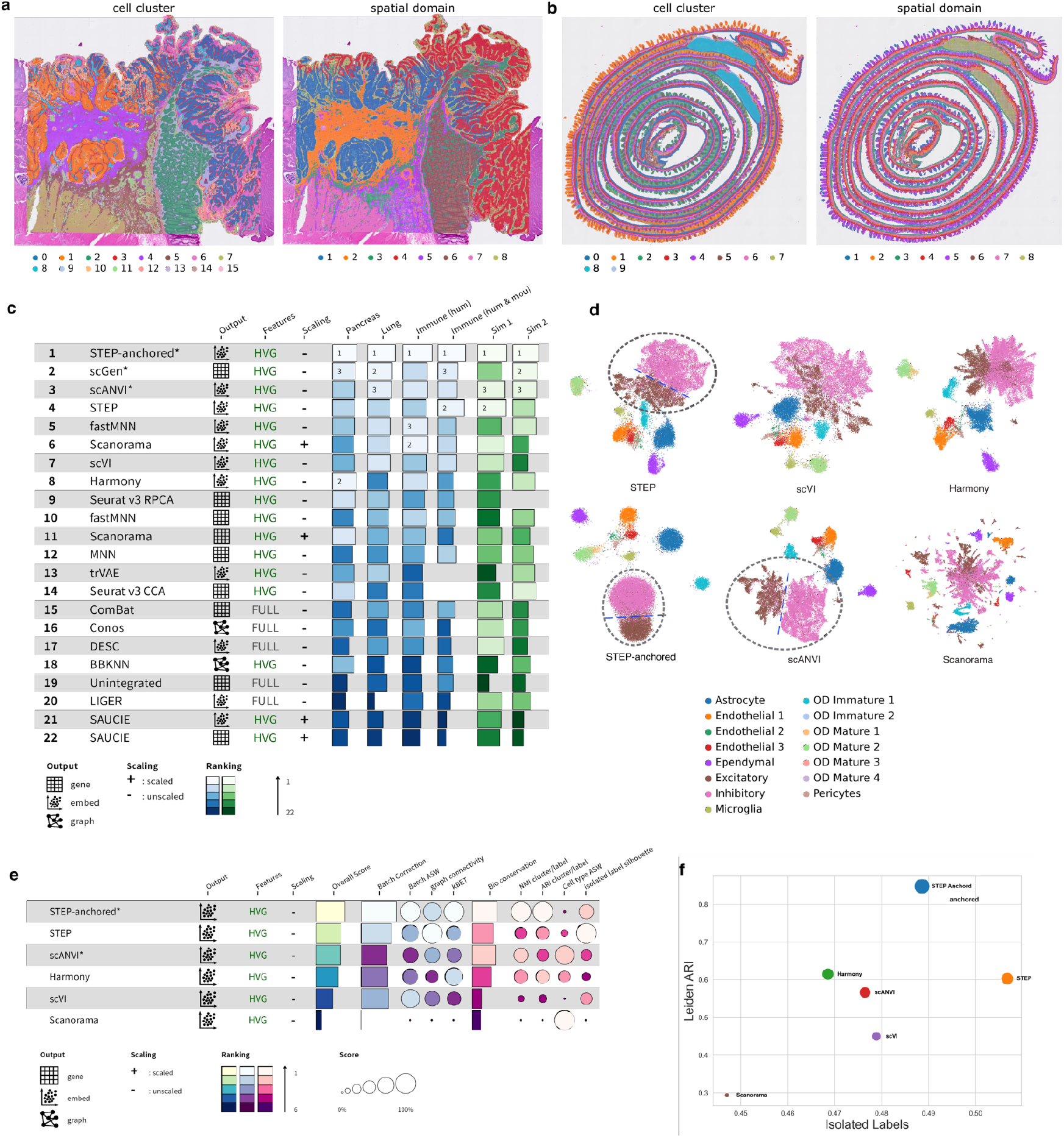
Analyses single-cell resolution data for the identifications of both cell type and spatial domain. **a**. Cell clustering and spatial domain identification results obtained from STEP for Human Colorectal Cancer (16 µm) by 10x Visium HD; **b**. Clustering results obtained from STEP for Mouse Intestine (8 µm) by 10x Visium HD; **c**. Ranking and bar plot for the performance of each tested methods with its the testing conditions (using HVGs only or all genes; scaled or not) and output style (graph, embedding or gene-expression); **d**. UMAP plots of results in MERFISH data, colored by cell-type annotation from original study. The differentiation of Excitatory and Inhibitory cells is highlighted with additional emphasis using dotted circles and straight dotted lines; **e**. Ranking and bar plot for the performances of STEP, STEP-anchored, scANVI, Harmony, scVI and Scanorama in a MERFISH dataset; **f**. Bubble plot of the clustering results obtained from each tested method. The x-axis represents the isolated-label silhouette, and the y-axis represents the ARI score for the clustering results using Leiden.

Echoing our prior discussion, in the context of multi-batch single-cell resolution data integration, variations across different data batches manifest in two primary forms: batch variations and biological/transcriptional variations. Regarding the two types of variations, batch variations denote technical shifts in sequencing data distribution, whereas biological variations reflect the inherent heterogeneities among cells in distinct cell types or states. With the aim to eliminate batch variations while preserving biological variations, we propose two solutions: 1. the combination of the BBM and BEM, referred as STEP for the simplicity. 2. STEP-anchored, where the BBM is further tuned by leveraging the pre-annotation when it is available. In the two methods above, the latter one, STEP-anchored, is operated under a semi-supervised learning paradigm: one “anchor point” is randomly initialized for each cell type as a cell-type-specific embedding and BBM is refined by introducing a classifier, which replaces the decoder after training STEP in the default self-supervised manner. This refinement process minimizes not only the KL-divergence but also the cross-entropy between the embedding of labeled cells and their respective labels, as well as every anchor point and its corresponding label. STEP-anchored combines pre-annotation results from all batches and enhances the model’s performance with a much more direct signal.

Evaluating data integration methods is a persistent challenge. However, utilizing the comprehensive benchmarking strategy established by ^48^, we manage to assess STEP across a spectrum of datasets. The benchmark encompasses two simulation datasets and four real atlas-level datasets (including Lung atlas, human pancreas, human immune, human & mouse immune datasets). This evaluation includes metrics for both batch effect removal and biological conservation. And its overall assessment criterion combines the Bio Conservation score (weighted at 0.7) and the Batch Correction score (weighted at 0.3).

Remarkably, STEP-anchored, leveraging pre-annotation information, demonstrates superior performance across all datasets for both batch effect removal and biological conservation metrics (Fig. 2c and Supplementary Table 1). Moreover, STEP, without relying on pre-annotation information, achieves the highest score in the first simulation dataset and the human & mouse immune atlas dataset, outperforming other methods including the ones utilizing available cell type annotations.

It’s important to highlight that within STEP-anchored, the “anchor points” efficiently anchor cell embeddings within their respective cell populations. This aspect notably bolsters the model performance in batch effect correction and biological conservation, although it may exhibit slightly less consistent trajectory conservation (Supplementary Fig. 3-5). Conversely, STEP demonstrates a well-balanced performance in integration and trajectory conservation (Supplementary Fig. 3-5). As a result, STEP-anchored shows promise as a tool for automated cell-type annotation, encompassing sub-types. Both STEP and STEP-anchored underscore the potency of our novel BEM combined with BBM, showcasing unmatched capabilities in capturing transcriptional variations and eliminating non-biological noise.

Regarding multi-section single-cell resolution SRT dataset, specifically the mouse hypothalamus data by MERFISH ^49^, we select five consistent tissue sections from Bregma distances of -0.04 mm (5488 cells), -0.09 mm (5557 cells), -0.14 mm (5926 cells), -0.19 mm (5803 cells), -0.24 mm (5543 cells) from the original study’s first animal. We utilize cell type annotations (15 cell types in total) from the original study as ground truth and exclude spatial location in this assessment. Comparing STEP against four other top-performing embedding-based integration methods (Fig. 2c): scVI ^50^, scANVI ^51^, Harmony ^27^, and Scanorama ^52^, STEP surpasses all these methods. The two methods successfully delineate a boundary between Excitatory cells and Inhibitory cells in the UMAP (Fig. 2d), a distinction that eludes the other methods. Additionally, we notice that a high value of principle components regression (PCR) often indicates an over-corrected state since the data itself does not appear to be affected by a severe batch-effect. Therefore, we exclude this metric from this test. In order to make up for this, we adjusted the weights of Batch Correction score to 0.4 and Bio Conservation score to 0.6, respectively. In specific evaluation, STEP-anchored adeptly integrates the five sections (KBET: 0.82, Silhouette: 0.97), and accurately defines cell types at the sub-type level through Leiden clustering (NMI: 0.85, ARI: 0.84). Meanwhile it achieves the highest scores for both Batch Correction (0.84) and Bio Conservation (0.67) with an aggregate score of 0.74 (Fig. 2e). STEP, without employing pre-annotation, closely follows STEP-anchored with a total score of 0.68 (Batch Correction 0.81; Bio Conservation 0.59; NMI 0.50; ARI 0.72). And it still outperforms the other four methods (Fig. 2e). Evaluation of biological conservation reveals that both STEP-anchored and STEP excel in discovering and conserving both common and rare cell populations (Fig. 2f).

### STEP accurately delineates spatial domains in human dorsolateral prefrontal cortex data in single-section scenario

In order to evaluate STEP’s performance in single-section SRT data with the combination of BBM and SpM, we employ the widely used DLPFC dataset ^53^ by Visium, encompassing 12 sections with manually annotated DLPFC layers and white matter (WM). Comparing STEP against GraphST ^14^, STAGATE ^16^, SpatialPCA ^15^, BayesSpace ^11^, and SpaGCN ^12^, we assess their proficiency in identifying spatial domains individually for each section. The accuracy of the identified spatial domains is quantified using the Adjusted Rand Index (ARI), measured against manual annotations (Fig. 3a). Additionally, we gauge the quality of identified domains for spatial continuity and smoothness using Moran’s I and Geary’s C metrics. These metrics rely on spatial neighboring graphs constructed via a k-Nearest-Neighbor (kNN) approach with *k* values of 1, 10, 20, and 30.

**Fig. 3.**
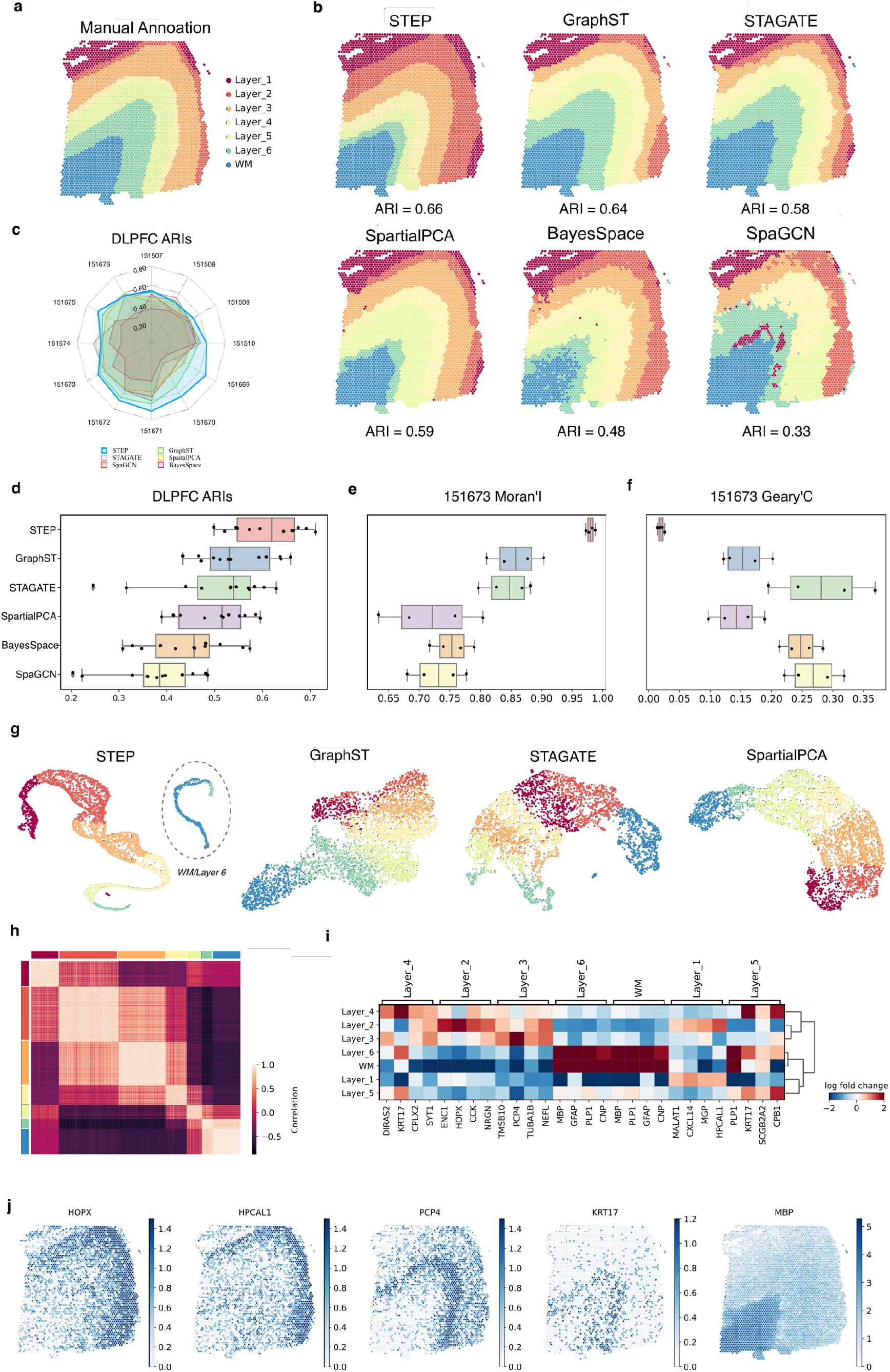
Analysis of human DLPFC data at single-batch-wise. **a**. Manual annotation from original study for section 151673; **b**. Spatial domains identified by each tested methods, with name at the top and ARI score at the bottom; **c**. Radar plots of the ARI scores for each method in each section; **d**. Box plots for the ARI scores of each method in each section; **e, f**. Box plots for Moran’s I and Geary’s C scores for the spatial domains in section 151673 obtained by each method; **g**. UMAP plots of embedding by STEP and compared methods; **h**. Heatmap of Pearson Correlation Coefficient by embeddings by STEP; **i**. Matrix plot of the log-fold-changes obtained by differentially expressed (DE) genes analysis based on STEP’s identified spatial domains; **j**. Expression of domain-specific genes obtained by Spatially DE genes analysis.

Particularly focusing on the results of section 151673, STEP’s identified domains are spatially smoother and more continuous compared to the other methods. GraphST, STAGATE, and SpatialPCA also exhibit smooth domains for several layers but fail to precisely capture the start of layer 2 as per manual annotations (Fig. 3b). The results from all sections further showcase STEP’s exceptional performance. For example, our method achieves the highest median ARI of 0.62 and an average ARI of 0.61 across all sections in single-batch analysis (Fig. 3c, d and Supplementary Fig. 6-8). Comparatively, GraphST, STAGATE, SpatialPCA, BayesSpace, and SpaGCN trail behind in median and mean ARI scores.

Besides, the quantitative measurement of spatial continuity via Moran’s I and Geary’s C further confirms the superior performance of STEP. Among all *k* values (1, 10, 20, 30), STEP consistently achieves the highest Moran’s I scores and the lowest Geary’s C scores, outperforming all the other methods (Fig. 3e, f). Specifically, STEP maintains a mean Moran’s I score of 0.98, and exhibits its superiority in ensuring spatial continuity compared to other evaluated methods (GraphST: mean Moran’s I = 0.86; STAGATE: mean Moran’s I = 0.84; SpatialPCA: mean Moran’s I = 0.72; BayesSpace: mean Moran’s I = 0.75; SpaGCN: mean Moran’s I = 0.73).

Beyond accurately identifying spatial domains, the underlying embeddings generated by STEP also clearly reveal the developmental trajectory of tissues (Fig. 3g and Supplementary Fig. 9-11). Employing UMAP for dimension reduction to visualize the embeddings of each method, the results demonstrate that STEP maintains a distinct stripe trajectory. Notably, STEP effectively delineates the WM/Layer 6 part. In contrast, STAGATE and spatialPCA illustrate the directional development relationship between domains but struggle to differentiate the WM part from other layers.

Further exploration of the embeddings and spatial domains derived from STEP involves analyzing the autocorrelation of spatial embeddings. The heatmap of Pearson Correlation (Fig. 3h) distinctly separates each domain, except for the WM and Layer 6, consistent with the UMAP visualization. Additionally, we conduct differential gene analysis between domains using the Wilcoxon rank sum test. This analysis successfully identifies several common spatially variable and domain-specific genes ^53^, such as HOPX and HPCAL1 for layer 2, PCP4 for layer 3, and KRT17 for layer 4 (Fig. 3i, j), further substantiating STEP’s proficiency in spatial domain identification.

### STEP uniquely aligns diverse contiguous and non-contiguous tissue sections

A distinguishing feature of STEP is its capability to synchronize various tissue sections, even if they are non-contiguous, thus enabling a remarkable integration at the level of spatial domain heterogeneity. This feature corroborates to be especially beneficial when working with intricate biological structures, such as sections from the same tissue type but from different sub-regions or donors. This significant advantage stems from STEP’s balanced modeling of transcriptional, batch and spatial variations through our modular design. By enabling a certain degree of independence between these three models, STEP facilitates section-independent embeddings for SRT data, despite the SpM operating on a section-by-section basis. This inherent feature underscores the flexibility and resilience of STEP.

At the outset, we employ STEP with the combo of all three models (BBM, BEM and SpM) to integrate multiple sections in the DLPFC data, and we focus on the first 4 sections (P1): 151507 (4226 spots), 151508 (4384 spots), 151509 (4789 spots), 151510 (4634 spots), totaling 18033 spots, which reveal a pattern akin to vertical alignment.

Comparatively, we evaluate BASS-Mult ^13^, PRECAST ^17^, STAligner ^25^ and MENDER ^24^, four methods that integrate sections without dependence on the positional relationships between the sections; and two image-aided methods: stGCL^18^ and STAIG^21^. By visualizing uniformly identified spatial domains in each section, STEP consistently displays ideal results akin to those observed in single-section analysis (Supplementary Fig. 6-11). Except MENDER fails to identify most spatial domains, the other five compared methods are all able to produce reasonable results at certain levels (Supplementary Fig. 12). Quantitatively, STEP showcases similar ARI scores to those obtained through single-section analysis, outperforming all the other methods in every section (STEP: mean ARI = 0.53; BASS-Mult: mean ARI = 0.44; PRECAST: mean ARI = 0.32; STAligner mean ARI = 0.49; stGCL mean ARI = 0.38; STAIG mean ARI = 0.20) (Fig. 4a and Supplementary Fig. 12). The spatial domains identified by STEP exhibit greater smoothness and continuity, supported by significantly higher and concentrated Moran’s I scores (Fig. 4b) and markedly lower and concentrated Geary’s C scores (Supplementary Fig. 13). Compared to STAligner and PRECAST, which are also embedding-based methods, a detailed examination of alignment results reveals that although both methods effectively remove batch effects, the trajectory from Layer 1 to Layer 6/WM is more accurately represented in the embedding by STEP than that by PRECAST (Fig. 4c). The UMAP visualization indicates that the STAligner’s and PRE-CAST’s embeddings lack developmental structure concerning manual annotation, underscoring the superior quality of embeddings produced by STEP. The other 2 groups in DLPFC dataset (P2: 151669 to 151672, P3: 151673 to 151676) are also tested. STEP yields an overwhelming performance on the p3 group as STEP completely resembles the layer 2 across 4 sections compared to manual annotations and achieves the highest ARI scores to date in every section (151673: 0.68; 151674: 0.67; 151675: 0.63; 151676: 0.63) in our knowledge (Supplementary Fig. 12). The spatially DE gene analysis based on identified domains reveals the coherent gene expression pattern for the same domain across sections. For instance, genes including GFAP, HPCAL, ENC1, NEFL, and TMSB10 exhibit high expression levels in Layer1 to Layer5, respectively (Supplementary Fig. 14, 15).

**Fig. 4.**
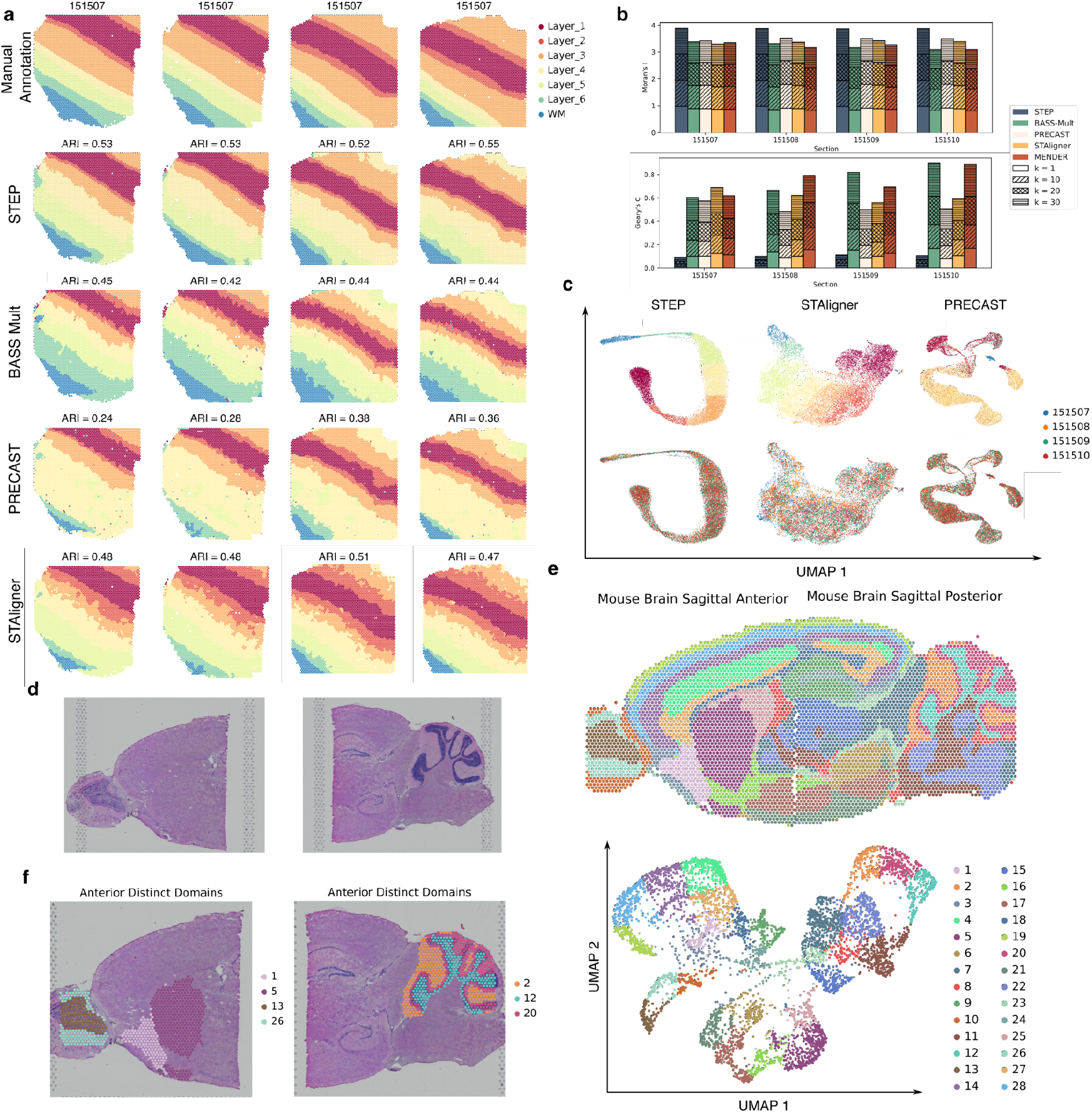
Analysis of Multiple-section alignment and spatial domains identification for human DLPFC, Mouse Brain sagittal anterior and posterior and mouse liver autophagy ATG7 KO by 10x Visium. **a**. Manual annotation and spatial domains identified each tested method: STEP, BASS-Mult, PRECAST, and STAligner with corresponding ARI scores (MENDER fails to identify the desired spatial domains); **b**. Moran’s I score for each method in each sample with each *k* value; **c**. UMAPs by embeddings obtained from STEP, STAligner and PRECAST, respectively, colored by spatial domain labels and section ids; **d**. Histology images for mouse brain sagittal anterior and posterior; **e**. Spatial domains identified by STEP across two sections (top panel) and UMAP colored by domain labels (bottom panel); **f**. Section-specific domains identified by STEP.

Subsequently, we investigate STEP’s efficacy in integrating horizontally aligned sections using data from the Mouse Brain’s sagittal anterior and posterior sections by 10x Visium. Despite being divided for sequencing, these sections, originating from physically connected tissues, exhibit distinct spatial domains due to variations in functional brain areas (Fig. 4d). The identification of 28 domains reveals a consistent appearance of domains across corresponding spatial locations referenced to the Allen Mouse Brain Atlas ^54^: the layers of the cerebral cortex are seamlessly connected across the two sections. Furthermore, distinct sub-regions in regions like the hippocampal formation (HPF), hypothalamus (HY), and Thalamus (TH) are successfully reconstructed (Fig. 4e). Embedding visualization illustrates five primary populations corresponding to MOB, CTX, CBX, HPF, and Caudoputamen (CP) regions (Fig. 4e).

These findings bear significant implications, showcasing STEP’s robust capability to seamlessly integrate tissue sections irrespective of their relative spatial relationships. Furthermore, these results pave the way for exploring transcriptional differences together with spatial context through the integrative analysis of multiple SRT datasets.

### STEP demonstrates the scalability in identifying spatial domains for single-cell resolution SRT from multiple sections

While STEP has significantly enhanced our capacity to accurately cluster cell types within single-cell resolution SRT data in previous introductions, emphasizing the identification of spatial domains remains an essential objective in research. The paramount advantage of spatial domains, conceptively compared to cell types, lies in their reinforcement of our comprehension regarding how cells operate collectively within their specific spatial context. This insight is crucial for understanding tissue structure, disease progression, and the biological mechanisms underlying various cellular processes. To further evaluate STEP, we first conduct tests on Mouse hypothalamus data by MERFISH and Mouse medial prefrontal cortex data by STARmap, focusing on spatial domain identification in a multi-section setting.

The first MERFISH dataset utilized in this investigation comprises five consecutive tissue sections extracted from the preoptic region of the mouse hypothalamus. Across all tissue sections, meticulous cell annotations are conducted by ^13^, identifying eight distinct structures: the third ventricle (V3), bed nuclei of the stria terminalis (BST), columns of the fornix (fx), medial preoptic area (MPA), medial preoptic nucleus (MPN), periventricular hypothalamic nucleus (PV), paraventricular hypothalamic nucleus (PVH), and paraventricular nucleus of the thalamus (PVT).

Due to the difference of the spot/cell spatial alignment between 10x Visium and MERFISH that the spots of 10x Visium form a regular hexagonal grid while a transcript unit in MERFISH is generated from image segmentation, meaning there is no regular alignment pattern for these units. As a result, the neighboring graph is constructed as in k-Nearest-Neighbors (kNN) method based on transcript units’ spatial locations. We set *k* = 30 (independent to the “*k*” in the evaluation of Moran’s I and Geary’s C) in kNN for this test, and again, STEP seamlessly eliminates batch-effects (Fig. 5a). In comparison with manual annotations, STEP still generates smooth spatial domains with clear boundaries and the high-level matches (Fig. 5b and Supplementary Fig. 16), even the resolutions of transcripts units increase a lot. However, STEP mis-clusters the bottom corner area of MPA domain to fx, and the upper area to MPN; and identifies more areas as PV than its annotation across sections. Considering these domains are adjacent, this mis-clustering may be caused by the relatively simple construction of the neighboring graph with kNN. BASS-Mult, as a method tailored for single-cell resolution SRT data, achieves similar performance with STEP, but is better at distinguishing of these adjacent domains: MPA, MPN and fx in an overall view (Bregma -0.14, Bregma -0.19). PRECAST fails to identify any domains except the V3, which may be caused by the fact that the identification target of PRECAST is the minimal unit of the transcripts in each dataset. When the resolution comes to single-cell level rather than non-single-cell level, PRECAST still tends to capture the heterogeneity of cell types rather than spatial domains. STAligner and MENDER also encounter a similar situation. And they can only identify the V3 or near V3 regions but fail to decipher the other regions. The first MERFISH dataset utilized in this investigation comprises five consecutive tissue sections extracted from the preoptic region of the mouse hypothalamus. Across all tissue sections, meticulous cell annotations are conducted by ^13^, identifying eight distinct structures: the third ventricle (V3), bed nuclei of the stria terminalis (BST), columns of the fornix (fx), medial preoptic area (MPA), medial preoptic nucleus (MPN), periventricular hypothalamic nucleus (PV), paraventricular hypothalamic nucleus (PVH), and paraventricular nucleus of the thalamus (PVT).

**Fig. 5.**
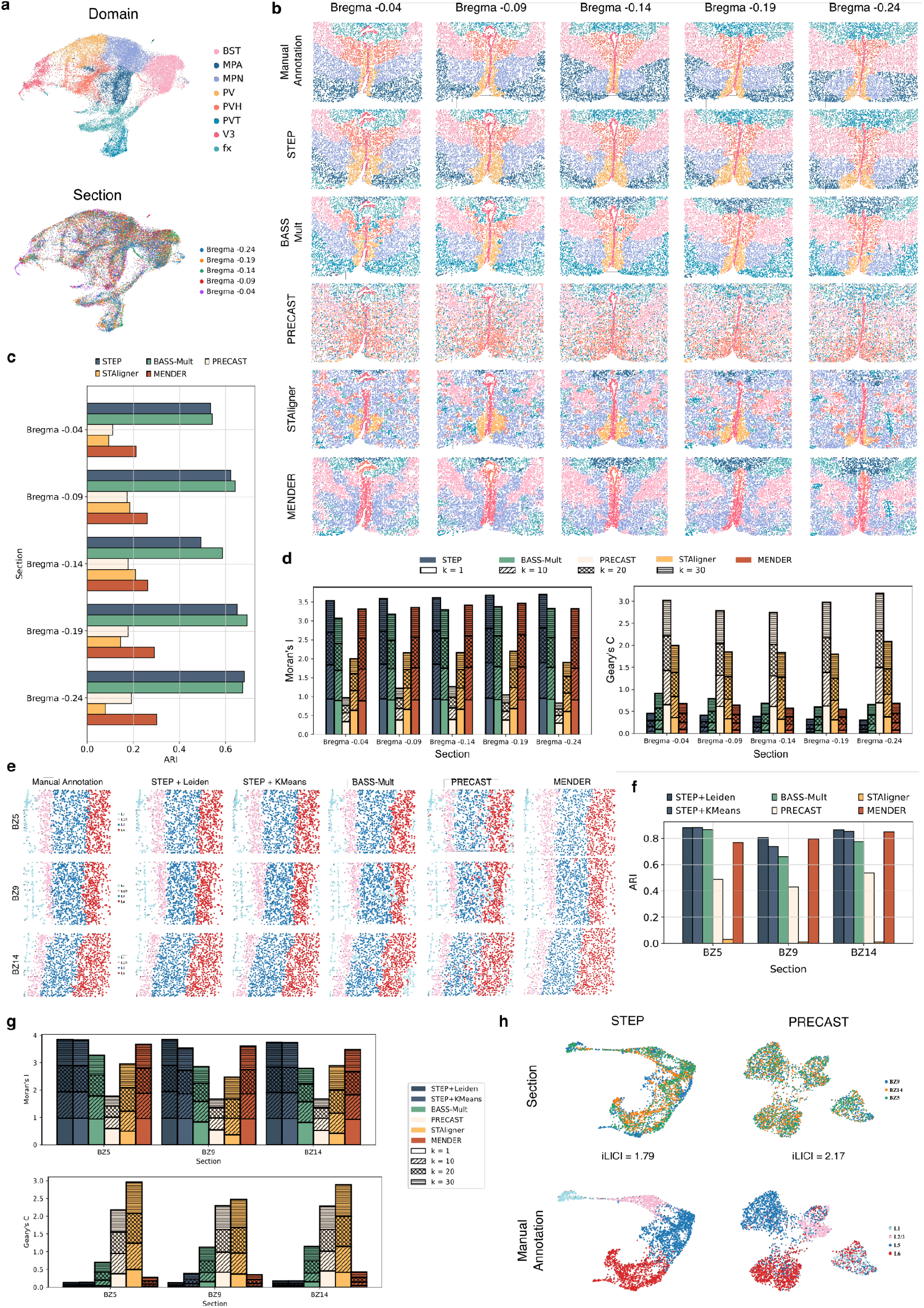
Analysis of multi-section alignment and spatial domain identification in Mouse hypothalamus by MERFISH and Mouse medial prefrontal cortex by STARmap. **a**. UMAP plot obtained by STEP and colored by spatial domains and section ids; **b**. Comparison for the manual annotation and spatial domains identified by each test methods: STEP, BASS-Mult, PRECAST, STAligner and MENDER; **c**. Bar plots for ARI scores of each methods in each section by MERFISH; **d**. Stacked bar plots of Moran’s I and Geary’s C scores of each method in each section by MERFISH with each *k* value; **e**. Comparison for the manual annotation and spatial domains identified by each test methods: STEP+KMeans, STEP+Leiden, BASS-Mult, PRECAST, STAligner and MENDER; **f**. Bar plots for ARI scores of each method in each section by STARmap; **g**. Stacked bar plot of Moran’s I and Geary’s C scores from each method in each STARmap section with each *k* value; **h**. Comparison of the UMAP results obtained from the embeddings of STEP and PRECAST, colored by manual annotation and section ids, respectively.

**Fig. 6.**
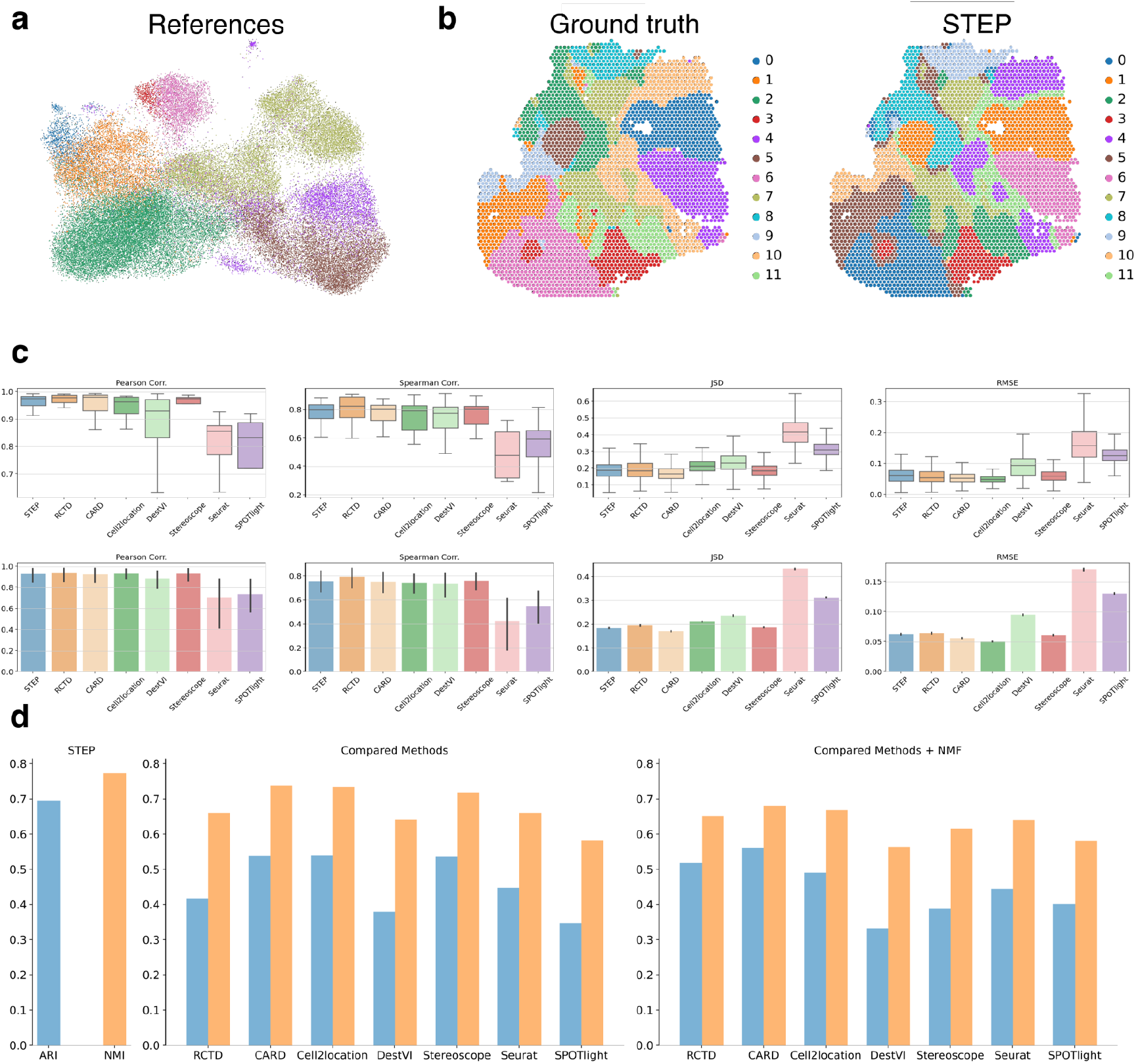
Analysis of the spatial domain identification and deconvolution on simulated SRT data. **a**. UMAP plot of the reference scRNA-seq data; **b**. Left: spatial plot of ground truth of spatial domains. Right: spatial plot of domains identified by STEP; **c**. Comparison for the accuracy of deconvolution through 4 metrics: PCC, SCC, JSD and RMSE (top: Boxplot, bottom: Bar plot); **d**. ARI and NMI scores for spatial domains inferred from each method.

From a metrics perspective, STEP achieves ARI scores that are comparable across sections when matched against BASS-Mult, although slightly weaker in the first 4 sections (Fig. 5c) (STEP: -0.04 ARI = 0.53, -0.09 ARI = 0.62, -0.14 ARI = 0.50, -0.19 ARI = 0.65, -0.24 ARI = 0.68; BASS-Mult: -0.04 ARI = 0.54, -0.09 ARI = 0.64, -0.14 ARI = 0.59, -0.19 ARI = 0.69, -0.24 ARI = 0.67). Contrastively, the smoothness of spatial domains by STEP is still dominant as the scores of Moran’s I and Geary’s C, outperforming other 2 methods in all cases of *k* values (Fig. 5d) (average Moran’s I by STEP: Bregma -0.04 = 0.89, Bregma -0.09 = 0.90, Bregma -0.14 = 0.90, Bregma -0.19 = 0.92, Bregma -0.24 = 0.93; average Moran’s I by BASS-Mult: Bregma -0.04 = 0.77, Bregma -0.09 = 0.90, Bregma -0.14 = 0.82, Bregma -0.19 = 0.85, Bregma -0.24 = 0.83). We also conduct spatially DE gene analysis on the spatial domains identified by STEP to find domain-specific genes, and the results shows a reasonable overlap with original studies: Cd24a and Nnat1 enriched in V3 domain; Slc18a2 enriched in PV domain; Esr1 enriched in PV, MPN and BST domains; and MBP, Ermn enriched in MPA domain (Supplementary Fig. 17).

Additionally, we also test STEP in the 7-th animal from the same original study with sections ranging widely from Bregma distance 0.26 to -0.29, in total 12 sections (0.26, 0.21, 0.16, 0.11, 0.06, 0.01, -0.04, -0.09, -0.14, -0.19, -0.24, -0.29). This dataset is used to test if STEP can capture the dynamic changes of domains, not only the changes of domain types but also the changes of area for the same domain type, with the changes of distance and angles along the tissue. From the results, STEP effectively identifies anterior commissure (ACA) domain present at 0.26 to -0.14, as well as the area of ACA changes from disconnected to connected status (Supplementary Fig. 18-24). And the identified V3 domain is at the bottom of the section at beginning, and it gradually “grows” up with the changes of Bregma distance along the tissue and reaches the top from -0.09 section (Supplementary Fig. 20, 21). In the meantime, identified PV domain begins with a larger connected area and is gradually “squashed” by the V3 domain to edge-like area (Supplementary Fig. 20, 21). The spatially DE results further support these developments (Supplementary Fig. 22-24). This test strongly validates the capacity of STEP to handle complex series of sections, and even the data resolution is single-cell level.

Next, we test STEP, BASS-Mult and PRECAST on the Mouse medial prefrontal cortex data generated by STARmap. This series of sections (3 sections in total: BZ5, BZ9, BZ14) consist of 4 domains including Layer1, Layer2/3, Layer5, Layer6. Besides testing the identification of spatial domains, we also evaluate the robustness of embeddings when working with different clustering methods: Leiden and KMeans. Referring to the manual annotations, the spatial domains across all sections identified by STEP with two clustering methods show that they all, overall, work well with embeddings generated by STEP. This phenomenon further demonstrates the robustness and biological significance of the embeddings (Fig. 5e and Supplementary Fig. 25). However, STEP with Kmeans mis-clusters a set of cells in the Layer 6 domain as Layer 1 in section BZ9 (Fig. 5e). Such mis-clustering situation can also be found in the domains obtained by BASS-Mult and PRECAST, but they mis-cluster not only this set of cells but also the cells in Layer 6 of all sections. In terms of quantity, the ARI scores (Fig. 5f) show a decent matching between manual annotation and STEP with Leiden (BZ5: 0.88; BZ9: 0.80; BZ14: 0.86) and STEP with Kmeans (BZ5: 0.88; BZ9: 0.74; BZ14: 0.85) as well as competitive performances by BASS-Mult (BZ5: 0.87; BZ9: 0.66; BZ14: 0.77) and MENDER (BZ5: 0.77; BZ9: 0.80; BZ14: 0.85). PRECAST overall conserves the domain-level biological heterogeneity (BZ5: 0.49; BZ9: 0.42; BZ14: 0.53), but STAligner completely fails to identify the layer structure (Supplementary Fig. 26). The scores of Moran’s I and Geary’s C with different *k* values are significantly higher, which once again prove the advantages of the spatial domain generated by STEP in terms of purity or spatial continuity (Fig. 5g). We further compare STEP and PRECAST in the view of batch-effect correction measured by the “integration Local Inverse Simpson Index” (iLISI). Even though PRECAST achieves a higher iLICI score (2.17) than STEP (1.79), we can, by visualizing the manual annotations in the UMAPs, observe the significant over-correction by PRECAST and its loss of developmental trajectory (Fig. 5h). Conversely, embeddings by STEP do not segregate apart but form a trajectory along the developmental orders of domains (Fig. 5h). It is more reasonable since the spatial domains in this data, unlike the cell types, exhibit biological continuity because of the coarser graininess of biological heterogeneity and the nature of the development of the tissue. Additional spatially DE genes analysis successfully locates the underlying genes that form the identified spatial domains (Supplementary Fig. 27).

### STEP seamlessly combines scRNA-seq data with spatial transcriptomics data, yielding intricate spatial domains and detailed deconvolution outcomes

To conduct a joint analysis of scRNA-seq data and SRT data, STEP co-embeds both modalities of data into a shared latent space. In this approach, a domain is conceptualized as a potentially ambiguous cell type, suggesting that spots within the same domain may possess similar cell type compositions, referred as ***P***_*d*_ ∈ **R**^|*C*|^ for domain *d*. The representation of ***P***_*d*_, modeled as a random variable, reflects the average cell-type composition within that domain. Instead of directly inferring the distribution and parameters of ***P***_*d*_, STEP implicitly incorporates this information within the embedding of SRT data as a regularization term. This regularization term is computed as the sample variance of estimated cell-type composition ***P***_*d*_ within the same domain. STEP primarily relies on the embedding and the resultant domains for cell-type deconvolution. However, it is notable that similar studies in image deconvolution have demonstrated that sampling from the latent space can lead to blurred reconstructions of the original data space images ^55^, a phenomenon also observed in cell-type deconvolution studies ^56^. To address this challenge, STEP integrates a reconstruction error directly derived from the gene expression level as an additional regularization technique in the deconvolution process. This dual-reconstruction approach separates the deconvolution problem into two sub-problems: 1. identifying cell types occurrences and 2. inferring coarse cell-type compositions in the spots through the latent space embedding, which are then refined in the gene-expression space (see Methods).

We initiate our deconvolution performance assessment with simulated data, labeled as pseudo-SRT. This simulation process involves a hierarchical strategy conducted on the reference scRNA-seq dataset (Fig. 6a) and an actual tissue data (Fig. 6b). The simulation commences by assigning an anticipated cell type composition (CTC) to each previously identified domain (Fig. 6b). We refer to such CTCs as domain-specific CTC. In this process, the CTC at each location is sampled from a normal distribution, using the mean equivalent to the corresponding domain-specific CTC. Moreover, we implement spatial smoothing on the sampled CTC for the simulation of spatial context. Next, each gene expression of pseudo-SRT is generated by convolving the sampled scRNA-seq data (gene expressions) with each CTC. The final step involves multiplying each generated gene expression data by a randomly sampled cell number from a uniform distribution, with predefined lower and upper bounds of 10 and 20 (see Methods).

STEP is benchmarked against several other methods including CARD ^57^, Cell2location ^58^, RCTD ^59^, SPOTLight ^60^, Stereoscope ^61^, DestVI ^56^, and Seurat ^62^ using simulation data. Metrics including Pearson correlation coefficient (PCC), Spearman correlation coefficient (SCC), Jensen–Shannon divergence (JSD), and mean squared root (MSE) are utilized for evaluation. Notably, STEP obtains competitive scores across all metrics: top-2 highest average PCC scores and SCC scores (STEP: PCC=0.93, SCC=0.76; RCTD: PCC=0.94, SCC=0.79; CARD: PCC=0.93, SCC=0.76; Cell2location: PCC=0.93, SCC=0.75; DestVI: PCC=0.88, SCC=0.73; Stereoscope: PCC=0.93, SCC=0.76; Seurat: PCC=0.71, SCC=0.42; SPOTlight: PCC=0.74, SCC=0.55); the 2 lowest average JSD scores (STEP: JSD=0.19; RCTD: JSD=0.20; CARD: JSD=0.17; Cell2location: JSD=0.21; DestVI: JSD=0.24; Stereoscope: JSD =0.19; Seurat: JSD =0.43; SPOTlight: JSD =0.31); and the top-3 lowest MSE scores (STEP: MSE=0.06; RCTD: MSE=0.06; CARD: MSE=0.06; Cell2location: MSE=0.05; DestVI: MSE=0.09; Steeoscope: MSE=0.06; Seurat: MSE=0.17; SPOTlight: MSE=0.13) (Fig. 6c and Supplementary Fig. 28, 29).

We also inspect the ability of each method to represent the spatial domains. Due to the inability of a direct inference for the spatial domains by the compared methods, we use two ways to infer the spatial domains from each CTC estimated by corresponding deconvolution methods: 1. Labeling the domain of each spot by the dominant cell types; 2. Labeling the domain of each spot by the argmax of the coefficients obtained by non-negative matrix factorization (NMF). The number of spatial domains labeled by the first way is not same with the ground truth due to the difference in the number of cell types and true spatial domains, while the latter labeling way yields the same number of spatial domains with ground truth by setting the number of components in NMF equal to the target number of spatial domains. ARI and NMI scores are then calculated to quantify the accuracy of inferred spatial domains. As a result, STEP achieves the highest scores in both metrics (ARI=0.70, NMI=0.77), outperforming all the other methods with both two kinds of inferred spatial domains. The ARI scores of compared methods are all lower than 0.55 and the NMI scores are all lower than 0.74 (Fig. 6d and Supplementary Fig. 30).

### STEP decrypts liver zonation and cellular compositions across multiple non-contiguous sections

The examination of non-contiguous section integration is conducted using 6 samples from human liver using 10x Visium. Besides, as previously mentioned, the micro-environment in livers is particularly complex, thereby the desired spatial domains are more functional and less spatially continuous. The datasets in previous tests of STEP are mostly about brains, of which the microenvironment and structures are simpler and more well-organized. This liver dataset accounts for a total of 15120 spots where four are derived from normal human, and the other two are from mice with Biliary atresia (BA) disease ^63, 64^. STEP is tasked to identify 5 spatial domains, successfully locating the desired Zone 1 (domain 4), Zone 2 (domain 3) and Zone 3 (domain 1) regions consistently across sections (Fig. 7a, b and Supplementary Fig. 31, 32). The rest two spatial domains are spatially arranged such that domain 5 is encompassed by the boundary of the domain 2, likely the immune region surrounded by the endothelial region. We further examine the gene expression profiles in each spatial domain by performing the DE gene analysis. Results show a mixture of markers’ expressions related to Cholangiocytes (AQP1, SOX9, KRT7, FXYD2), Endothelial cells (PECAM1, CCL21, TIMP1, VCAM1), hepatic stellate cells/Fibroblasts (DCN, PTGDS) and Immune cells (IGKC, IGHA1, IGHG1, MYL9) in domain 2 and domain 5 (Supplementary Fig. 31, 32). Incorporating the spatial locations and the gene expression profiles, we annotate domain 2 and 5 together as “Immune & Endo regions” for simplicity.

**Fig. 7.**
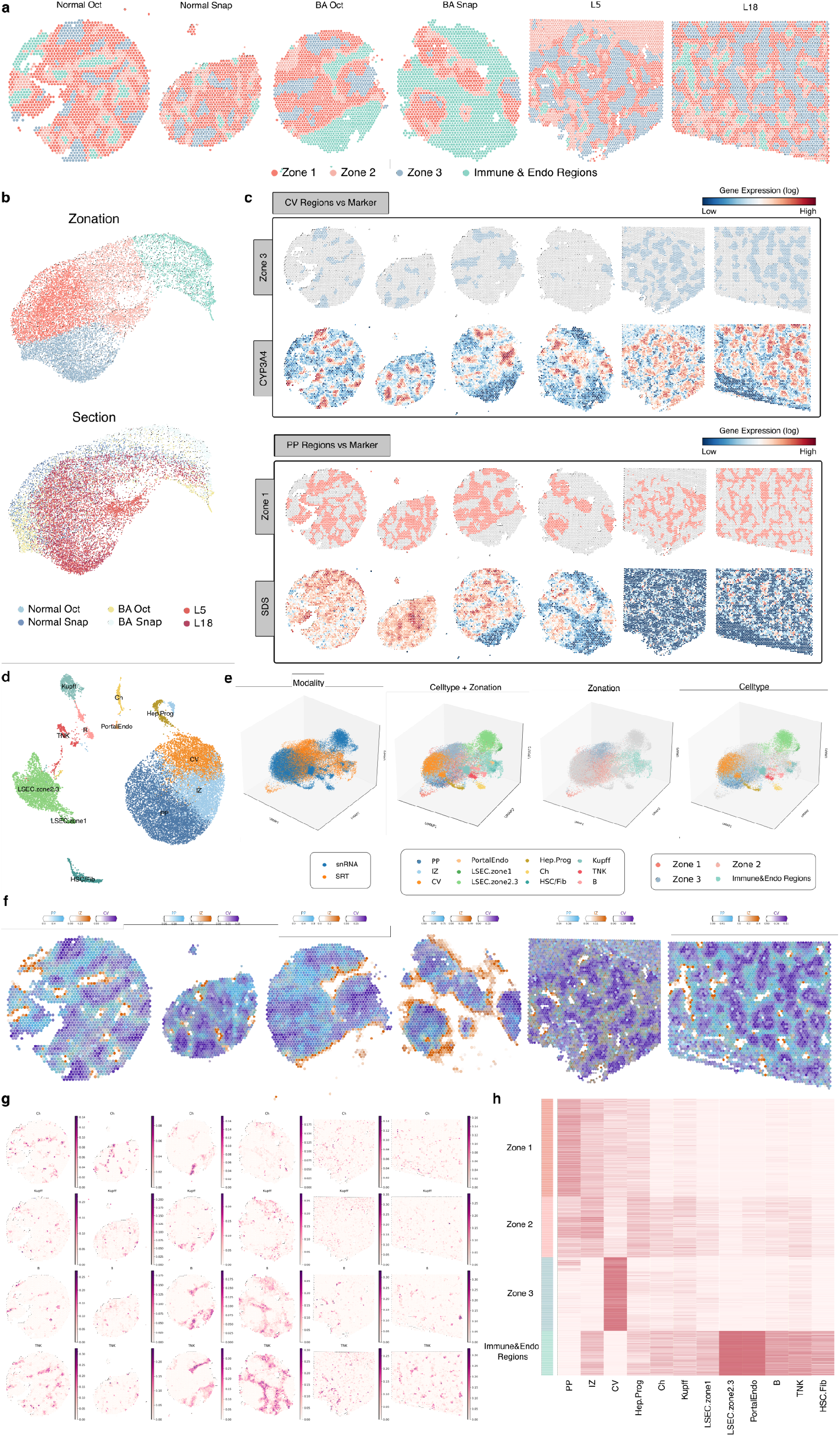
Comprehensive spatial transcriptomics analysis of liver tissue using STEP. **a**. Unified liver zonation results from six non-contiguous liver sections, simultaneously obtained using STEP. Three main distinct zones (periportal/Zone 1, intermediate/Zone 2, and pericentral/Zone 3) across the liver tissue are successfully identified by STEP; **b**. UMAP visualization of embeddings generated by STEP, colored by zones and section ids, respectively; **c**. Comparison of spatial patterns between Zone 1 (peri-portal) and Zone 3 (pericentral) as identified by the model (top row), with corresponding marker gene expression patterns (bottom row). Color intensity indicates logarithm transformed expression level; **d**. MAP plot of single-nucleus RNA-seq reference data from liver tissue, colored and annotated by cell types; **e**. Three-dimensional UMAP visualization of co-embedded SRT and snRNA-seq data, colored by data modality, cell-type annotation and zonation; **f**. Spatial plots showing the deconvolution results of peri-portal (PP), intermediate zone (IZ), and central vein (CV) hepatocyte subtypes across the six liver sections. Each plot represents a liver section, with spots colored according to the proportions of PP, IZ, and CV hepatocytes as determined by STEP’s deconvolution algorithm. The color intensity at each spot indicates the relative abundance of each hepatocyte subtype in that spatial location; **g**. Spatial plots of abundances of Cholangiocytes, Kupffer, B and T/NK cells in each spot inferred by STEP; **h**. Heatmap showing the relationship between liver zones (rows) and cell-type compositions/abundances (columns).

Comparison between spatial patterns of identified zones and the corresponding markers ^65-68^ exhibits a high-level match and resemblances, strongly indicating the biological accuracy of STEP in liver zonation task (Fig. 7c). The other four methods, BASS, PRECAST, STAligner and MENDER are also tested on this liver zonation task. However, they all fail to identify unified domains across all sections (Supplementary Fig. 33, unable to handle the technical shifts between sections or spatial biases within sections and the complexity of liver microenvironment. As we navigate the complex terrain of hepatic zonation, STEP stands out as a pioneering computational method, marking the first instance of an analytical tool that can delineate the three primary zones of the liver with accuracy.

In order to make a profound study on cellular functions within each zone, we use STEP to perform cell type deconvolution using snRNA-seq data from the same study as reference. This approach allows STEP to dissect multi-scale biological heterogeneities, a capacity that not only enables the precise differentiation of liver zones, but also uncovers the inherent complexities within different cell types, especially in the complicated “Immune & Endo Regions” as we defined before. Following the processing steps from the original study, we first annotate the data with three zonal hepatocytes: periportal (PP), interzonal (IZ) and central vein (CV); along with hepatocytes progenitor (Hep. Prog), cholangiocytes (Ch), liver sinusoidal endothelial cells zone 1 (LSEC zone 1), LSEC zone 2&3, T/NK cells, B cells, Kupffer cells (Kupff), hepatic stellate cells/fibroblasts (HSC/Fib), and portal endothelial cells (PortalEndo) (Fig. 7d). Subsequently, we use STEP to perform an unsupervised co-embedding procedure on snRNA-seq data and SRT data upon which our deconvolution algorithm is built as we described before. The results of co-embedding again exhibit the powerful capability of STEP in capturing the biological variations appearing in the data even from different data modalities and resolutions (Fig. 7e). From the perspective of single modality, whether for snRNA-seq or SRT data, the inherent biological heterogeneity structures, i.e. cell-type level for snRNA-seq and spatial-domain level for SRT data, are remarkably conserved (Fig. 7e). More importantly, the related cells and spots, whose belonging annotations refer to the same biological substances (like CV cell type and CV region), are locating together and forming continuous areas in the embedding space (Fig. 7e). Given this solid co-embedding procedure, STEP consistently infers the cell-type composition in each spot in biological wise. The abundances of three primary zonal hepatocytes, PP, IZ and CV, are spatially complementary and dominating in the corresponding areas according to the previous zonation, meanwhile in proper gradient patterns (Fig. 7f). In addition, the expected Cholangiocytes, Kupffer cells, B cells and T/NK cells are all found enriching in the “Immune & Endo Regions” (Fig. 7g and Supplementary Fig. 34). BA samples contain larger areas of immune cells compared to the other 4 normal samples, which is coherent to the disease condition and provide a more detailed view than spatial domains. We also inspect an overall landscape of zones and cell type abundances. With the blind spatial domain regularization, meaning STEP only knows “anonymous” groups of spots when proceeding deconvolution process, STEP achieves a balanced and accurate result in cell-type deconvolution as the inferred cell type abundances clearly form the expected zones (Fig. 7h).

With STEP’s proficiency in identifying these zones and cell type compositions, we gain a comprehensive view that not only reinforces the established understanding of hepatic biology but also extends it by adding depth to the spatial continuum of liver functionality. Such detailed zonal mapping is of immense value in the study of hepatic diseases, where understanding the zonation can provide a deeper insight of zonal susceptibility, disease progression, and potential therapeutic interventions.

## Discussion

The proposed foundation AI architecture for SRT data, STEP, effectively decouples the modeling of gene-expression, batch-effect and spatial correlation, and balances these variations. In this way, STEP exhibits its capacity on the integration of multiple batches/sections, identification of different grained heterogeneities (cell types and spatial domains) and generalization to different resolution of transcriptomics data. STEP also successfully bridges the gap between spatial domain identification and cell-type deconvolution, providing a unified framework to better decode the intricate biological systems. These are notable features that are different from most current analytical solutions that primarily fail to balance the different sources of variation.

In more detail, the novelty of STEP’s backbone model lies in its ability to model gene expression data using learnable gene module transformations together with Transformer, which allows a much more complex capture of transcriptional variations. And using GCN as a spatial filter in STEP’s spatial model enables fast and effective identification of spatial domains by preserving the principal biological signals in a cell or spot. Compared to existing graph neural networks (GNNs) based methods like GraphST, STAGATE and SpaGCN, STEP not only performs better identification of spatial domains, but also demonstrates more excellent consistency in model utilization with the definition of spatial domains. Moreover, our newly proposed paired batch-effect model further enhances the robustness of STEP when facing multiple batches or sections, even in the cross-modality situation. This elaborately designed architecture enables STEP to fully capitalize on interaction within gene modules and spatial information within the data and can be easily generalized to multiple batches scenario, a facet lacking in current techniques. Our approaches, therefore, contribute to a more nuanced understanding of both non-single-cell and single-cell resolution spatial transcriptomics data. Of considerable significance is STEP’s performance across multiple tasks, including cell type identification and data integration in various batches, platforms and protocols; spatial domains identification and integration of both contiguous and non-contiguous sections; and cell-type deconvolution in integrative analysis across modalities. STEP exhibits a notable proficiency in dealing with varying complexities of dataset management, showcasing a higher level of accuracy compared to the existing solutions.

Despite the promising results, several areas still require further investigation. For instance, embeddings are not fully utilized in cell-type deconvolution, the gene expression imputation performance is not as remarkable as other tailored methods, and our method relies on other selection methods for highly variable genes (HVGs) ^43, 69, 70^ or spatially variable genes (SVGs) ^71^. These constraints might primarily stem from a lack of a more powerful decoder and a superior pretext task beyond simple gene-expression profile reconstruction. Accordingly, the potential inconsistency in STEP’s encoding and decoding processes is also evident. While the encoding part is designed to model underlying biological effects, the decoding part is solely focused on modeling the statistical properties of observed gene expressions. Ideally, the decoding process should also model gene modules’ interactions, even though this might intensify the computational load. Furthermore, there are methods based on the Transformer and trained within a self-supervised paradigm ^72, 73^. However, these methods often mirror the “Masked Language Model” ^74, 75^, a prevalent training paradigm in natural language processing (NLP). In this paradigm, the expressions of some genes are randomly masked during training, and the model is encouraged to predict the masked values by “referring” to other genes’ expressions. These methods overlook the fact that genes operate in groups as gene modules and assume uniform importance across all genes. Consequently, despite being a good attempt, these methods can possibly be further improved by incorporating our biological modeling principles. In addition, there’s another method for cell-type annotation called TOSICA ^76^ which also incorporates the idea of interactions of gene modules (the term “gene-set” is used in their work, instead). Nevertheless, TOSICA uses the predefined gene-set from pathway as exerting knowledge, and this may restrict the exploration of subtle and dynamic changes in biological activities. Conversely, our intent is to develop an explainer based on STEP to better interpret the biological activities captured by STEP, such as gene regulation networks and pathways. Moreover, hyperparameter tuning also poses challenges, a native problem for deep learning models that may comparatively hinders STEP to reach the potentially highest usability and optimal performance. Going forward, we plan to employ strategies like model distilling to simplify STEP’s training or tuning processes and increase its consistency simultaneously. Our future focus is to address these limitations and refine the model in upcoming works.

Additionally, while most of our experiments focus on 10x Chromium, 10x Visium, MERFISH, STARmap protocols, exploring STEP’s applicability on other datasets can contribute to its further refinement. The potential integration of STEP with other analytical methodologies such as Slide-seq V2 and Stereo-seq, might also lead to unforeseen synergistic effects, further enriching our understanding of spatial transcriptomics and single-cell RNA sequencing. Taking note of recent works around the development of more complex and significantly larger spatially resolved single-cell atlases ^77-80^, we aim to apply STEP/STEP-anchored on such datasets, and such possibilities open exciting avenues for the future research.

In essence, our work forms an initial yet crucial step towards the identification of different types of biological heterogeneity, while integrating multiple batches and sections. Besides, our work demonstrates a competitive integrative analysis across two modalities and underscores the need for stronger analytical strategies and tools in the field.

## Methods

### The STEP backbone model (BBM)

For a batch of gene expression 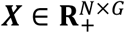 with *N* observations and *G* sequenced genes, the *n*-th gene expression data **x**_*n*_ ∈ ***X*** is firstly performed by a layer normalization. Then, it is mapped to a length *K* sequence corresponding to the gene module ***M***_*n*_ ∈ **R**^*K*×*d*^ by a series of randomly initialized linear transformations **W** ∈ **R**^*G*×*d*×*K*^, where *d* ≪ *G* and each transformation aims to learn one gene module. Also, **x**_*n*_ is inputted into a ReLU function after layer normalization,

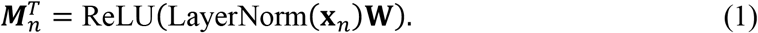

Then, a cls-token ***z***^*cls*^ ∈ **R**^*D*^ is concatenated to ***M***_*n*_ for extraction of the biological information behind **x**_*n*_ and the final low-dimensional representation of **x**_*n*_ is referred to. By denoting 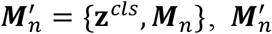, is up-sampled (expanded) to 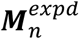 with dimension *D* through another affine transformation. Up-sampling is purposed to counteract any potential negative regulation within each gene module. Since the ReLU function outputs positive values only, each element within each gene module is positive. Also, the attention scores are exclusively positive due to the properties of the softmax function. Next, 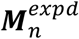 is fed with an additive positional embedding *E*_*pos*_ ∈ *R*^(1+*K*)×*d*^ to *L* Transformer encoders 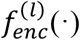, *l* = 1,2,…,*L* (default *L* = 3)

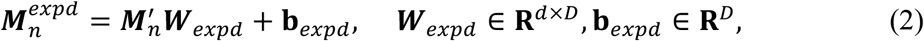

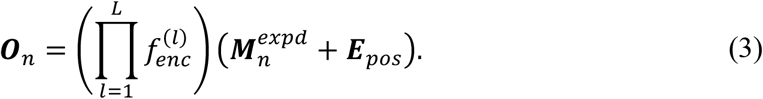

Here, the positional embedding ***E***_*ops*_ is used to differentiate each gene module. Each Transformer encoder 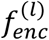 consists of a multi-head self attention module and a feed forward network. The multi-head attention part can be described as

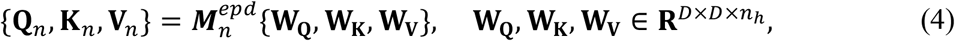

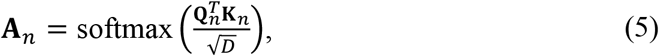

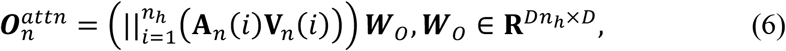

where *n*_*h*_ represents the number of head in multi-head self-attention, 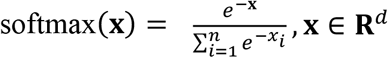 and 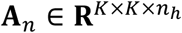 refers to the attention score matrix for each attention head. The products of **A**_*n*_(*i*) and **V**_*n*_(*i*) are then concatenated in head-wise and multiplied by another weight ***W***_**o**_ to obtain the final output 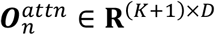 of a multi-head attention module. Also, the feedforward network consisting of two fully connected layers and a non-linear activation function is performed on 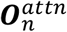

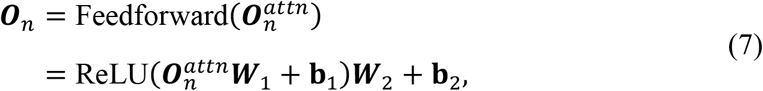

where ***W***_1_ ∈ **R**^*D*×4*D*^, ***W***_2_ ∈ **R**^4*D*×*D*^. The first column of **o**_*n*_ corresponding to the previous cls-token ***z***^*cls*^ is extracted as the non-standardized embedding, and it is denoted as **h**_*n*_.

Afterwards, based on the reparameterization trick used in all VAE models, the **h**_*n*_ is processed by a Readout module *f*_*rout*_ (*⋅*) to estimate the mean ***µ***_*n*_ and the logarithm of variance 2 log ***σ***_***n***_ of the standardized low-dimensional embedding 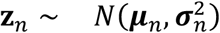.

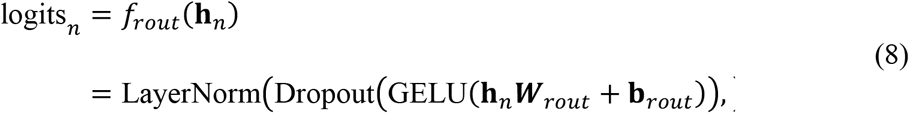

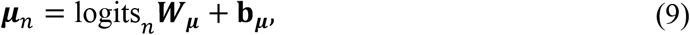

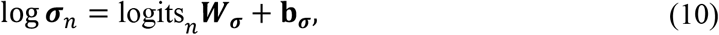

where ***W***_*rout*_ ∈ **R**^*D*×*D*^, ***W***_***µ***_ ∈ **R**^*D*×*d*^. ***z***_*n*_ is inputted into a non-linear decoder *f*_*deC*= 0_(·) to obtain the parameter set of the dispersion, which is shared across cells/spots, and the zero-inflation (**r, π**_*n*_) of the ZINB distribution, as well as the mean expression 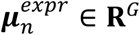,

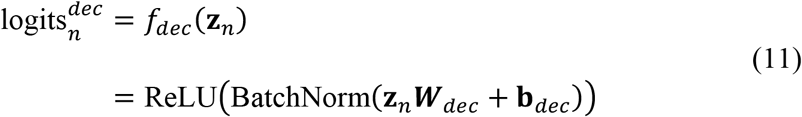

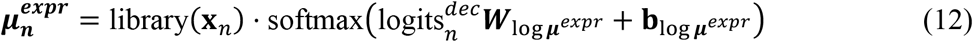

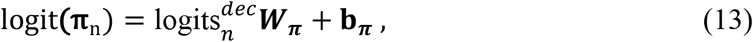

where 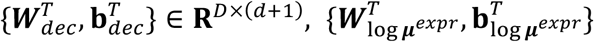 and 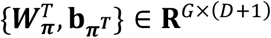 and 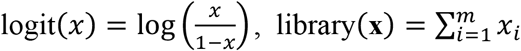 when **x** ∈ **R**^*m*^. The dispersion **r** is randomly initialized and learned during the process of training. After training, the reconstruction/imputation 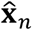 can be sampled for the gene expression data **x**_*n*_ from ZINB(***µ***_*n*_, **r, π**_*n*_). The probability density function (p.d.f.) of 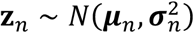 is de-noted as *q*_*ϕ*_ (***z***_*n*_), where *ϕ* refers to the parameters to learn. The p.d.f. of the standard normal distribution, with the dimension equal to that of ***z***_*n*_, is denoted as *p*(***z***_*n*_). The backbone model is trained by minimizing the objective function as follows:

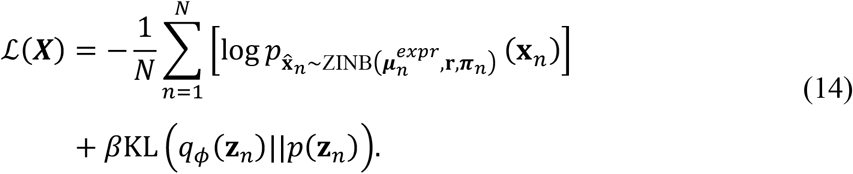

The BBM is then extended with another two models to perform more complex analysis and the combined model is trained by different objectives for different tasks.

### Batch effect model (BEM)

We propose a novel and simple batch-wise layer normalization named BatchAware LayerNorm and use it to replace the LayerNorm before gene module transformations (in Eq.1) when it comes to multi-batches data analysis. Firstly, the one-hot encoding of each batch indicator *b* ∈ *B* is performed and a batch-embedding ***E***_*B*_ ∈ **R**^|*B*|×*D*^ is randomly initialized. Thus, our model can be guided to produce a batch-invariant low-dimensional representation ***z***_*n*_. The computation of BatchAwareLayerNorm can be described as:

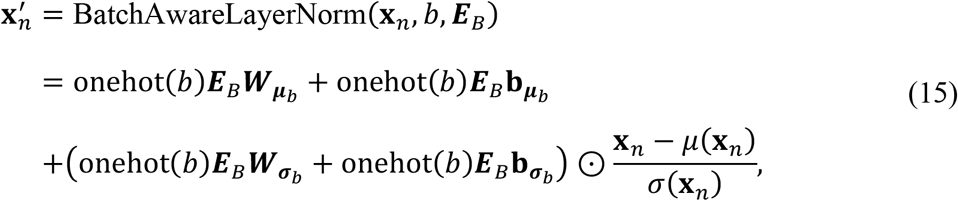

Where 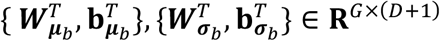. To maintain the consistency in the decoding part, BatchAwareScale is performed on ***z***_*n*_ to retrieve the batch variations before it is sent to decoder *f*_*deC*_ (·):

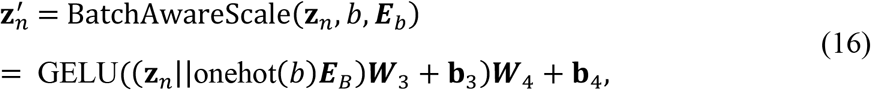

Where 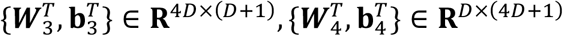. These two paired operations form the BEM. The combination of BBM and BEM is still trained with the basic objective (Eq.14).

If the cell-type annotations set *C* can be accessed, they can be used as labels to train a cell-type classifier *f*_*clsf*_ (·) for a much more precise integration and generation of the centroids for all cell types. Specifically, after the model is trained without annotation information, the set of centroids ***Z***^*anChor*^ ∈ **R**^|*C*|×*d*^ is randomly initialized and the cell-type classifier *f*_*clsf*_ (·) is introduced after the Readout module *f*_*rout*_ (·). For a mini-batch of embeddings 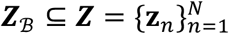, the predicted probability matrices of cell types 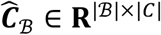 and 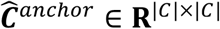 for mini-batches data and ***Z***^*anChor*^, respectively, can be inferred collectively as follows,

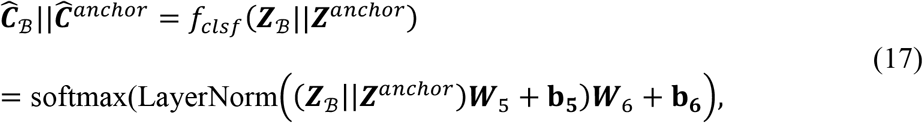

where 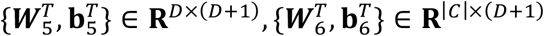.

To train the cell-type classifier while refining the embedding of the BBM, the learning rate of the modules except *f*_*clsf*_ is set to a very low level (10^−5^) and another objective ℒ_*clsf*_ is minimized:

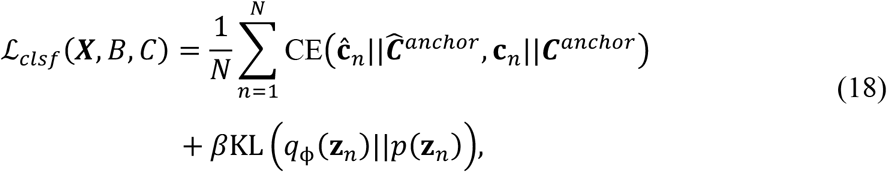

where CE represents cross-entropy, ***c***_*n*_ and ***C***^*anchor*^ represent the annotations of the *n*-th cell and the anchor points, respectively.

### Spatial model (SpM)

To understand the spatial context in tissues, the use of STEP is proposed to generate spatially informed embeddings with the assistance of SpM, which is denoted as *g*(·). STEP starts by constructing the spatial neighboring graphs for each tissue. The SRT data of one section is denoted as ***Y***. For 10x Visium data, a spatial neighboring graph ***A***_***Y***_ is constructed using the hexagonal grids aligned in each tissue. For single-cell resolution SRT data, we compute kNN graph(s) based on the spatial locations in each tissue. ***A***_***Y***_ has binary edges and include self-loops. Next, a vanilla GCN (without residual connection and activation) is applied as a spatial low-pass filter for non-single-cell resolution SRT data, or a smooth operator like a parametric graph Laplacian for single-cell resolution SRT data. SpM filters or smooths the non-standardized embed-ding 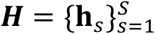 in (Eq.3) before passing it to the Readout module *f*_*rount*_ (·). The output of SpM is denoted as 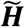. SpM uses *L* (default=4) graph convolutional layers for computation,

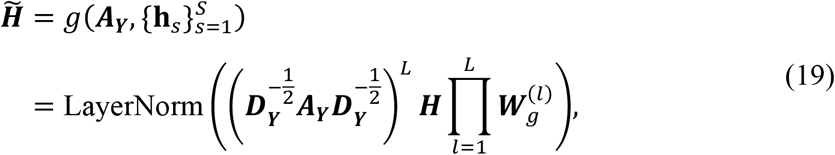

where 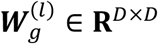 is the weight of the *l* − *th* graph convolutional layer, *l* ∈ {1,2,…,*L*}.

As mentioned earlier, the “fast mode” is adopted to ensure proper running time and direct access to spatial domains for non-single-cell resolution SRT data. It has an architecture like the SimCSE for end-to-end training with the contrastive loss incorporated. The forward process in the BBM of non-standardized embedding is branched, with one path leading to the readout and decoder, and the other path being filtered/smoothed by SpM before joining the same path as the former. In this fast mode, STEP is trained by minimizing the losses calculated through these two forward paths and the contrastive loss of the filtered path and the other path. The contrastive loss servers as a regularization term to restrict the strength of spatial filtering or smoothing. Formally, for each **y**_2_ ∈ *Y*, denoting the non-filtered branch as 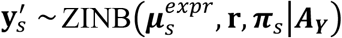 and the filtered branch as 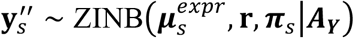, we have the objective ℒ_*st*_ as follow:

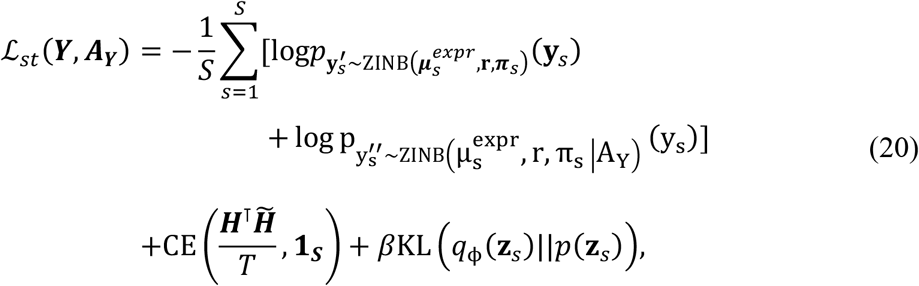

where *T* represents the temperature parameter, and **1**_***S***_ ∈ ***R***^***S***^ is a vector of which the elements are all 1. Also, the SAINT node sampler ^81^ is used to further boost the training speed where the sampler randomly samples several nodes to form a sub-graph in each iteration.

### Revealing targeted biological heterogeneities

STEP effectively balances major variations in the SRT and scRNA-seq data and condenses them into the low-dimensional embedding: ***H*** (non-standardized), ***Z*** (standardized), and 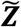 (spatially smoothed and standardized). With these embeddings, it is easy to reveal the targeted biological heterogeneities, cell types (states) and spatial domains through commonly used community detection or clustering algorithms. This allows users to run STEP once to try multiple domain numbers, just like the standard scRNA-seq data analysis workflow. For the access of sub-cell-types and sub-spatial-domains, users can achieve this purpose by simply increasing the number of clusters to detect or run clustering method in a hierarchical manner.

A special case is the integrative analysis for a pair of annotated scRNA-seq datasets and non-single-cell resolution SRT datasets. Herein, a cell type deconvolution algorithm reliant on their co-embedding is proposed. Particularly, in the deconvolution process of STEP, we denote scRNA-seq dataset as 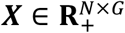 and SRT dataset as 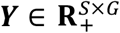. The co-embedding result (using non-standardized embedding) is denoted as ***H***^*T*^ = {(***H***^*SC*^)^*T*^; (***H***^*st*^)^*T*^} ∈ **R**^*D*×(*N*+*S*)^. The spatial domains in the tissues are assumed as identified to estimate the cell-type composition 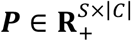 for the SRT dataset, where *C* represents the cell type annotation set. Therefore, the first step is to generate cell-type anchors ***H***^*anchor*^ ∈ **R**^|*C*|×*D*^ by averaging the non-standardized embedding 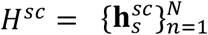 based on cell types. Then, the cell-type signatures 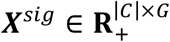 are com-puted by normalizing and averaging the gene expressions in reference scRNA-seq dataset to a probability matrix based on cell types. For each spot (*s*-t*h* spot), STEP concatenates each 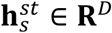 and cell-type-anchors ***H***^*anchor*^, and feeds them into the new Transformer encoders 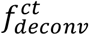. The output is split to 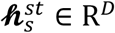 and 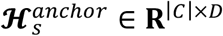 where 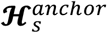 is considered as the spot-specific cell-type-anchors and it is constrained to be similar with 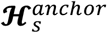 as possible. They are further fed as query and key into a sparse attention module *f*_*attn*_ to calculate the non-negative cell-type similarity matrix 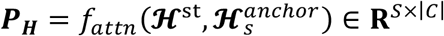 in the latent space where 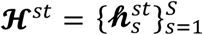 and the “convolution” of scRNA-seq data embedding in the latent space is represented by 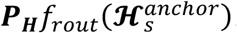. This convoluted result is then sent to the decoder to compute the reconstruction of data. As its name suggests, each entry of ***P***_***H***_ represents the similarity between some cell types and some spots.

The estimation 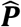 of cell-type composition ***P*** at gene-expression level is transformed from the similarity matrix 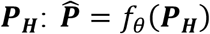 by some transformation *f*_θ_. Since the embedding is sufficiently informative and the subtle difference will be complemented by minimizing the reconstruction error term, the transformation *f*_θ_ is not expected to change ***P***_***H***_ significantly. Also, according to our assumption, ***P***_***H***_ is defined as the cell-type similarity matrix. Therefore, it is supposed to recognize what cell types are included in the spot. Thus, the transformation *f*_θ_ is modeled as an element-wise scale function: 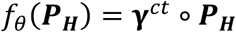, where ***γ***^*ct*^ models the cell-type-wise scale.

The gene expression data is modeled by ZINB(***µ*, r, π**), where parameters (***µ*, r, π**) refer to mean, dispersion and zero-inflation rate, respectively. The zero-inflation parameter is modeled as a cell-dependent or spot-dependent parameter. We additionally model a “convolution” at the gene expression level: 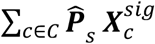, where 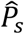 is the *s*-th row of 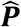 and 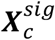 is the *c*-th column of ***X***^*sig*^. For the *s*-th spot **y**_*s*_, the NB part is directly parameterized by the shared dispersion **r** and two mean modelling ways as 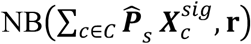 and 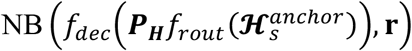, while the zero-inflated rate **π**_2_ is inferred from the embedding 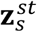 of this spot. Thus, the objective ℒ_*’deConv*_ of deconvolution process can be written as:

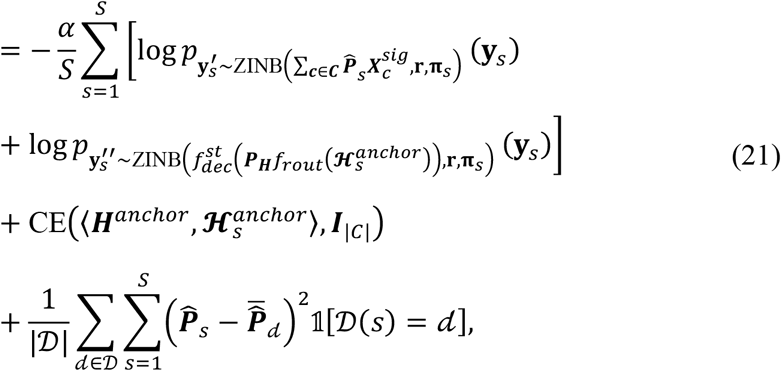

where *α* is a hyper-parameter (default 0.1), 𝒟 represents the set of identified spatial domains, 𝒟(*s*) represents the spatial domain to which spot *s* belongs, ***I***_|*C*|_ ∈ **R**^|*C*|×| *C* |^ represents the identity matrix, 𝟙[*⋅*] represents the indicator function, and 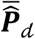 represents the average estimated cell-type composition of spots in spatial domain *d*.

### Hyper-parameter settings

We use Adam as our optimizer and set the learning rate to 0.001, and the validation rate is set to 0.2. The number of training epochs is set to 400 and Early Stopping is enabled by default. The number of gene modules *K* is set to 32 by default, and this value is tested on almost all datasets studied in this paper, showing a strong robustness. However, a high value of *K* means a longer running time, and this value can be reduced to 10-12 on some simple datasets (like Human DLPFC dataset) with only a few cell types or spatial domains. Similarly, the dimensions of ***H*** and ***Z*** are set to 64 and 30, respectively. However, they can be reduced to 40 and 20 when the loading and complexity of data are moderate, like Mouse medial prefrontal cortex data by STARmap. Overall, STEP is robust to the setting of these hype-parameters. The critical setting of hyper-parameters relates to the value of *β* (default 10^−2^). In the range from 10^−2^ to 10^−4^, STEP performs consistently well. For the best performance on a certain dataset, however, like attempting to produce the best batch-removal performance, it is necessary to tune the *β* carefully (often increase the *β* value from 10^−1^) since an excessively high value of *β* may lead to overfitting on the bottleneck layer and cause the model to collapse. On the contrary, STEP without the variational approach can also work well and is also good in trajectory conservation but the points cannot cluster too tightly in the view of UMAP. Also, its performance deteriorates in the batch correction score, which might lead to the mis-cluster of cell types, although spatial domains are quite robust to the value of *β*. In conclusion, too high a value of *β* may lead to collapse of the model while the absence of the variational approach STEP may cause trouble in clustering cells in the common ways.

Focusing on SRT data, the number of iterations is set to 2000 for single-batch in the fast mode, and the number of sampled spots/cells is set to 2048. The number of graph convolutional layers in spatial model is set to 4 by default. Since this hyper-parameter works for the filtering or smoothing range in the tissue, different values affect the smoothness and continuity of the identified domains. According to our tests, the performance of STEP is consistent when the number of graph convolutional layers ranges 4 to 6. An important model setting that causes overfitting of the model is whether to use BatchNorm in the decoder. There is no deterioration in performance for STEP without BatchNorm since BatchAwareScale operation plays a similar role to BatchNorm in a categorical batch-wise manner rather than mini-batch-wise manner. Notably, STEP can be trained much faster with much fewer epochs with BatchNorm. This phenomenon occurs widely in the field of deep learning. However, regarding the occurrence of an extremely unbalanced level of counts, or sequencing depths, for the batches in a dataset, the use of BatchNorm is prone to overfitting. This is understandable, since BatchNorm performs mini-batch-wise normalization where the mini-batch in training STEP is comprised of several sampled sections, and such universal normalization is clearly incompatible with a significant variation in the level of counts. Otherwise, when this level is balanced, it means a similar level of expression. As a universal way of normalization, BatchNorm can promote the convergence of STEP.

## Metrics

### Cell-type level Multi-batch Integration Metrics

The whole process of evaluation follows the scib ^48^ pipeline with the same metrics used. The evaluation conducted in scib pipeline involves two main aspects. One relates to the metrics for the Bio conservation, including normalized mutual information (NMI), adjusted rand index (ARI), average silhouette width (ASW) measured by cell-type (ASW cell-type), isolated label scores (F1 and ASW) and graph local inverse Simpson index (LISI) for cell-type. The other relates to the metrics for the Batch correction, including average silhouette width measured by batch (ASW batch), principal component regression (PCR), kBET and iLISI.

Except the results obtained through STEP, all the other results are directly obtained from the original study. Additionally, all the graph-based metrics (in this test) are evaluated in the kNN graph built on the outputted embedding of each method with the same *k* values.

### Normalized Mutual Information (NMI)

NMI evaluates the match level between clustering results and ground truth by examining their mutual information. Then, NMI normalizes it based on their individual entropies, generally resulting in a value between 0 to 1. High NMI means a substantial mutual relationship is present, usually indicating a satisfactory outcome of clustering that matches the ground-truth labels. Formally, for a clustering result ***X*** and the ground truth ***Y***, NMI can be computed as

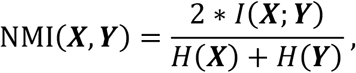

where *I*(*⋅* ; *⋅*) represents the mutual information and *H*(*⋅*) represents the entropy.

### Adjusted Rand Index (ARI)

ARI is a metric for evaluating the correspondence between clustering results and ground truth labels. It is normalized between -1 and 1. A high positive ARI signifies that the clustering result aligns well with the ground-truth labels, with 1 indicating the perfect match up and negative values indicating independent labels. Formally, for a set of *n* elements and two partitions of these elements ***a*** and ***b***. ARI can be computed as

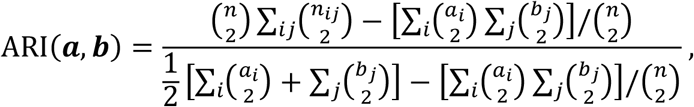

where *a*_*i*_ and *b*_*j*_ are the sum of pairs in the same set within ***a*** and ***b***.

### Silhouette Coefficient

The Silhouette Coefficient is a metric used to evaluate the consistency within clusters in a dataset. For each sample in the data set, the Silhouette Coefficient is evaluated using the mean intra-cluster distance, which refers to the average distance of the sample to all other data points in its assigned cluster, and the mean nearest-cluster distance, which refers to the average distance of the sample to all samples of the next nearest cluster. Defined for each data *i*, it is composed of intra-cluster distance *a*(*i*) and mean nearest-cluster distance *b*(*i*):

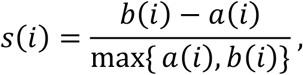

### LISI

LISI is a measure used to evaluate the quality of clustering results in mixed samples. It evaluates the diversity of labels in the local neighborhood of each sample, enabling an assessment of the mix of batches or cell types in a reduced space. For each data *i, C*_*i*_ denotes the cluster or the category that the sample belongs to, and *N*(*i*) represents the data that are neighboring to *i*. The LISI score is individually calculated for each sample *i* through the inverse Simpson concentration of its neighboring samples, which is referred to as LISI(*i*). The overall Local Inverse Simpson Index for the entire dataset is determined by taking the harmonic mean of the LISI(*i*) values across all samples in the dataset. A higher LISI value indicates a better mix of the categories within the dataset,

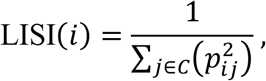

where ***p***_*i,j*_ denotes the proportion of category *j* in the *N*(*i*).

### kBET

kBET ^82^ is a statistical test focusing on the local batch label distribution in the k-nearest neighbor graph and compares it with the global batch label distribution to derive a null distribution. For each data, the batch label distribution of its k-nearest neighbors is compared to the expected batch label distribution. After performing this computation for a subset or all data points, the kBET rejection rate can be calculated, which quantifies the overall batch effects (a low rejection rate implies minor batch effects).

### Principal Component Regression (PCR)

PCR, through PCA, decomposes the original data matrix into principal components (PCs) for assessing the variance of this data as explained by the batch covariates. By the independences between PCs, the variance of data matrix explained by batch covariate is calculated by the sum of the product of the variance of data matrix explained by each *PC*_*i*_ and the variance of each *PC*_*i*_ explained by batch covariates:

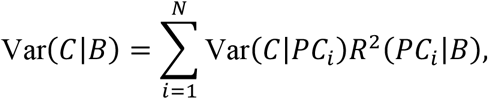

where *C* is the data matrix, and *B* is the batch covariate.

### Isolated Labels

**“**Isolated labels” is a metric for the identification accuracy of cell labels shared in only a few batches. This metric is calculated in two ways: F1 score and ASW score.

### Spatial-domain level Multi-batch Integration Metrics

Identifying spatial domains is a task distinct from cell-type clustering, due to its reliance on the spatial location of data points. In contrast, cell-type clustering focuses more on the inherent properties and similarities among data points, without consideration for geographical context. To evaluate the spatial domains identified, we use two additional metrics: Moran’s I and Geary’s C. By employing these metrics, we can comprehensively evaluate the performance of spatial domain identification in terms of continuity and smoothness.

### Moran’s I

Moran’s I measures the level of spatial autocorrelation and serves as an indicator of spatial pattern divergence from randomness. This assists in understanding the spatial continuity within the identified regions. Given a set of *N* cells/spots with their neigh-boring graph 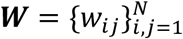 and the corresponding spatial domains 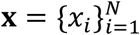 with its mean 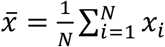, the Moran’s I is calculated as

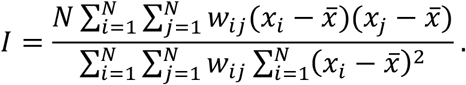

The neighboring graph ***W*** is constructed as kNN graph with *k* values ranging from 1, 10, 20, 30.

### Geary’s C

Geary’s C is applicable to assess spatial heterogeneity by measuring dissimilarity between neighboring locations, which reflects the smoothness of the spatial distribution. Given a set of *N* cells/spots with their neighboring graph 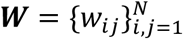 and the corresponding spatial domains 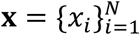, the Geary’s C is calculated as

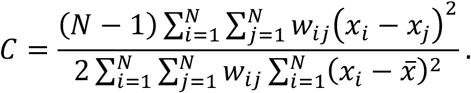

The neighboring graph ***W*** is constructed as kNN graph with the values of *k* ranging from 1, 10, 20, 30.

### Cell-type Deconvolution Metrics

#### Pearson Correlation Coefficient (PCC)

PCC is a measure of the linear correlation between two variables. It’s a value between -1 and 1 where -1 is total negative linear correlation, 0 is no linear correlation, and 1 is total positive linear correlation.

Given two vectors **x** and **y** of *n* samples, the Pearson Correlation Coefficient *ρ* can be calculated with sample means 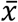 and 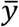 using the following formula:

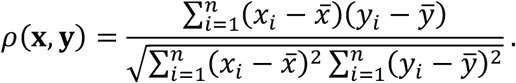

### Spearman Correlation Coefficient (SCC)

SCC measures the strength and direction of monotonic relationship between two variables. This is a non-parametric measure and unlike the Pearson correlation, it doesn’t require the assumption of normality or linearity. The coefficient is in the range of -1 to 1, where -1 indicates a perfect decreasing monotonic relationship, 0 indicates no monotonic relationship, and 1 indicates a perfect increasing monotonic relationship. Let **x** and **y** be the rank orders of two vectors of *n* samples. The Spearman Correlation Coefficient *ρ*_*S*_ can be calculated as:

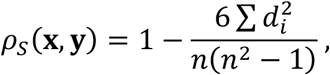

where *d*_*i*_ represents the difference between the ranks of each observation on *x*_*i*_ and *y*_*i*_.

### Jensen–Shannon Divergence (JSD)

JSD is a method to measure the similarity between two probability distributions. Given two probability distributions *P* and *Q*, the Jensen-Shannon Divergence can be calculated as follows:

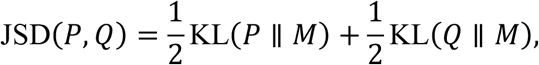

where KL is the Kullback-Leibler divergences and 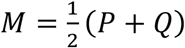.

### Mean Squared Error (MSE)

MSE is a common metric used to evaluate how well a model’s predictions match the actual outcomes, where lower values indicate better performance. The MSE is calculated based on true value vector 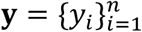 and prediction vector 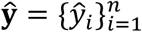 us-ing the following formula:

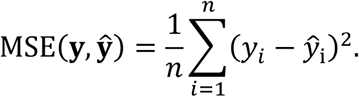

### Synthetic data for validation of deconvolution

The pseudo-SRT data is constructed in the following process: 1. Choose one SRT data and its tissue section and a multi-batches scRNA-seq dataset as reference; 2. One batch of scRNA-seq data is chosen for the simulation of SRT gene-expression and that of technical effect between scRNA-seq and SRT data; 3. The spatial domains denoted as 𝒟 are generated for the tissue of SRT data, with STEP directly used to identify the spatial domains based on the original SRT data, and the mean cell-type composition *E*[***P***_*d*_] of each domain *d*; 4. For each spot, the number of cells, cell-type composition and corresponding gene expression are sampled from uniform distribution, normal distribution and scRNA-seq dataset, respectively. 5. A spatially local smooth is performed on cell-type composition {***P***_*d*_}_*d*∈𝒟_ to simulate the spatially correlation with 2 graph convolutions; 6. The sampled gene expressions and cell-type composition are mixed as gene profiles for the spots.

### Analyzed Datasets and Analysis Details

#### Human Colorectal Cancer and Mouse Small Intestine by 10x Visium HD

We downloaded Human Colorectal Cancer and Mouse Small Intestine data from 10x public datasets (https://www.10xgenomics.com/resources/datasets). Human Colorectal Cancer data with a bin size of 16µm was used to perform test. Mouse Small Intestine data with a bin size of 8µm was tested. STEP was applied to analyze these two datasets under the default parameters setting.

#### Integration of Atlas-level scRNA-seq datasets

We downloaded the 6 benchmarking scRNA-seq datasets from GitHub repository of scib-reproducibility (https://github.com/theislab/scib-reproducibility), including 2 simulation datasets (Simulation1, Simulation2) and 4 real datasets (Human Pancreas, Human Lung, Human Immune and Human & Mouse Immune). The Simulation1 dataset contains 12097 cells and 6 batches. Simulation2 dataset contains 19318 cells and 16 batches. Human Pancreas dataset contains 16382 cells and 9 batches. Human Lung dataset contains 32472 cells and 16 donors. Human Immune dataset contains 33056 cells and 10 donors. Human & Mouse Immune dataset contains 97952 and 23 samples. All the results of compared methods were also obtained from scib-reproducibility. STEP was tested with 2000 HVGs by referring to the scib-pipeline and the same parameters-setting across all datasets: 32 modules; hidden dimensions *D* = 64, *d* = 30; *β* = 0.01. Also, it was trained in 400 maximum epochs with the batch size set to 1024. All the results of STEP are presented in the Supplementary Figures and Tables.

### Human DLFPC dataset by 10x Visium

We obtained 12 space ranger outputs of DLFPC dataset from GitHub repository spatialLIBD (https://github.com/LieberInstitute/HumanPilot) as well as the manual annotations.

For the spatial domain identification of a single section, STEP was tested with the parameters listed in (Supplementary Tables 2). The compared methods were all tested according to the original studies. The complete results are presented in the Supplementary Figures.

For the integration of multiple sections, we set the parameters of STEP as follows: top-3000 HVGs,16 modules, hidden dimensions *D* = 64, *d* = 30, *β* = 0.001, 4 graph convolutional layers. Also, the STEP was trained in 1000 iterations, with the graph sample size set to 1. The compared method BASS-Mult was tested by following the official workflow. The other method PRECAST was tested in the default settings. Other compared methods were also tested by following their related official documentations. The complete results are presented on Supplementary Figures and Tables.

#### Mouse Sagittal Anterior and Posterior by 10x Visium

These two sections were obtained from 10x Visium public dataset (https://www.10xgenomics.com/resources/datasets). We set the parameters of STEP as follow: top-3000 HVGs,16 modules, hidden dimensions *D* = 40, *d* = 30, *β* = 0.01, 3 graph convolutional layers. STEP was trained in 800 iterations with the graph sample size set to 2.

#### Mouse Hypothalamus data by MERFISH

We downloaded this data from Dryad (https://datadryad.org/stash/dataset/doi:10.5061/dryad.8t8s248). The test was conducted for the cell-type level integration and domain level integration in the 5 sections (Bregma distance -0.04 to -0.24) from the first animal. The manual annotations for functional regions were obtained from (Li & Zhou, 2022), and all genes were used to perform tests. For the cell-type level integration test, STEP was set as 20 modules, hidden dimensions *D* = 40, *d* = 30, *β* = 0.01 and was trained in 400 epochs. The compared methods were all tested using default parameters. For the test on domain-level integration, STEP was set with additional 4 graph convolutional layers and was trained in 1600 iterations with the graph sample size set to 2. In this test, BASS-Mult was tested according to the official tutorial and PRECAST was tested with 30 nearest neighbors and all genes.

Furthermore, the additional integration of 12 sections (Bregma distance from 0.26 to -0.29) from the 7-th animal in the original study was tested by STEP. The complete results are presented in Supplementary Figures.

#### Mouse Medial Prefrontal Cortex data by STARmap

We downloaded this dataset from STARmap public dataset (https://www.Starmapresources.org/data). The test was conducted on 3 sections: BZ5, BZ9, BZ14, with manual annotations obtained from (Li & Zhou, 2022). For the simplicity of the data, STEP was set to 8 modules, hidden dimensions *D* = 40, *d* = 30, *β* = 0.01 and was trained in 1600 iterations, with the graph sample size set to 2. BASS-Mult was tested by following the official workflow and PRECAST was tested with all genes and 30 nearest neighbors.

#### Human Liver Normal and BA data by 10x Visium

Four of the SRT data were downloaded from (https://github.com/julietusc/Transcriptomics_technical_optimization) and the other two were downloaded from (https://doi.org/10.6084/m9.figshare.22321447.v1). The reference snRNA-seq data was also obtained from (https://github.com/julietusc/Transcriptomics_technical_optimization). For the spatial domain identification, we selected top-2000 HVGs and set STEP with 20 modules, hidden dimensions *D* = 64, *d* = 30; *β* = 0.01. STEP was set to run 2200 iterations with 0.7 node sampling rate in fast mode. For the co-embedding stage, we used top-2000 HVGs and set STEP with 32 modules, hidden dimensions *D* = 64, *d* = 30; *β* = 0.01. In this stage, STEP was trained in 400 maximum epochs, with the batch-size set to 1024. The spatial domains were identified alone without using co-embedding and ST-decoder. In the deconvolution stage, STEP was set to run 1500 iterations based on the identified spatial domains. All the results are presented in the Supplementary Figures.

## Data availability

All the relevant data supporting the key findings of this study are available within the article. The datasets used for the analysis in this study are available from the following sources: (1) Human Colorectal Cancer and Mouse Small Intestine data from 10x Visium HD at https://www.10xgenomics.com/resources/datasets;(2) six benchmarking scRNA-seq datasets from scib-reproducibility GitHub repository at https://github.com/theislab/scib-reproducibility, including Simulation1, Simulation2, Human Pancreas, Human Lung, Human Immune, and Human & Mouse Immune datasets; (3) 12 space ranger outputs of DLFPC dataset from spatial-LIBD GitHub repository at https://github.com/LieberInstitute/HumanPilot; (4) Mouse Sagittal Anterior and Posterior sections data from 10x Visium public dataset at https://www.10xgenomics.com/resources/datasets; (5) Mouse Hypothalamus data by MERFISH from Dryad at https://datadryad.org/stash/dataset/doi:10.5061/dryad.8t8s248; (6) Mouse medial prefrontal cortex data by STARmap at https://www.starmapresources.org/data; and (7) Human Normal and BA liver data by 10x Visium and snRNA-seq data by 10x at https://github.com/julietusc/Transcriptomics_technical_optimization; (8) Human Normal liver data by 10x Visium at https://doi.org/10.6084/m9.figshare.22321447.v1.

## Code availability

The open-source Python package of STEP is available at GitHub repository: https://github.com/SGGb0nd/step. The tutorials and documentations are being hosted at https://sggb0nd.github.io/step.

## Author contributions

Z.L. and X.X. designed the study. L.L. designed the model and algorithm. L.L. developed STEP software and wrote the related documentations. L.L. and X.X. wrote the manuscript. L.L. did all visualization work. L.L. and X.X. performed data analyses. X.Y. performed the mouse liver cell type and domain annotations, as well as the interpretation for the results of the analyses. All authors read and approved the manuscript.

## Founding

This work is supported by the National Natural Science Foundation of China under Grant No. 12171434 and Science and Technology Plan Project of Huzhou City, China (no. 2022GZ51).

## Notes

### Competing Interest Statement

The authors have declared no competing interest.

### Summary of Updates

Replace the case-study of mouse liver dataset with the one of human liver dataset

