## Supplementary Information for "STEP: Spatial Transcriptomics Embedding Procedure for Multi-scale Biological Heterogeneities Revelation in Multiple Samples"

### Contents

**1 Supplementary Information** **2**

**2 Supplementary Notes** **2**

2.1 Additional studies . . . . . . . . . . . . . . . . . . . . . . . . . . . . . . . . . . . . . 2

2.1.1 STEP eliminates the batch-effect and corrects gene-expression of scRNA-seq 2

2.1.2 Exploring the Cellular Characteristics of Human Lymph Nodes using STEP. 2

2.1.3 STEP enables scRNA-seq reference mapping in PDAC data by ST . . . . 3

**3 Supplementary Figures** **4**

**4 Supplementary Tables** **129**

**References** **137**

### 1 Supplementary Information

#### STEP: Spatial Transcriptomics Embedding Procedure for Multi-scale Biological Heterogeneities Revelation in Multiple Samples

### 2 Supplementary Notes

#### 2.1 Additional studies

##### 2.1.1 STEP eliminates the batch-effect and corrects gene-expression of scRNA-seq

The decoupling of batch-effect model and gene-expression backbone model constructs some kind of map from batch id, actually batch-embedding, to the batch-effect, enabling a direct control of batch-effect. As a result, seeing one batch as one kind of style in the area of image generation, we can transfer all count data from their original batch (style) to a given batch (style) by giving the same batch id to the batch-decoder in the decoding process, just like any conditional generative model (GAN, VAE and diffusion model).

In this way, all gene-expression data are transferred into a same feature space. More robust way can be using the average or weighted average of batch-embedding since some batch-effects represent a serve serious damage on the biological signals. We benchmarked this gene-expression-correction method in the same way with the integration of multi-batch scRNA-seq dataset (Luecken et al., 2022) by performing the PCA on the corrected gene-expression data. The benchmarking results demonstrate a significant improvement for the PCs from corrected gene-expression data compared to the PCs from the raw gene-expression data (Supplementary Fig. 37-42) in the sense of biological conservation.

Despite of the advantageous results, the limitation of this method is also obvious: only the genes used for the model training, HVGs in most case, can be corrected so that the further usage for gene-expression related downstream analysis will be hindered, like differential expressed gene analysis. Same issue was also encountered by SCTransform(Hafemeister & Satija, 2019), and such limitation need further research to address.

##### 2.1.2 Exploring the Cellular Characteristics of Human Lymph Nodes using STEP.

To further demonstrate the adaptability of STEP, we employ it to dissect the complex tissue microenvironments within human lymph nodes. Unlike structured organs such as the brain, human lymph nodes encompass dynamically intertwined cellular ecosystems. Leveraging a publicly available SRT dataset of human lymph nodes from 10x Genomics, we utilize STEP to extract spatial clusters from a reference scRNA-seq data composed of 34 sub cell types and 73,260 cells derived from the analysis in original study (James et al., 2020; King et al., 2021; Park et al., 2020) (Supplementary Fig. 43a).

We perform the STEP-featured integrative analysis on SC-data and SRT-data. First, we obtain the remarkable co-embedding result (Supplementary Fig. 43b) that SC-data still preserves the cell-type structure even as it is aligning with SRT-data and SRT itself is surrounded by SC-data and saturates into the populations of related cell-types. Such co-embedding result demonstrates the capacity of STEP to handle such cross-modality integration task.

Anatomical assessment of the lymph node sample reveals several distinctive features, including numerous germinal centers (GCs). After acquiring STEP's co-embedding results, we leverage the embeddings to uncover spatial domains and to reveal the cellular composition in each spot and each domain by performing the cell-type deconvolution, allowing us to predict potential cellular heterogeneities and gain deeper insights into tissue organization. As a result, total 8 domains and 24 sub-domains are identified (Supplementary Fig. 43c, d) and the germinal centers (GCs) are successfully recognized as the domain 8 referring to the histology image. The cell-type-specific domains, like T cell region (domain 4) and Plasma region (domain 5), are revealed and verified by the cell-type composition inferred by STEP (Supplementary Fig. 43e). Interestingly, STEP finds the follicle region (domain 1) as a boundary-like spatial domain, surrounding the spots in GCs region and separating this region and T-cell-specific region. The cell-types mapped into this domain also confirm this assertion as the non-matured B cells (B naive, B activated, etc.) are all enriched in this domain (Supplementary Fig. 46e).

To further study this follicle region for the potential dynamical development insights including proliferation and differentiation, we additionally select 2 spatially adjacent and cellular composition related domains: GCs (domain 8) and T cell region (domain 4), to perform a combined analysis on both levels of the gene expression profile and cell-type composition. The Spatially DE genes analysis discerns the much more similarity between follicle region and GCs region and the similarity between follicle region and T cell region in the gene expression profile, suggesting the GCs and follicle region are distinct to T cell region in the view of tissue compartments (Supplementary Fig. 43f). By the examination of scaled cellular composition, it remarks that the differentiation of B cells in the follicle regions as the compositions of B activated and B "preGC" are significantly higher than GC region (Supplementary Fig. 43g). This discrimination facilitates the identification of regions/domains harboring concentrated populations of specific cell types.

Aligning with these visual observations, STEP generates comprehensive maps pinpointing key regions/domains within the lymph node tissue (Supplementary Fig. 44, 45), notably highlighting structures and cell types such as aforementioned T cell region, GCs and follicle region. These collective findings strongly indicate that STEP serves as a potent tool for mapping complex tissues marked by interconnected cell types spatially. This capability significantly contributes to biological interpretation by jointly uncovering spatial domains and cellular compositions.

###### 2.1.3 STEP enables scRNA-seq reference mapping in PDAC data by ST

It is known that one of the biggest issues of some spatial resolved transcriptomics data, like 10x Visium and Spatial Transcriptomics (ST) is the non-single-cell resolution of the sequencing units. Though STEP has shown its flexibility to reveal cell-type/cell-state level by cell-type/cell-state deconvolution combined with domain identification, there is another solution for this issue that the scRNA-seq data will be mapped into the appropriate spatial locations of the tissue. With the high-quality of the co-embedding of scRNA-seq and SRT data produced by STEP, STEP also has potential to achieve this task. Specifically, for each location in the tissue, we simply find its k nearest neighbors from scRNA-seq reference data by evaluating the similarities between embedding of these two modalities. In spite of the simplicity, the mapping results are quite good thanks to the STEP's high-quality co-embedding as we tested the method in the series of pancreatic ductal adenocarcinomas (PDAC) data by ST(Moncada et al., 2020), a kind of SRT data with relatively lowest resolution.

Moreover, with the batch-effect correction ability of STEP, the scRNA-seq can be simultaneously and uniformly mapped to multiple ST samples (Supplementary Fig. S46). We evaluate the mapping quality by visualizing and comparing the expression pattern of marker genes between mapped and raw data (Supplementary Fig. 47-52). All results show the high-consistency of the expression pattern in both cases of enrichment and depletion. In addition, we separately perform spatial domain identification and cell-type deconvolution on PDAC-A ST dataset and PDAC-B dataset to inspect the scalability of STEP to the low resolution SRT data, and obtain the robust results compared to the original study (Supplementary Fig. 53-62).

#### 3 Supplementary Figures

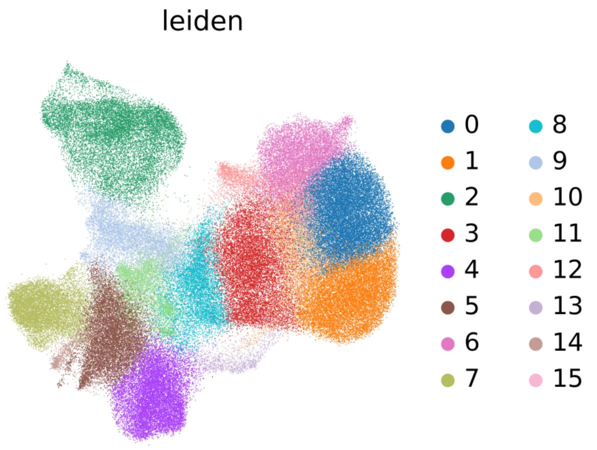
leiden

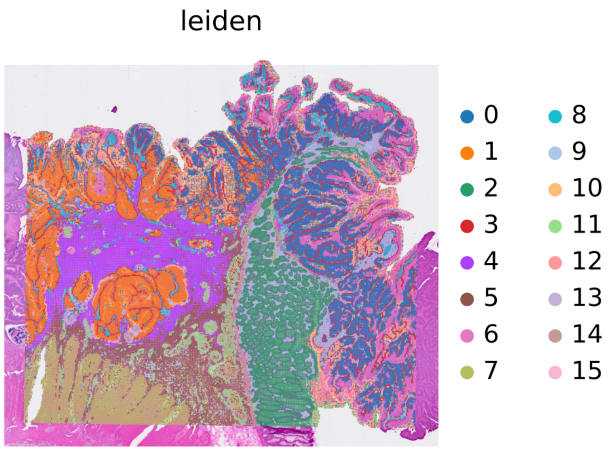
leiden

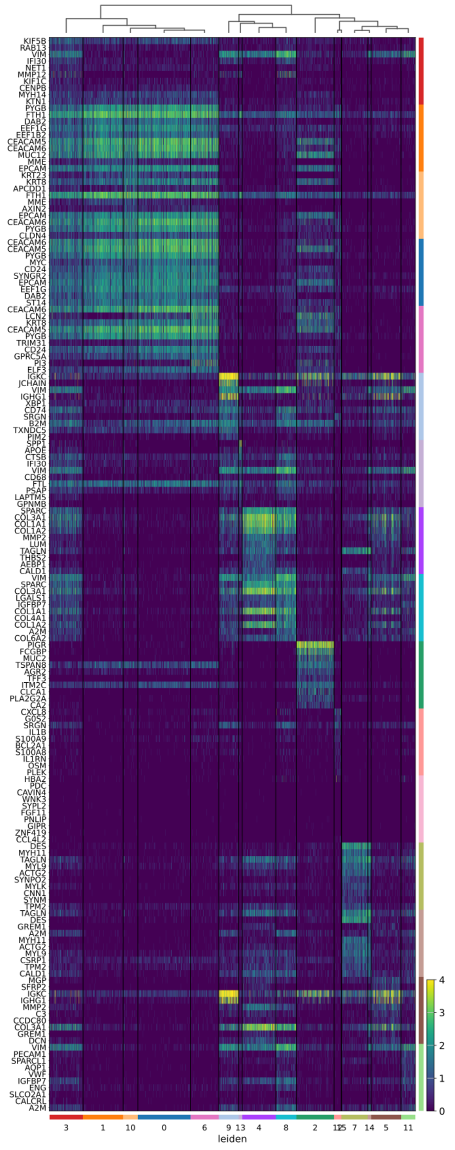
leiden

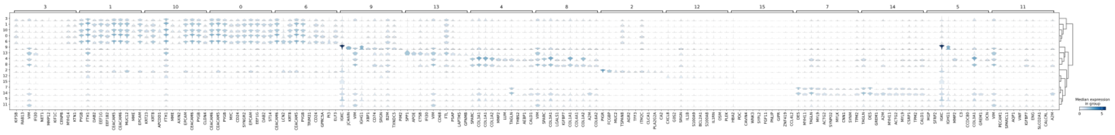

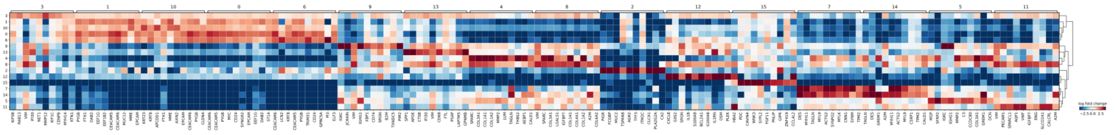

**Fig. S1 Results of basic analyses including clustering and marker genes of Human Colorectal Cancer by Visium HD.** The first column shows the clustering results on UMAP and tissue section, respectively. The second column shows the heatmap of expressions of marker genes in each cluster. The last 2 rows show the stacked violin plot of the expressions of marker genes, and the matrix plot of log-foldchange values of each marker gene in each cluster.

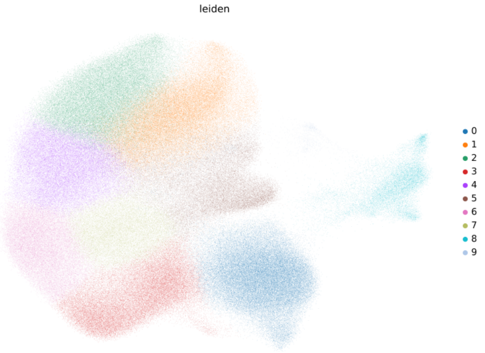
leiden
0
1
2
3
4
5
6
7
8
9

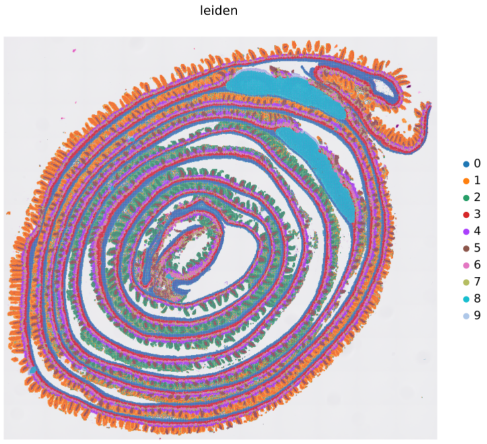
leiden
0
1
2
3
4
5
6
7
8
9

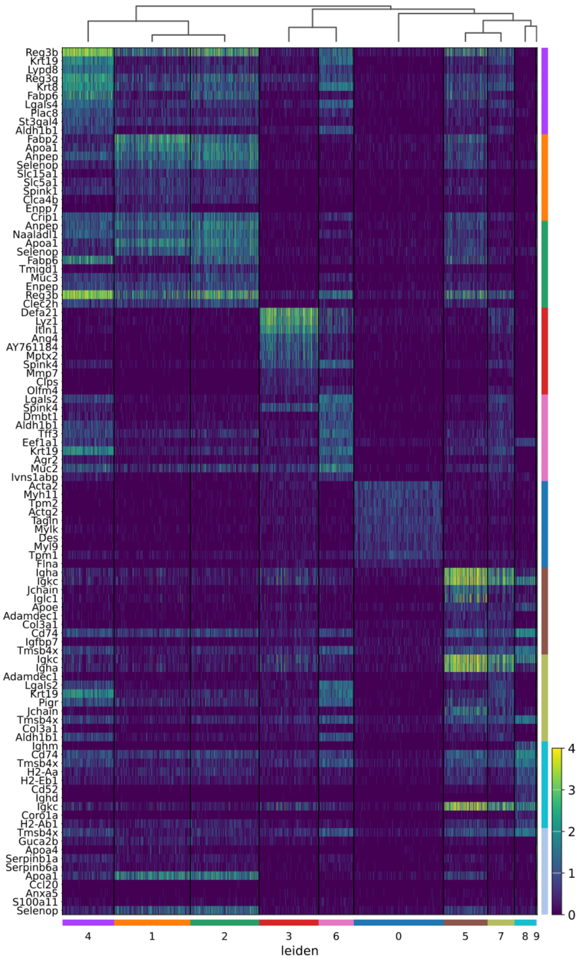
Reg3b
Krt19
Lypd8
Reg3g
Krt8
Fabp6
Lgals4
Plac8
St3gal4
Aldh1b1
Fabp2
Apoa1
Anpep
Selenop
Slc15a1
Slc5a1
Spink1
Clca4b
Enpp7
Crip1
Anpep
Naaladl1
Apoa1
Selenop
Fabp6
Tmigd1
Muc3
Enpep
Reg3b
Clec2h
Defa21
Lyz1
Itln1
Ang4
AY761184
Mptx2
Spink4
Mmp7
Clps
Olfm4
Lgals2
Spink4
Dmbt1
Aldh1b1
Tff3
Eef1a1
Krt19
Agr2
Muc2
Ivns1abp
Acta2
Myh11
Tpm2
Actg2
Tagln
Mylk
Des
Myl9
Tpm1
Flna
Igha
Igkc
Jchain
Iglc1
Apoe
Adamdec1
Col3a1
Cd74
Igfbp7
Tmsb4x
Igkc
Igha
Adamdec1
Lgals2
Krt19
Pigr
Jchain
Tmsb4x
Col3a1
Aldh1b1
Ighm
Cd74
Tmsb4x
H2-Aa
H2-Eb1
Cd52
Ighd
Igkc
Coro1a
H2-Ab1
Tmsb4x
Guca2b
Apoa4
Serpinb1a
Serpinb6a
Apoa1
Ccl20
Anxa5
S100a11
Selenop
4
3
2
1
0
4
1
2
3
6
0
5
7
8
9
leiden

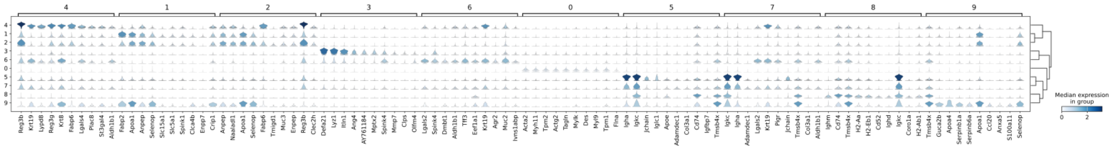
4
1
2
3
6
0
5
7
8
9
Median expression in group

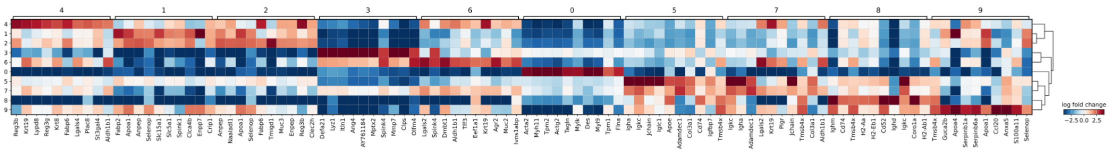
4
1
2
3
6
0
5
7
8
9
log fold change
−2.5 0.0 2.5

**Fig. S2 Results of basic analyses including clustering and marker genes of Mouse Small Intestine by Visium HD.** The first column shows the clustering results on UMAP and tissue section, respectively. The second column shows the heatmap of expressions of marker genes in each cluster. The last 2 rows show the stacked violin plot of the expressions of marker genes, and the matrix plot of log-foldchange values of each marker gene in each cluster.

### Human & Mouse Immune

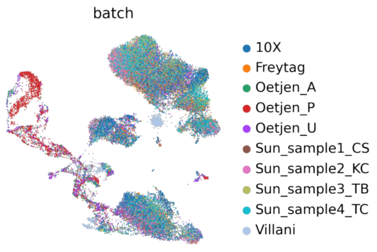

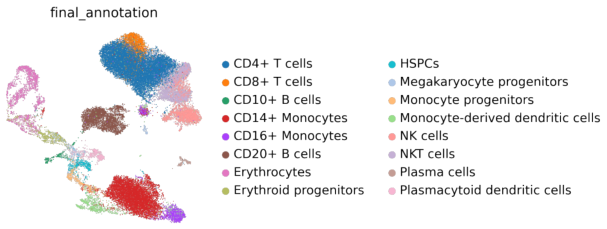

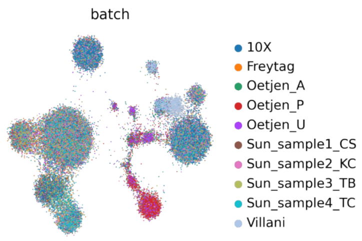

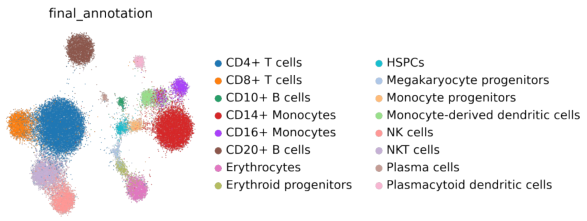

### Human  Immune

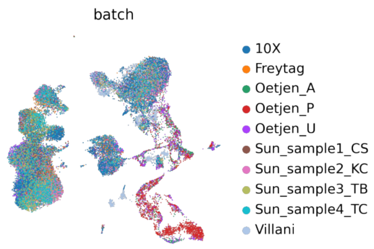

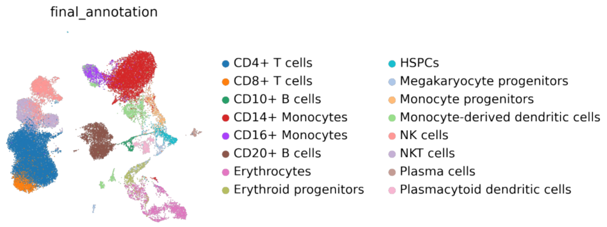

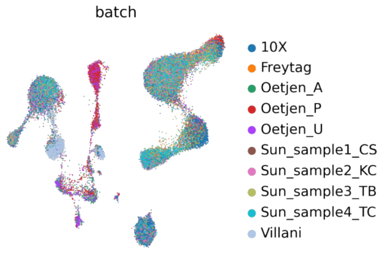

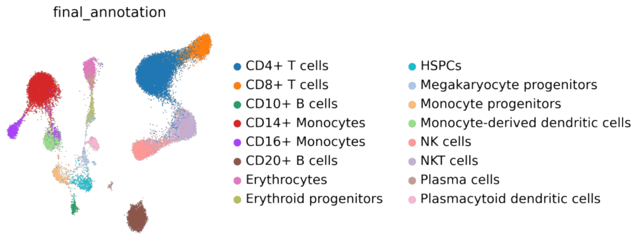

**Fig. S3 UMAP of the cell-type level integration results obtained by STEP.** Top panel: Human & Mouse Immune data. Bottom panel: Human Immune data.

#### Human Pancreas

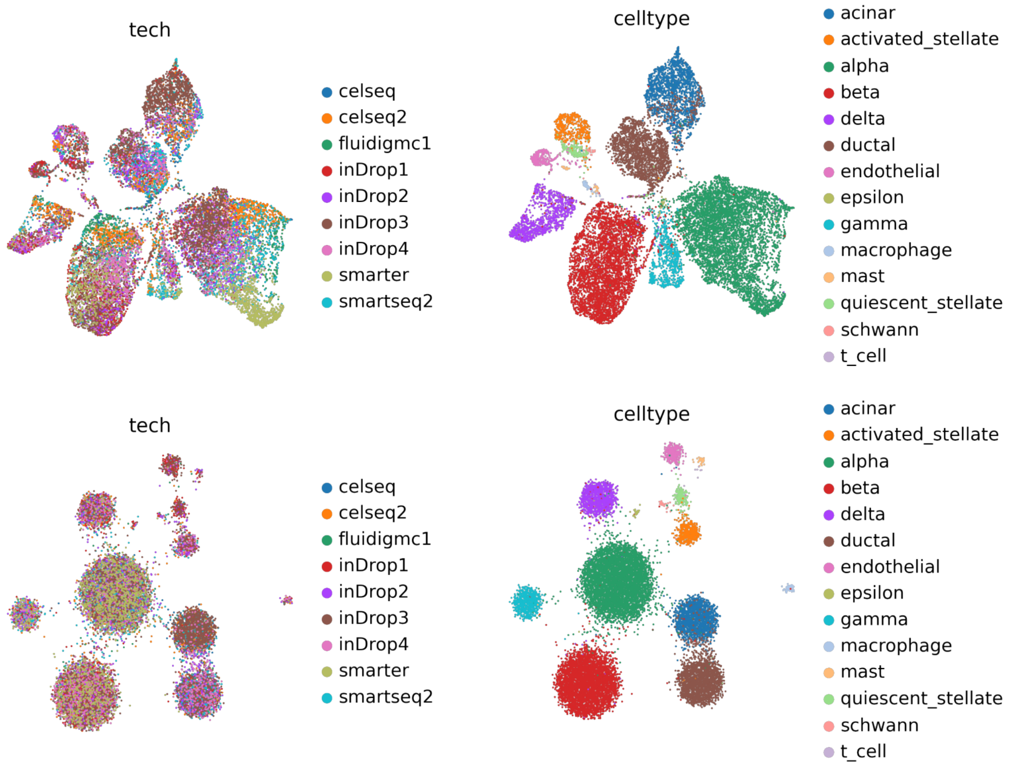

#### Human Lung

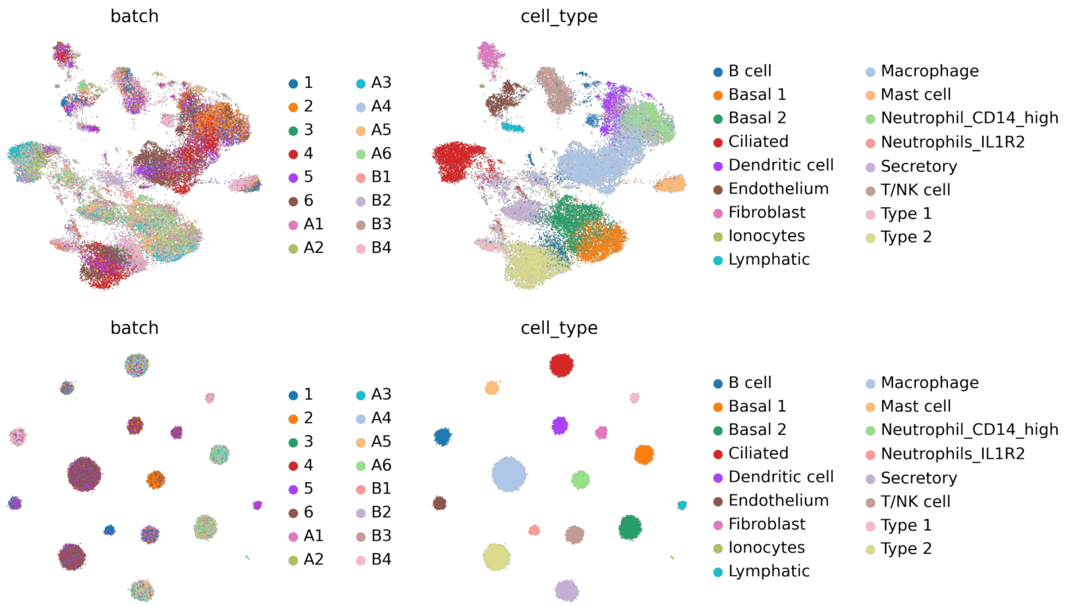

**Fig. S4 UMAP of the cell-type level integration results obtained by STEP.** Top panel: Human Pancreas. Bottom panel: Human Lung atlas.

#### Simulation 1

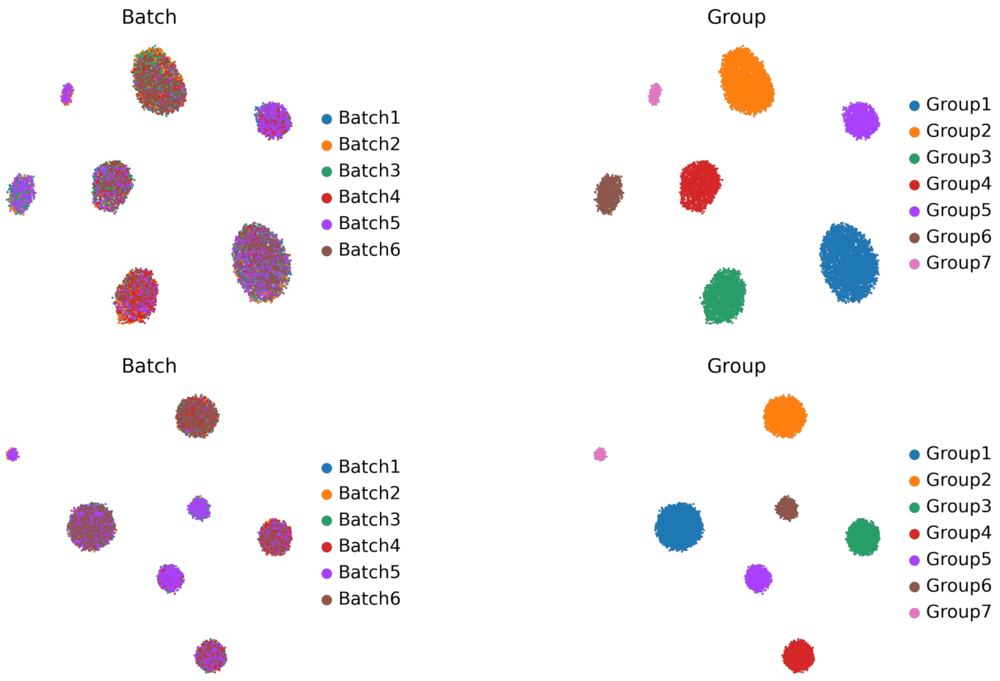

#### Simulation 2

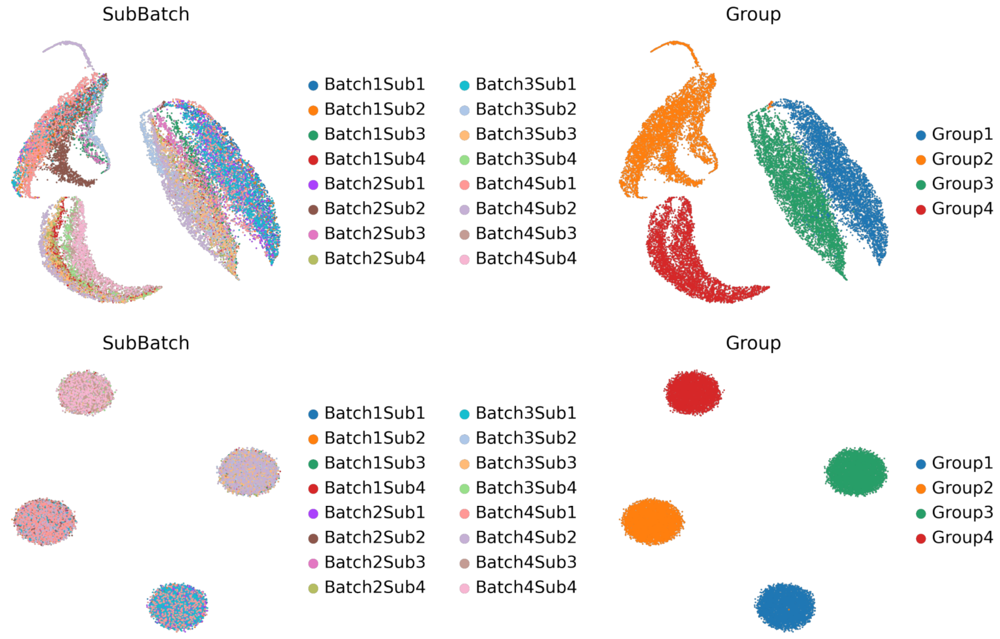

**Fig. S5 UMAP of the cell-type level integration results obtained by STEP.** Top panel: Simulation 1. Bottom panel: Simulation 2.

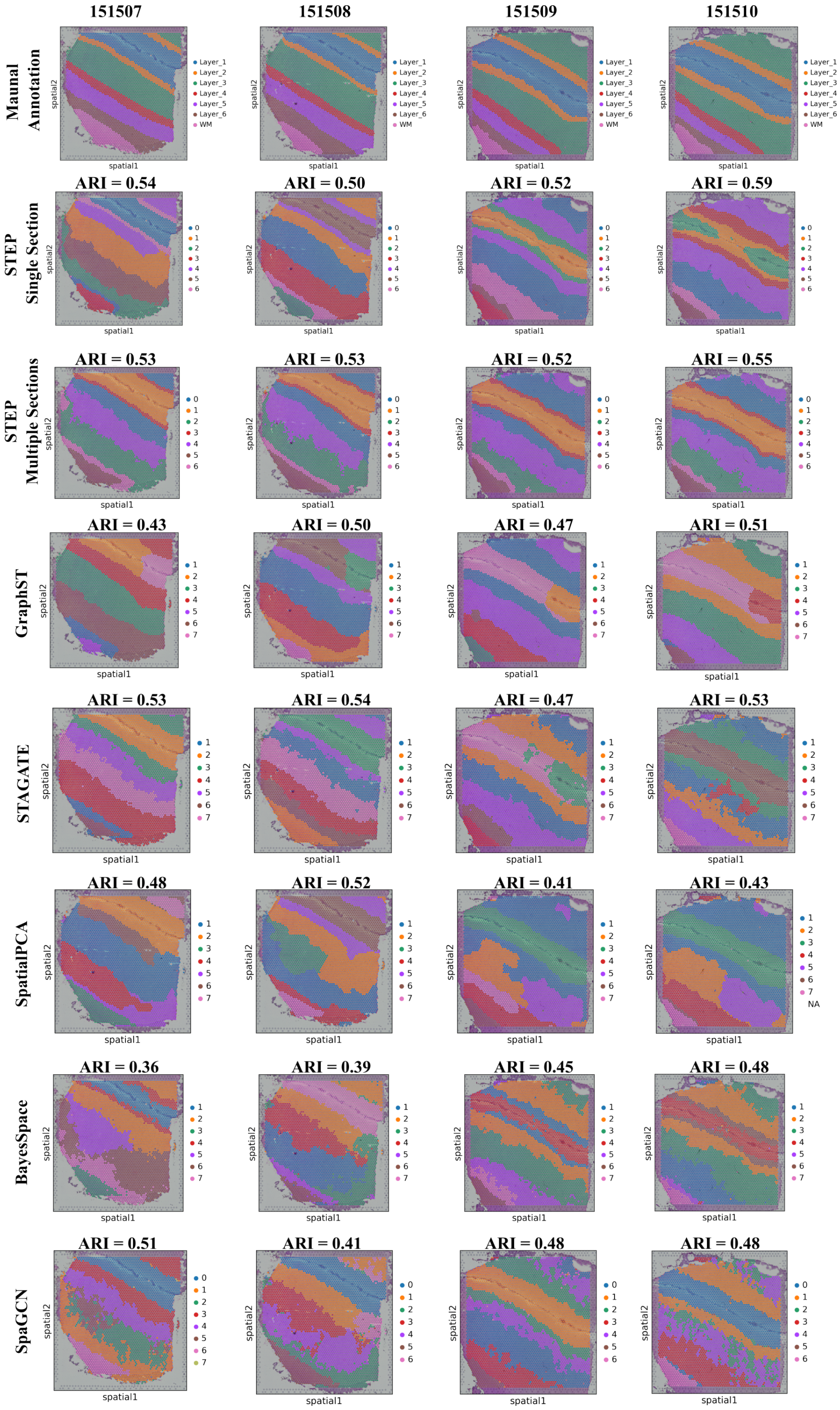

151507
151508
151509
151510
Maunal Annotation
STEP Single Section
STEP Multiple Sections
GraphST
STAGATE
SpatialPCA
BayesSpace
SpaGCN
ARI = 0.54
ARI = 0.50
ARI = 0.52
ARI = 0.59
ARI = 0.53
ARI = 0.53
ARI = 0.52
ARI = 0.55
ARI = 0.43
ARI = 0.50
ARI = 0.47
ARI = 0.51
ARI = 0.53
ARI = 0.54
ARI = 0.47
ARI = 0.53
ARI = 0.48
ARI = 0.52
ARI = 0.41
ARI = 0.43
ARI = 0.36
ARI = 0.39
ARI = 0.45
ARI = 0.48
ARI = 0.51
ARI = 0.41
ARI = 0.48
ARI = 0.48

**Fig. S6 Spatial scatter plot of the spatial domains of DLPFC dataset (151507 to 151510) obtained from manual annotations and each method.** ARI scores are displayed at the bottom of each result. All compared methods were tested individually on each section.

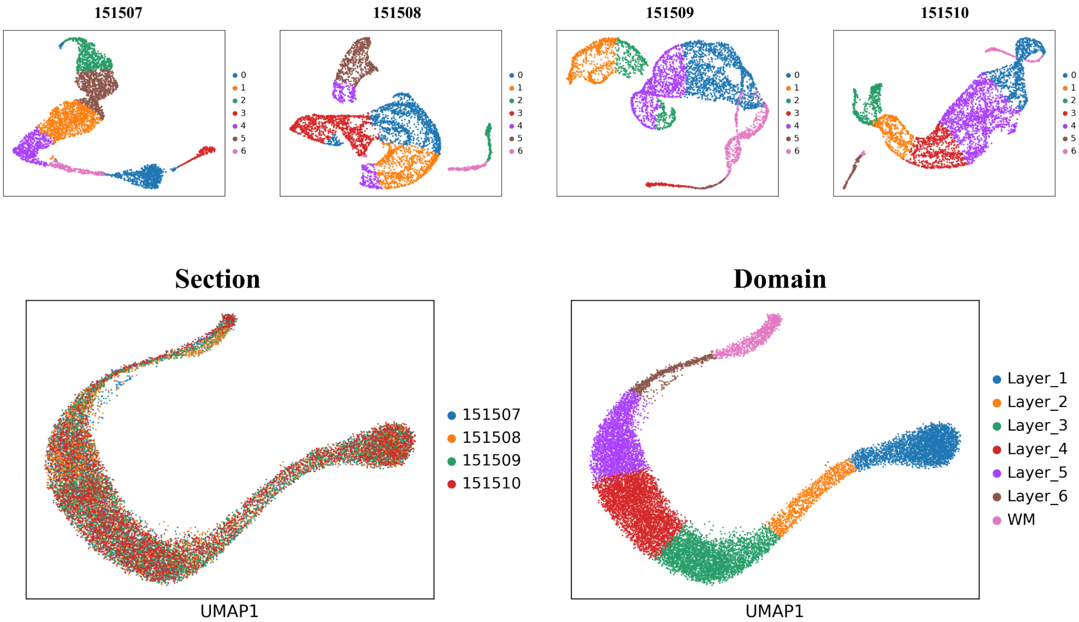

151507
151508
151509
151510
0
1
2
3
4
5
6
Section
UMAP1
Domain
Layer_1
Layer_2
Layer_3
Layer_4
Layer_5
Layer_6
WM

**Fig. S7 Spatial scatter plot of the spatial domains of DLPFC dataset (151669 to 15172) obtained from manual annotations and each method.** ARI scores are displayed at the bottom of each result. All compared methods were tested individually on each section.

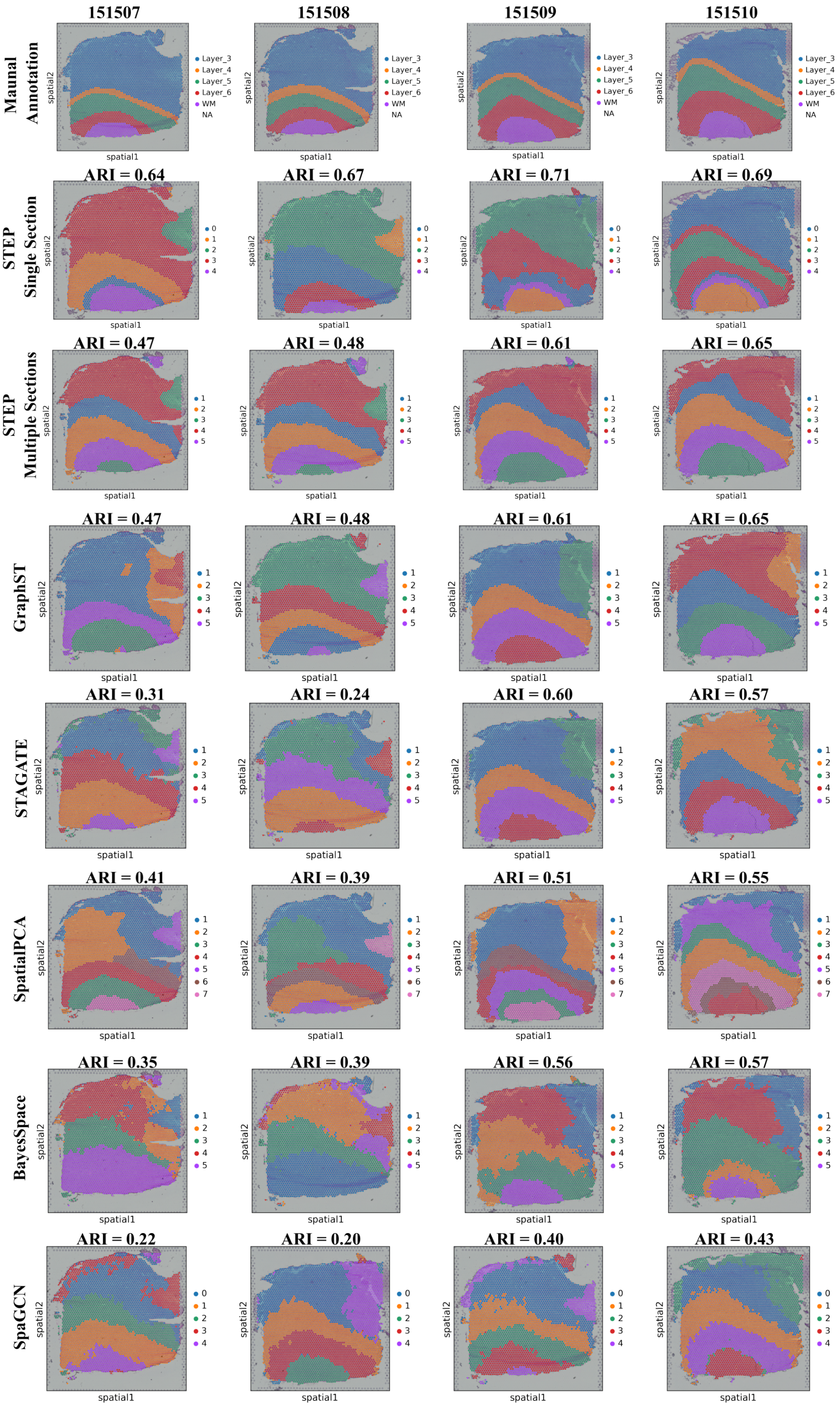
151507
151508
151509
151510
Maunal Annotation
Layer_3
Layer_4
Layer_5
Layer_6
WM
NA
spatial1
spatial2
STEP Single Section
ARI = 0.64
ARI = 0.67
ARI = 0.71
ARI = 0.69
STEP Multiple Sections
ARI = 0.47
ARI = 0.48
ARI = 0.61
ARI = 0.65
GraphST
ARI = 0.47
ARI = 0.48
ARI = 0.61
ARI = 0.65
STAGATE
ARI = 0.31
ARI = 0.24
ARI = 0.60
ARI = 0.57
SpatialPCA
ARI = 0.41
ARI = 0.39
ARI = 0.51
ARI = 0.55
BayesSpace
ARI = 0.35
ARI = 0.39
ARI = 0.56
ARI = 0.57
SpaGCN
ARI = 0.22
ARI = 0.20
ARI = 0.40
ARI = 0.43

**Fig. S8 Spatial scatter plot of the spatial domains of DLPFC dataset (151673 to 15176) obtained from manual annotations and each method.** ARI scores are displayed at the bottom of each result. All compared methods were tested individually on each section.

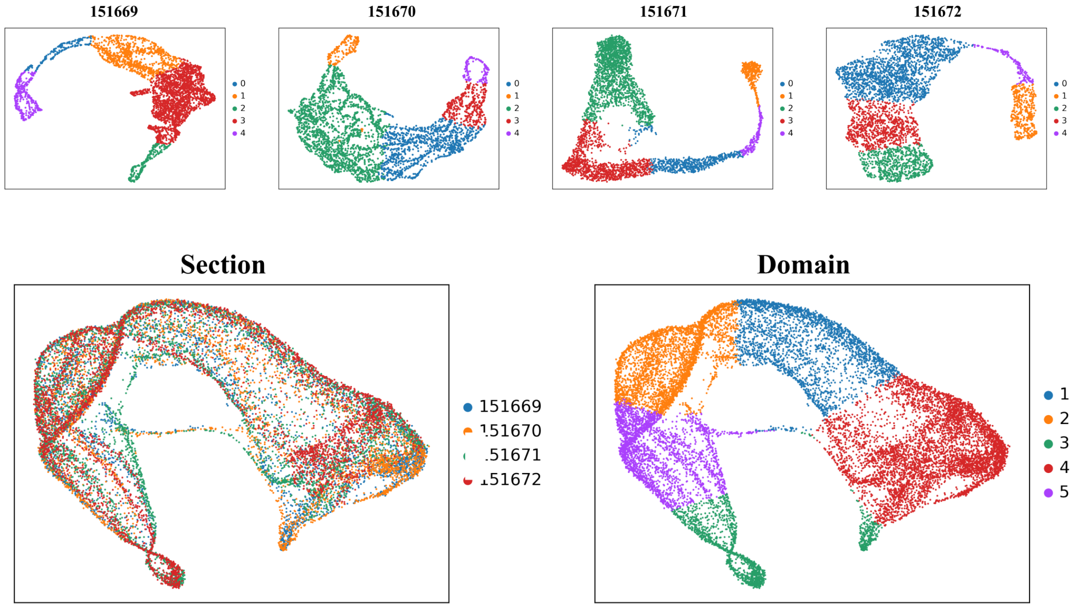
151669
151670
151671
151672
0
1
2
3
4
Section
151669
Domain
1
2
3
4
5

**Fig. S9 UMAPs of the embedding obtained by STEP and are colored by spatial domains identified by STEP.** Top panel: results of STEP when tested in single section. Bottom panel: results of STEP when tested in the integration of multiple sections.

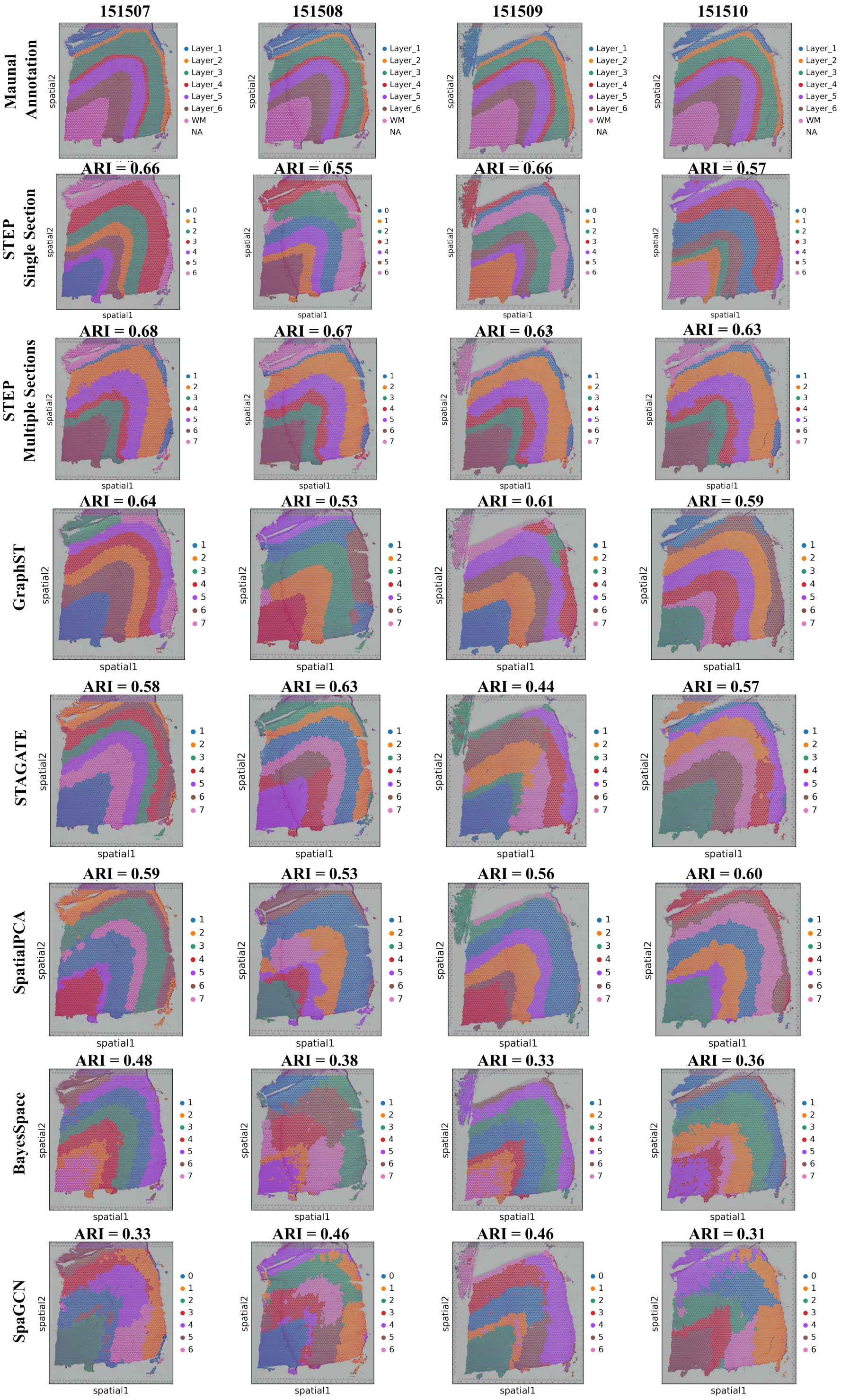

151507
151508
151509
151510
Maunal Annotation
STEP Single Section
STEP Multiple Sections
GraphST
STAGATE
SpatialPCA
BayesSpace
SpaGCN
ARI = 0.66
ARI = 0.55
ARI = 0.66
ARI = 0.57
ARI = 0.68
ARI = 0.67
ARI = 0.63
ARI = 0.63
ARI = 0.64
ARI = 0.53
ARI = 0.61
ARI = 0.59
ARI = 0.58
ARI = 0.63
ARI = 0.44
ARI = 0.57
ARI = 0.59
ARI = 0.53
ARI = 0.56
ARI = 0.60
ARI = 0.48
ARI = 0.38
ARI = 0.33
ARI = 0.36
ARI = 0.33
ARI = 0.46
ARI = 0.46
ARI = 0.31
Layer_1
Layer_2
Layer_3
Layer_4
Layer_5
Layer_6
WM
NA
spatial1
spatial2

**Fig. S10 UMAPs of the embedding obtained by STEP and are colored by spatial domains identified by STEP.** Top panel: results of STEP when tested in single section. Bottom panel: results of STEP when tested in the integration of multiple sections.

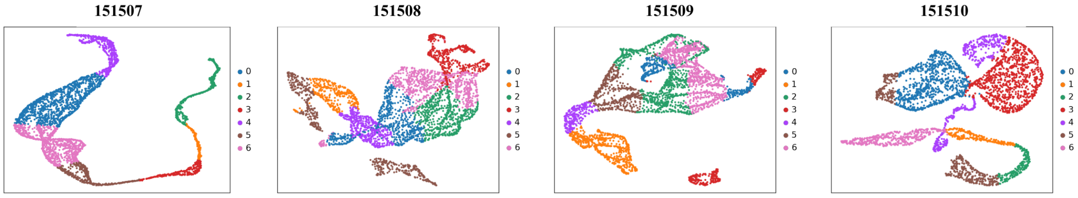
151507
151508
151509
151510
0
1
2
3
4
5
6

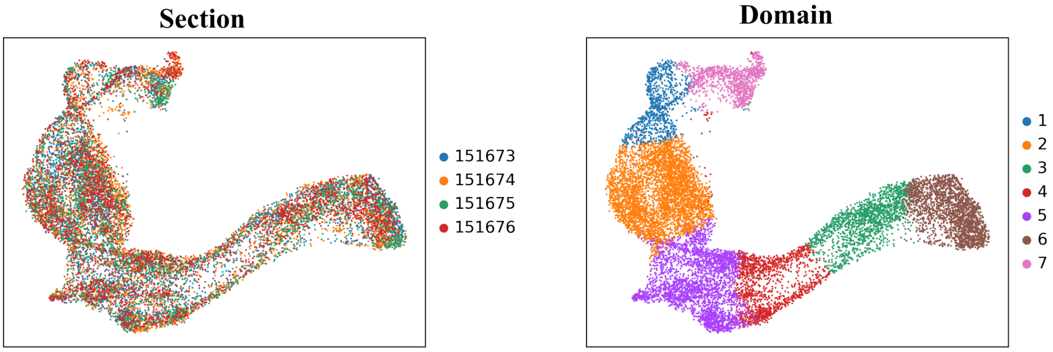
Section
151673
151674
151675
151676
Domain
1
2
3
4
5
6
7

**Fig. S11 UMAPs of the embedding obtained by STEP and are colored by spatial domains identified by STEP.** Top panel: results of STEP when tested in single section. Bottom panel: results of STEP when tested in the integration of multiple sections.

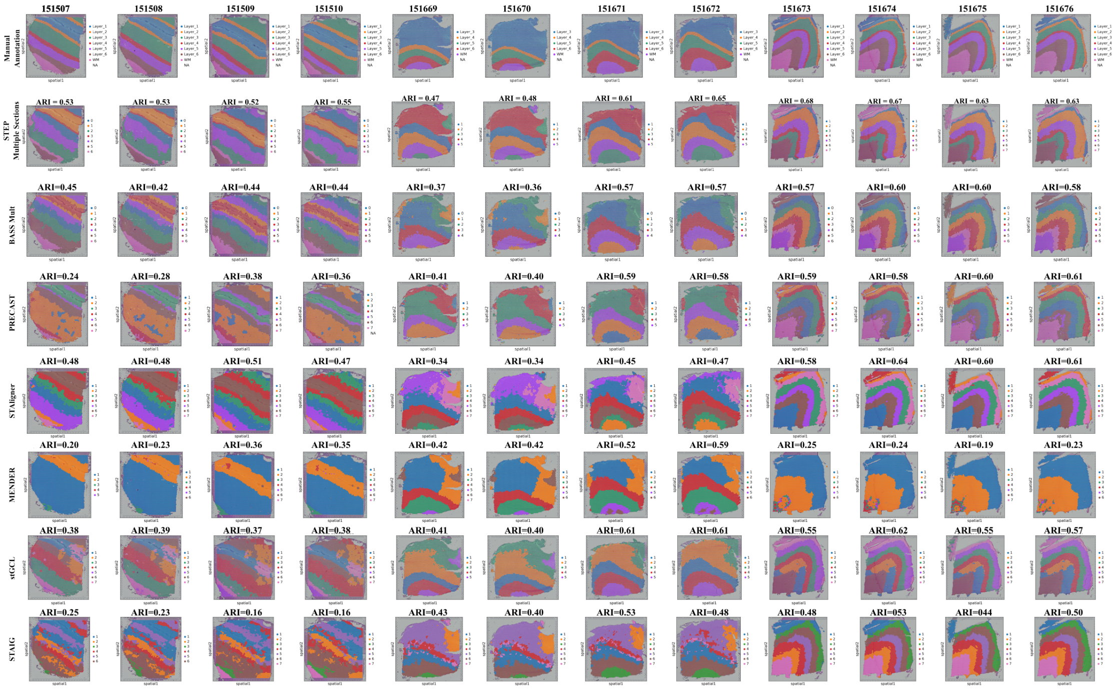
151507
151508
151509
151510
151669
151670
151671
151672
151673
151674
151675
151676
Manual Annotation
STEP Multiple Sections
ARI = 0.53
ARI = 0.53
ARI = 0.52
ARI = 0.55
ARI = 0.47
ARI = 0.48
ARI = 0.61
ARI = 0.65
ARI = 0.68
ARI = 0.67
ARI = 0.63
ARI = 0.63
BASS Mult
ARI=0.45
ARI=0.42
ARI=0.44
ARI=0.44
ARI=0.37
ARI=0.36
ARI=0.57
ARI=0.57
ARI=0.57
ARI=0.60
ARI=0.60
ARI=0.58
PRECAST
ARI=0.24
ARI=0.28
ARI=0.38
ARI=0.36
ARI=0.41
ARI=0.40
ARI=0.59
ARI=0.58
ARI=0.59
ARI=0.58
ARI=0.60
ARI=0.61
STAligner
ARI=0.48
ARI=0.48
ARI=0.51
ARI=0.47
ARI=0.34
ARI=0.34
ARI=0.45
ARI=0.47
ARI=0.58
ARI=0.64
ARI=0.60
ARI=0.61
MENDER
ARI=0.20
ARI=0.23
ARI=0.36
ARI=0.35
ARI=0.42
ARI=0.42
ARI=0.52
ARI=0.59
ARI=0.25
ARI=0.24
ARI=0.19
ARI=0.23
stGCL
ARI=0.38
ARI=0.39
ARI=0.37
ARI=0.38
ARI=0.41
ARI=0.40
ARI=0.61
ARI=0.61
ARI=0.55
ARI=0.62
ARI=0.55
ARI=0.57
STAIG
ARI=0.25
ARI=0.23
ARI=0.16
ARI=0.16
ARI=0.43
ARI=0.40
ARI=0.53
ARI=0.48
ARI=0.48
ARI=053
ARI=044
ARI=0.50

**Fig. S12 Comparison of STEP, BASS, PRECAST, STAligner, MENDER, stGCL and STAIG in the scenario of integrating multiple sections from DLPFC dataset.**

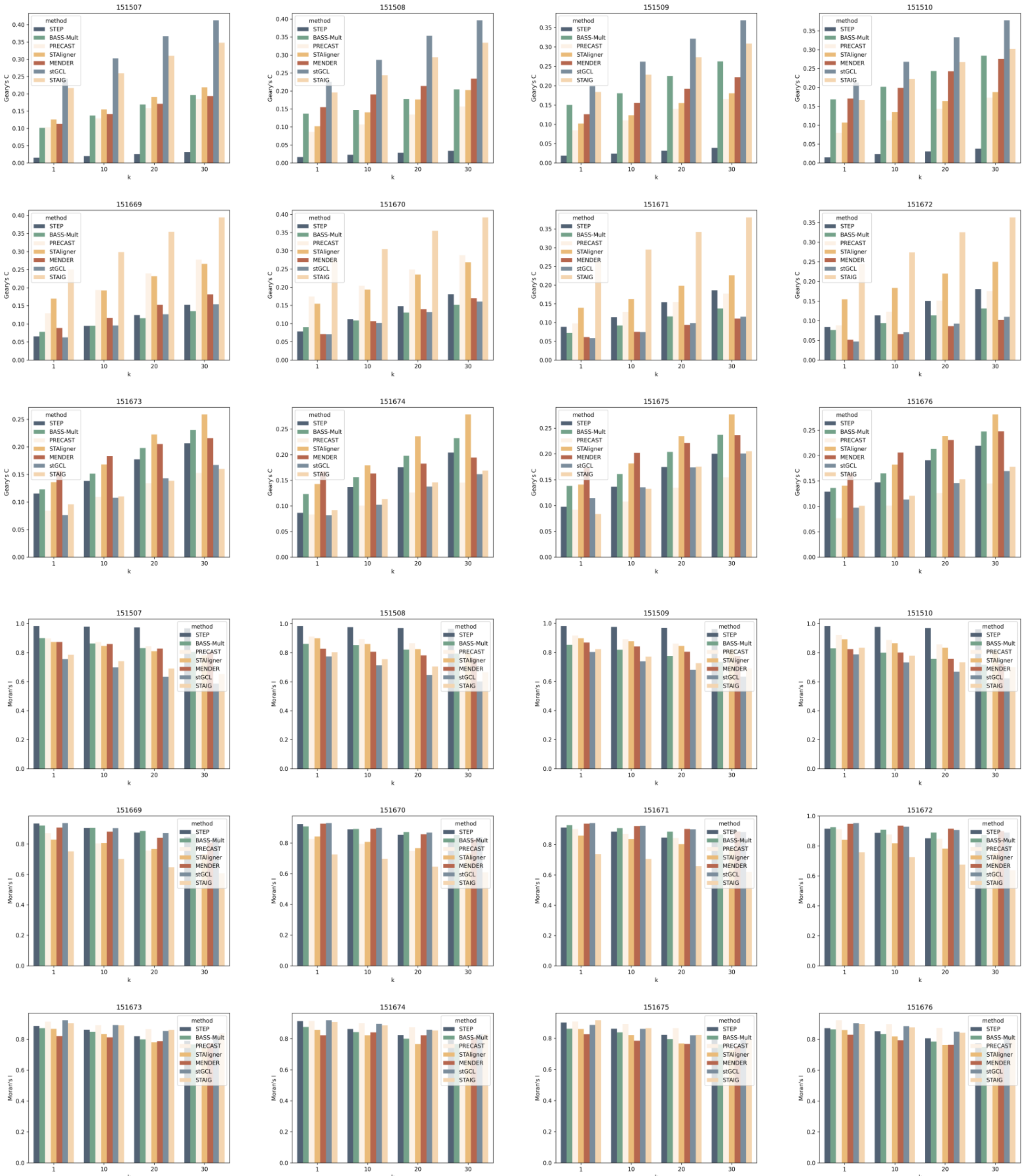
151507
151508
151509
151510
151669
151670
151671
151672
151673
151674
151675
151676
method
STEP
BASS-Mult
PRECAST
STAligner
MENDER
stGCL
STAIG
Geary's C
Moran's I
k

**Fig. S13 Geary's C and Moran's I scores for DLFPC dataset calculated based on the identified domains from each method in multi-section scenario.** First 4 rows: Spatial domains identified in DLPFC dataset across sections; Last two rows: Geary's C and Moran's I calculated based on identified domains of each method based on different kNN settings (k = 1, 10, 20, 30).

# P1: 151507 to 151510

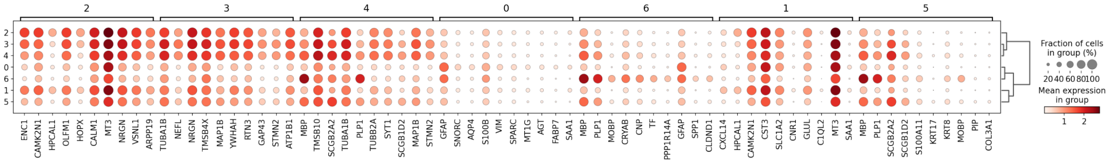

# P2: 151669 to 151672

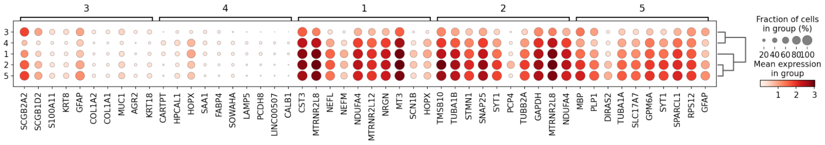

# P3: 151673 to 151676

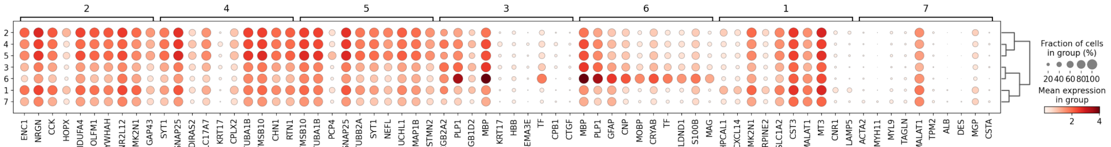

**Fig. S14 Results of spatially differentially expressed genes analysis conducted on each group: P1, P2, P3 in DLPFC dataset.** And it is displayed by the dot plot of gene expression level in each domain.

# P1: 151507 to 151510

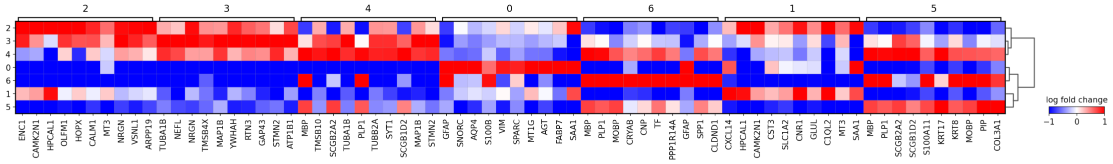

# P2: 151669 to 151672

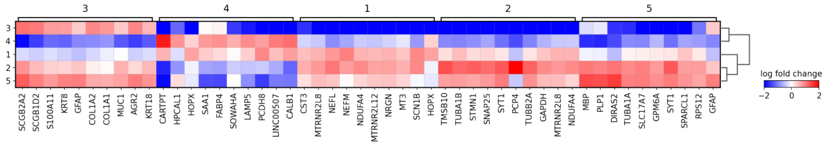

# P3: 151672 to 151676

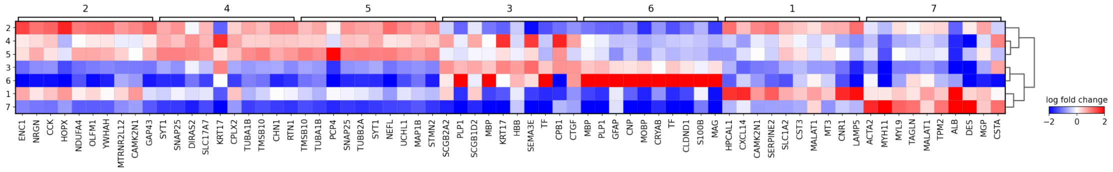

**Fig. S15 Results of spatially differentially expressed genes analysis conducted on each group: P1, P2, P3 in DLPFC dataset.** And it is displayed by the matrix plot of log fold changes of gene expression in each domain.

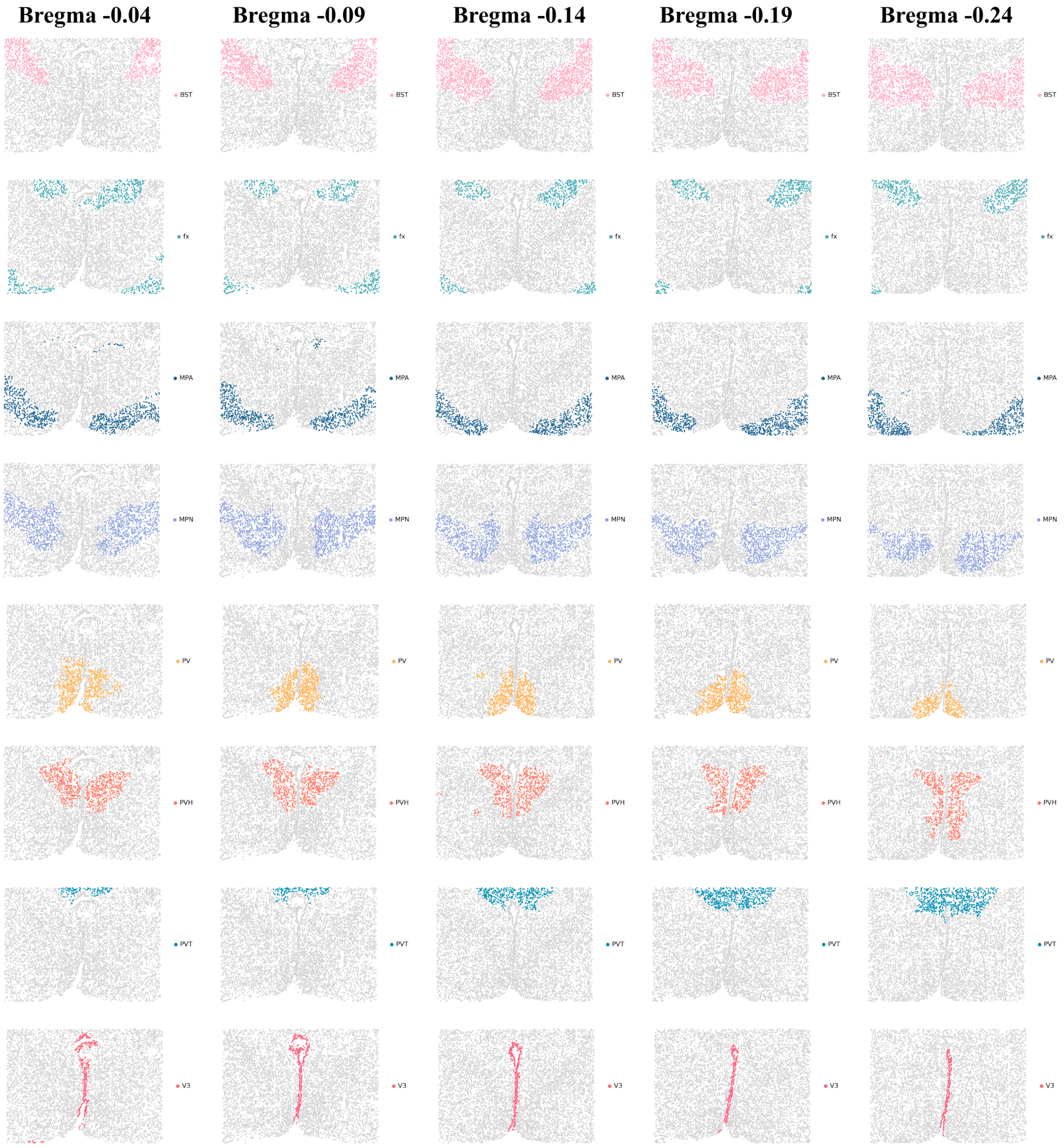

Bregma -0.04
Bregma -0.09
Bregma -0.14
Bregma -0.19
Bregma -0.24
BST
fx
MPA
MPN
PV
PVH
PVT
V3

**Fig. S16 Annotated spatial domains identified by STEP across 5 sections: Bregma -0.04, -0.09, -0.14, -0.19 and -0.24 of Mouse hypothalamus data by MERFISH.** Each row represents one identified and annotated domain across all sections.

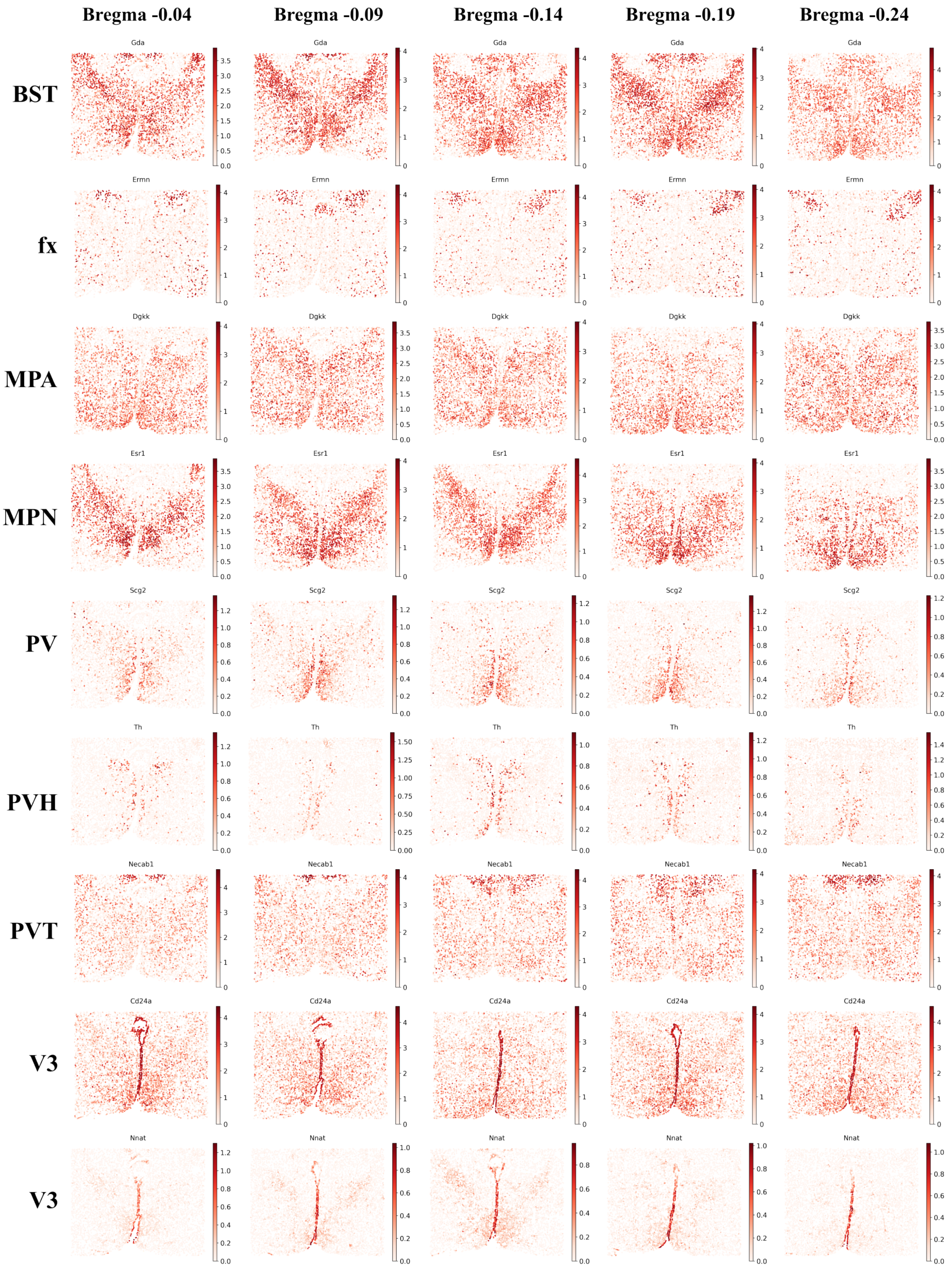
Bregma -0.04
Bregma -0.09
Bregma -0.14
Bregma -0.19
Bregma -0.24
BST
Gda
fx
Ermn
MPA
Dgkk
MPN
Esr1
PV
Scg2
PVH
Th
PVT
Necab1
V3
Cd24a
V3
Nnat

**Fig. S17 Expression of spatially and differentially expressed genes obtained by differentially expressed genes analysis.**

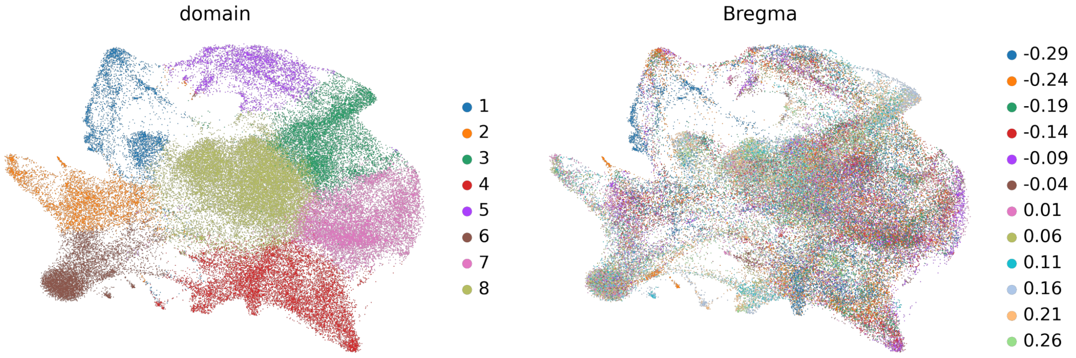
domain
1
2
3
4
5
6
7
8
Bregma
-0.29
-0.24
-0.19
-0.14
-0.09
-0.04
0.01
0.06
0.11
0.16
0.21
0.26

**Fig. S18 Results of the integration of 12 sections, obtained by STEP, of the animal 7 in original dataset (Mouse hypothalamus data by MERFISH), ranging from Bregma -0.29 to 0.26.** UMAPs of the embeddings colored by identified spatial domains (left) and section id (right).

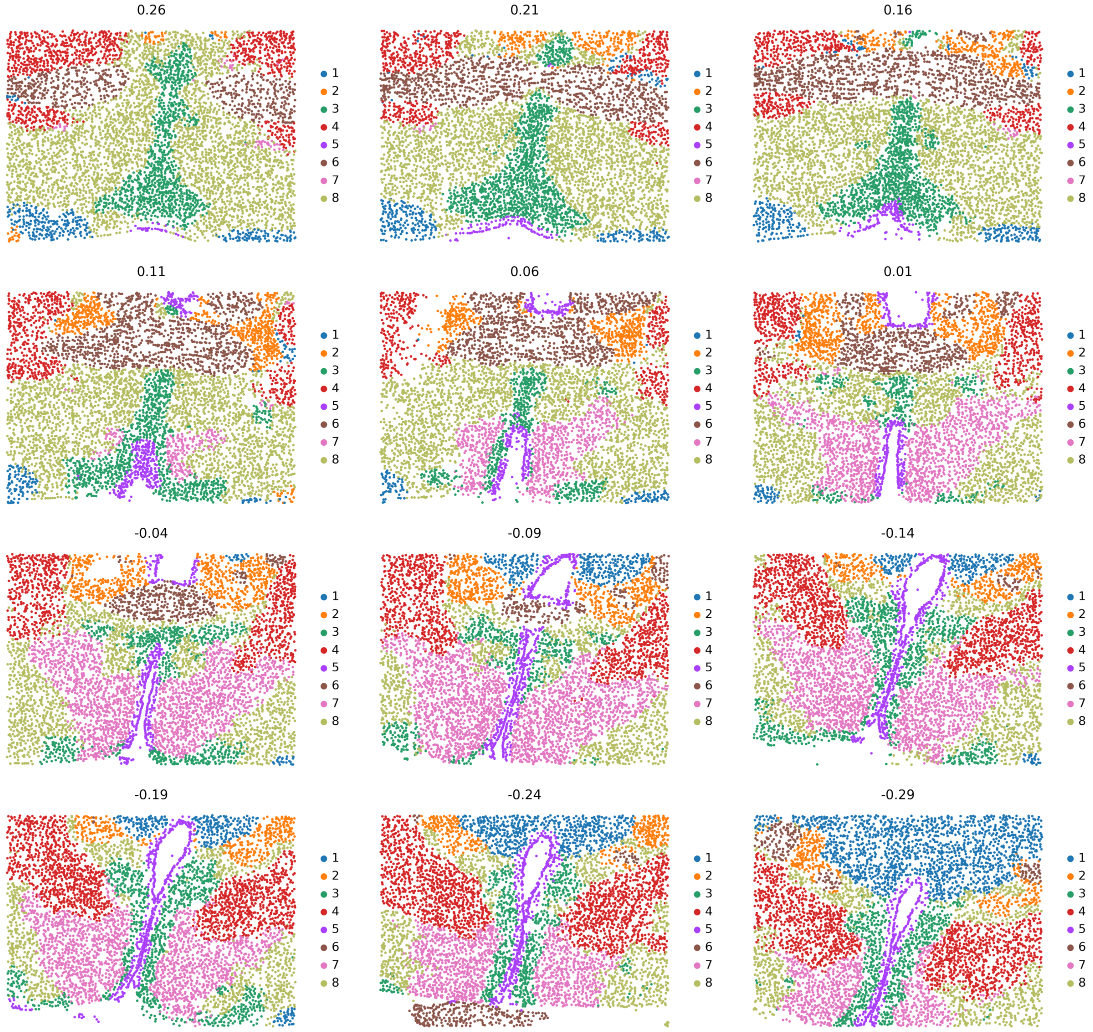
0.26
0.21
0.16
0.11
0.06
0.01
-0.04
-0.09
-0.14
-0.19
-0.24
-0.29
1
2
3
4
5
6
7
8

**Fig. S19 Results of the integration of 12 sections, obtained by STEP, of the animal 7 in original dataset, ranging from Bregma -0.29 to 0.26.** Spatial domains identified across 12 sections.

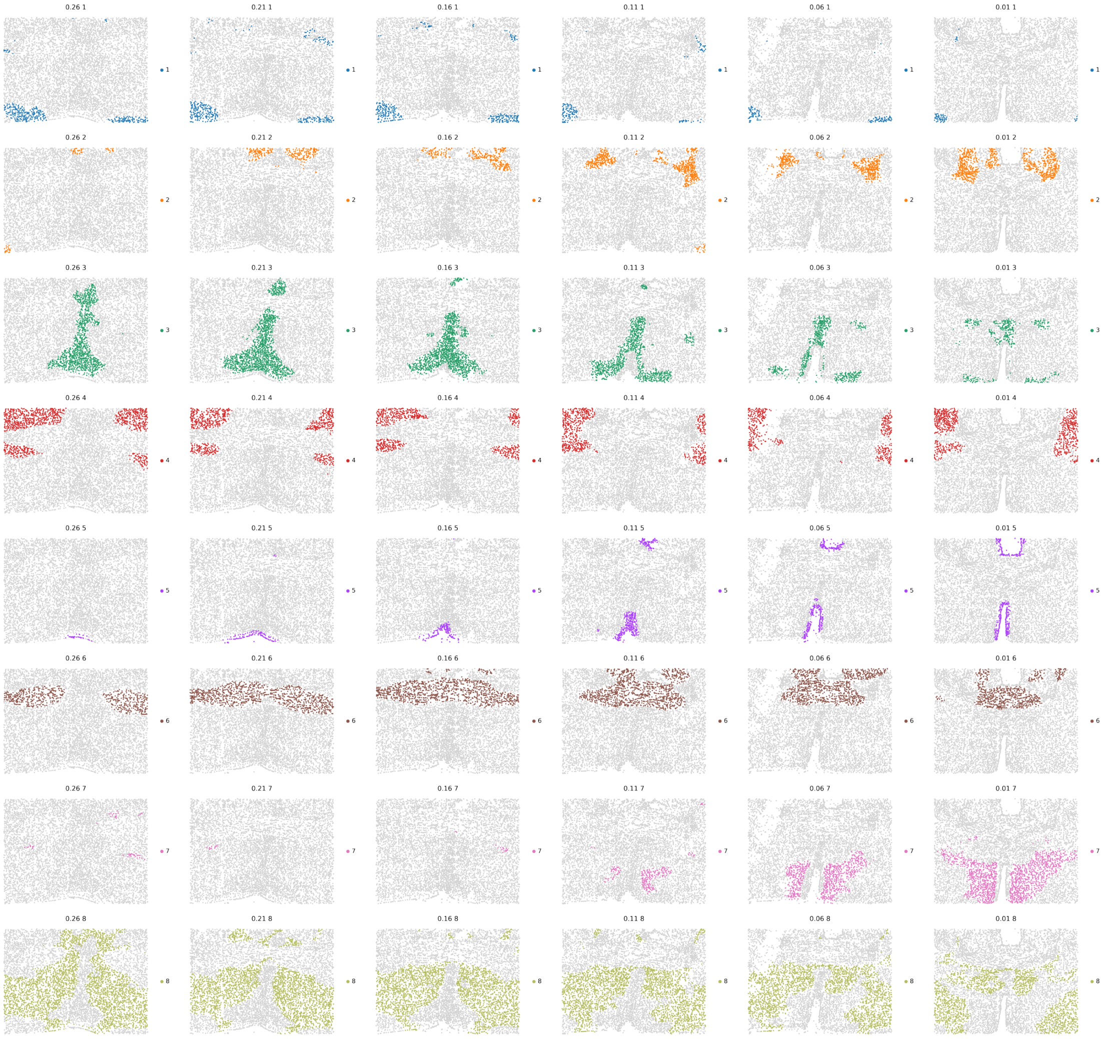
0.26 1
0.21 1
0.16 1
0.11 1
0.06 1
0.01 1
0.26 2
0.21 2
0.16 2
0.11 2
0.06 2
0.01 2
0.26 3
0.21 3
0.16 3
0.11 3
0.06 3
0.01 3
0.26 4
0.21 4
0.16 4
0.11 4
0.06 4
0.01 4
0.26 5
0.21 5
0.16 5
0.11 5
0.06 5
0.01 5
0.26 6
0.21 6
0.16 6
0.11 6
0.06 6
0.01 6
0.26 7
0.21 7
0.16 7
0.11 7
0.06 7
0.01 7
0.26 8
0.21 8
0.16 8
0.11 8
0.06 8
0.01 8

**Fig. S20 Spatial domains identified by STEP in the sections from Bregma 0.26 to 0.01.** Each row represents one domain across all sections.

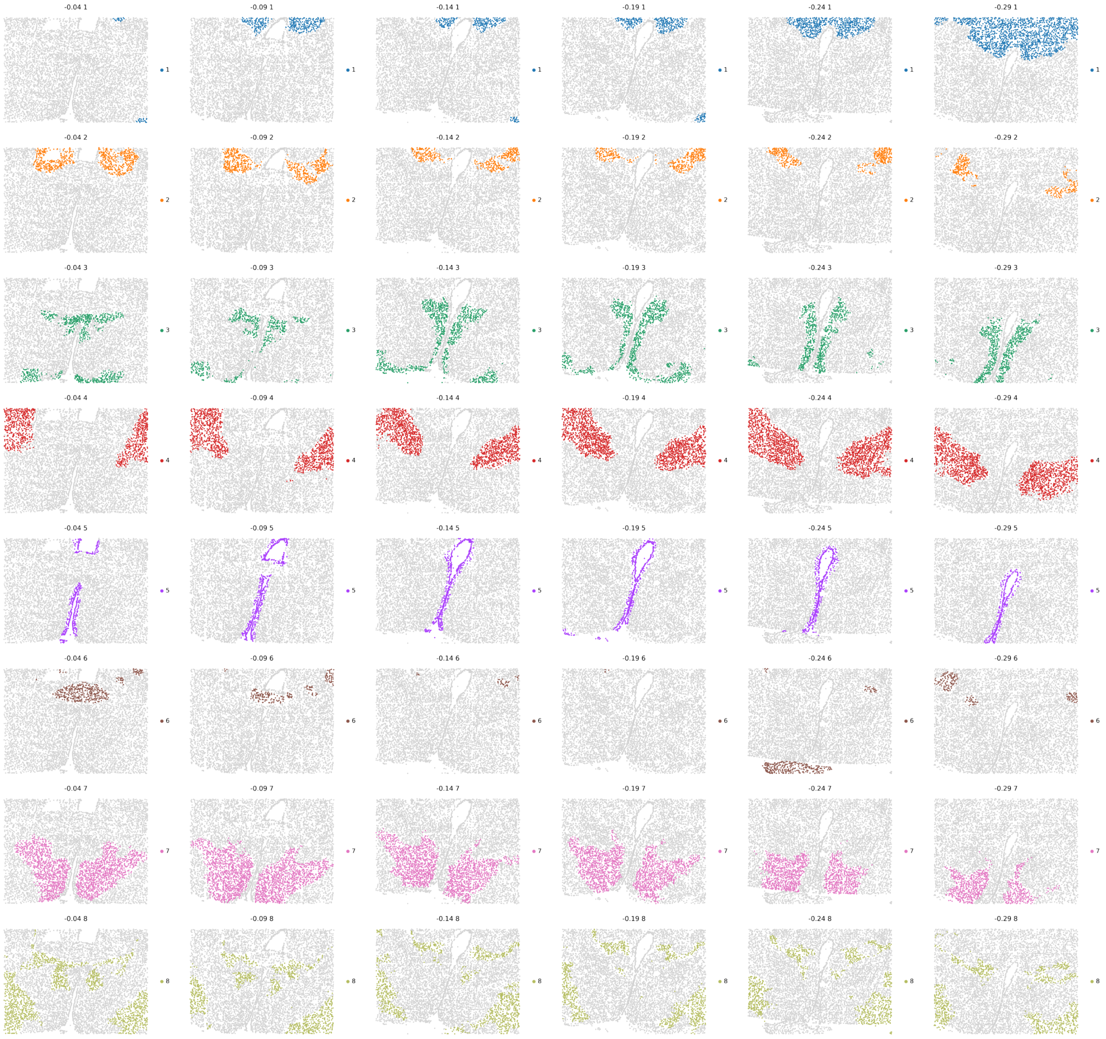
-0.04 1
-0.09 1
-0.14 1
-0.19 1
-0.24 1
-0.29 1
-0.04 2
-0.09 2
-0.14 2
-0.19 2
-0.24 2
-0.29 2
-0.04 3
-0.09 3
-0.14 3
-0.19 3
-0.24 3
-0.29 3
-0.04 4
-0.09 4
-0.14 4
-0.19 4
-0.24 4
-0.29 4
-0.04 5
-0.09 5
-0.14 5
-0.19 5
-0.24 5
-0.29 5
-0.04 6
-0.09 6
-0.14 6
-0.19 6
-0.24 6
-0.29 6
-0.04 7
-0.09 7
-0.14 7
-0.19 7
-0.24 7
-0.29 7
-0.04 8
-0.09 8
-0.14 8
-0.19 8
-0.24 8
-0.29 8

**Fig. S21 Spatial domains identified by STEP in the sections from Bregma -0.04 to -0.29.** Each row represents one domain across all sections.

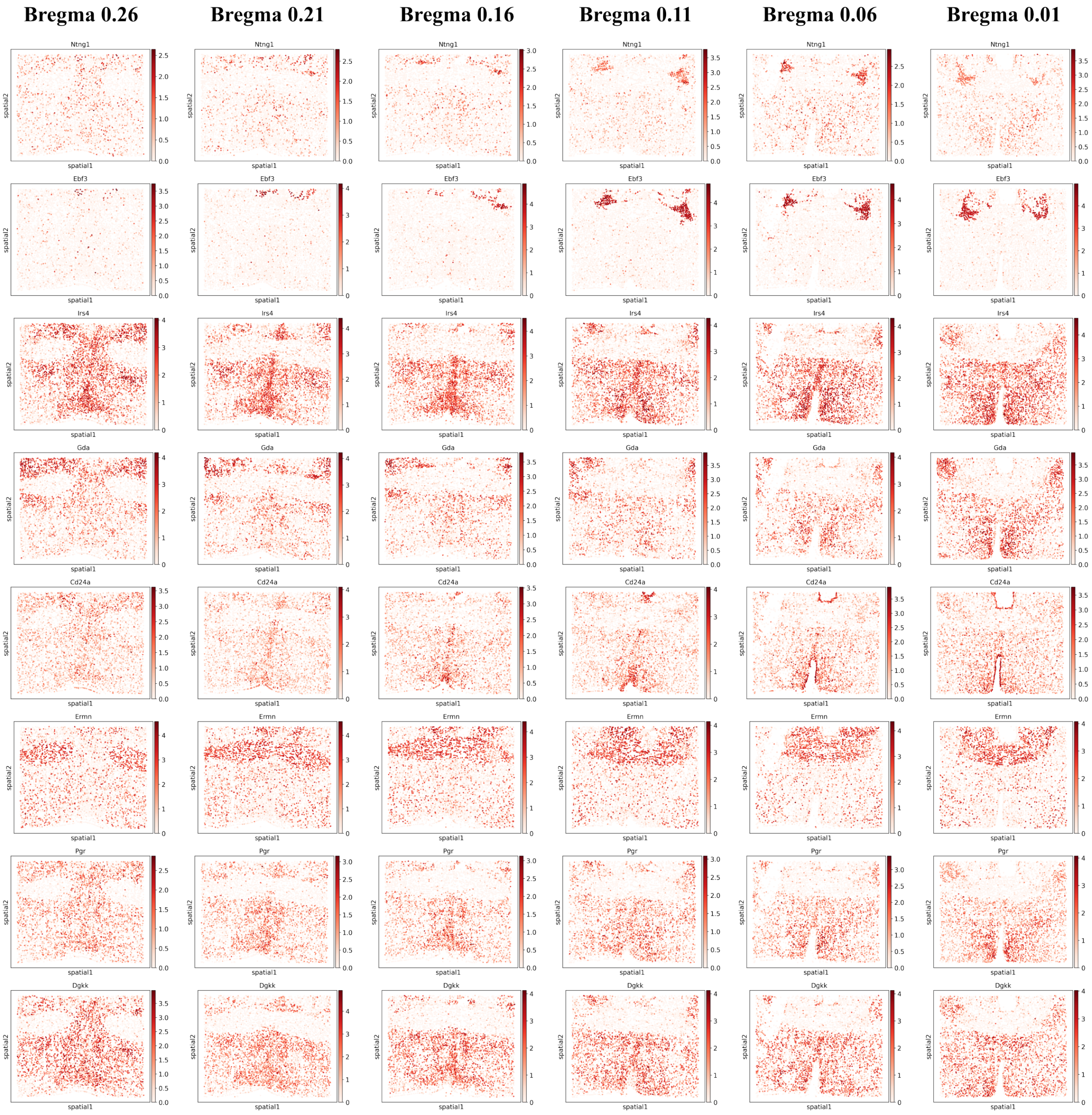

Bregma 0.26
Bregma 0.21
Bregma 0.16
Bregma 0.11
Bregma 0.06
Bregma 0.01
Ntng1
Ebf3
Irs4
Gda
Cd24a
Ermn
Pgr
Dgkk
spatial1
spatial2

**Fig. S22 Expressions of spatially differentially expressed genes (sections of Bregma 0.26 to 0.01)** detected across sections by performing differentially expressed genes analysis referring to identified spatial domains.

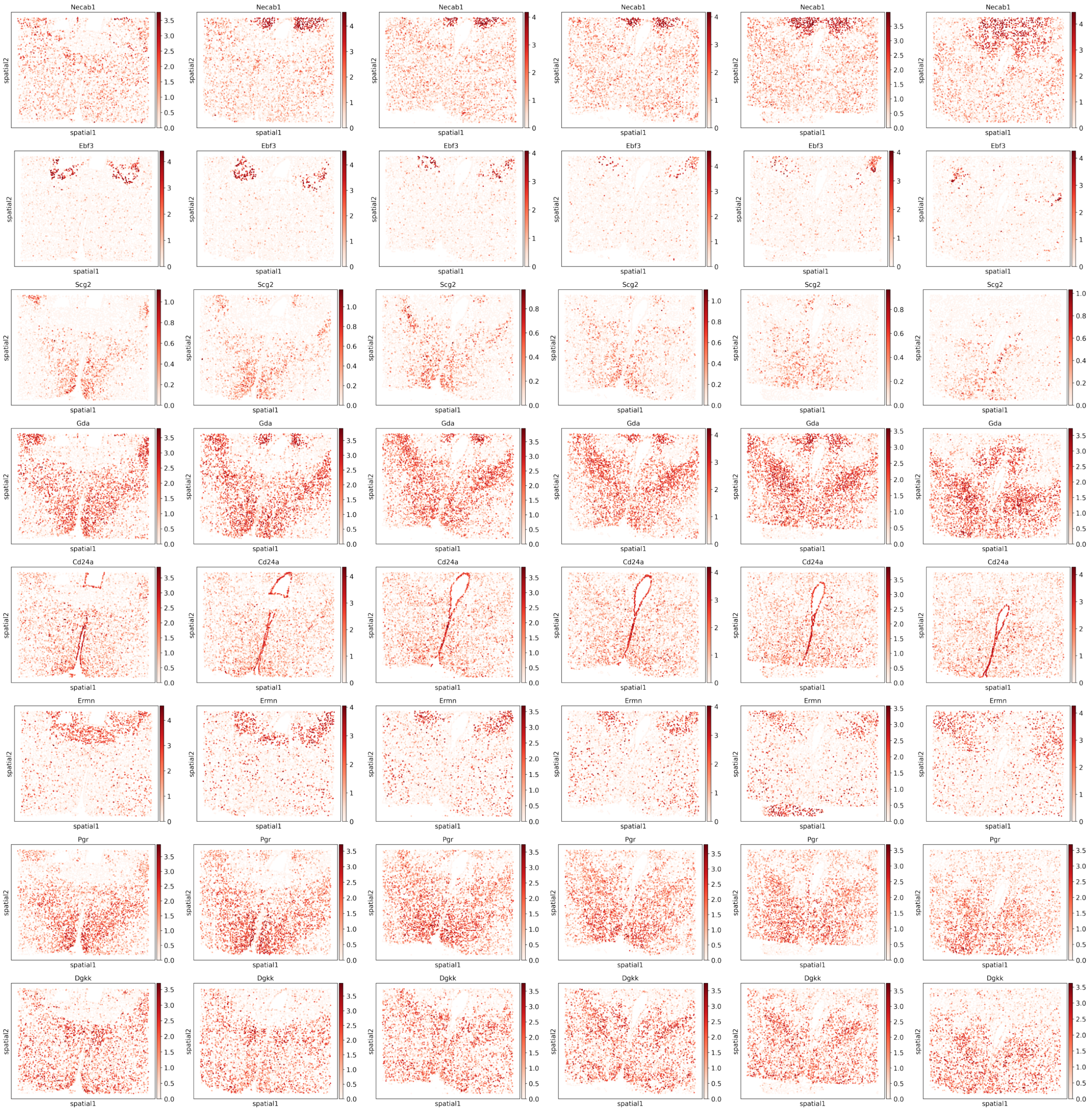
Necab1
Ebf3
Scg2
Gda
Cd24a
Ermn
Pgr
Dgkk
spatial1
spatial2

**Fig. S23 Expressions of spatially differentially expressed genes (sections of Bregma -0.04 to -0.29)** detected across sections by performing differentially expressed genes analysis referring to identified spatial domains.

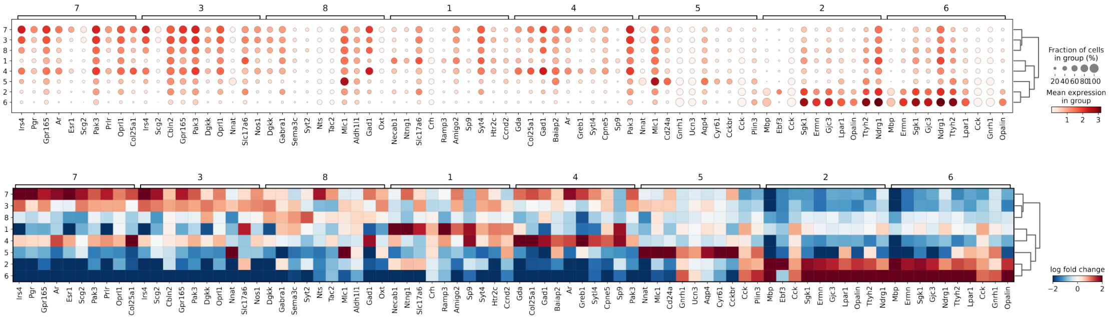

7
3
8
1
4
5
2
6
Irs4 Pgr Gpr165 Ar Esr1 Scg2 Pak3 Prlr Oprl1 Col25a1
Irs4 Scg2 Cbln2 Gpr165 Pak3 Dgkk Oprl1 Nnat Slc17a6 Nos1
Dgkk Gabra1 Sema3c Syt2 Nts Tac2 Mlc1 Aldh1l1 Gad1 Oxt
Necab1 Ntng1 Slc17a6 Crh Ramp3 Amigo2 Sp9 Syt4 Htr2c Ccnd2
Gda Col25a1 Gad1 Baiap2 Ar Greb1 Sytl4 Cpne5 Sp9 Pak3
Nnat Mlc1 Cd24a Gnrh1 Ucn3 Aqp4 Cyr61 Cckbr Cck Plin3
Mbp Ebf3 Cck Sgk1 Ermn Gjc3 Lpar1 Opalin Ttyh2 Ndrg1
Mbp Ermn Sgk1 Gjc3 Ndrg1 Ttyh2 Lpar1 Cck Gnrh1 Opalin
Fraction of cells in group (%)
20 40 60 80 100
Mean expression in group
1 2 3
log fold change
−2 0 2

**Fig. S24 Results of spatially differentially expressed genes analysis (sections of Bregma -0.04 to -0.29)**. Top panel: dot plot of the average gene expression levels and proportions of expressed genes in each domain; Bottom panel: matrix plot of the average log fold changes of the gene expression levels in each domain when referring to the rest domains.

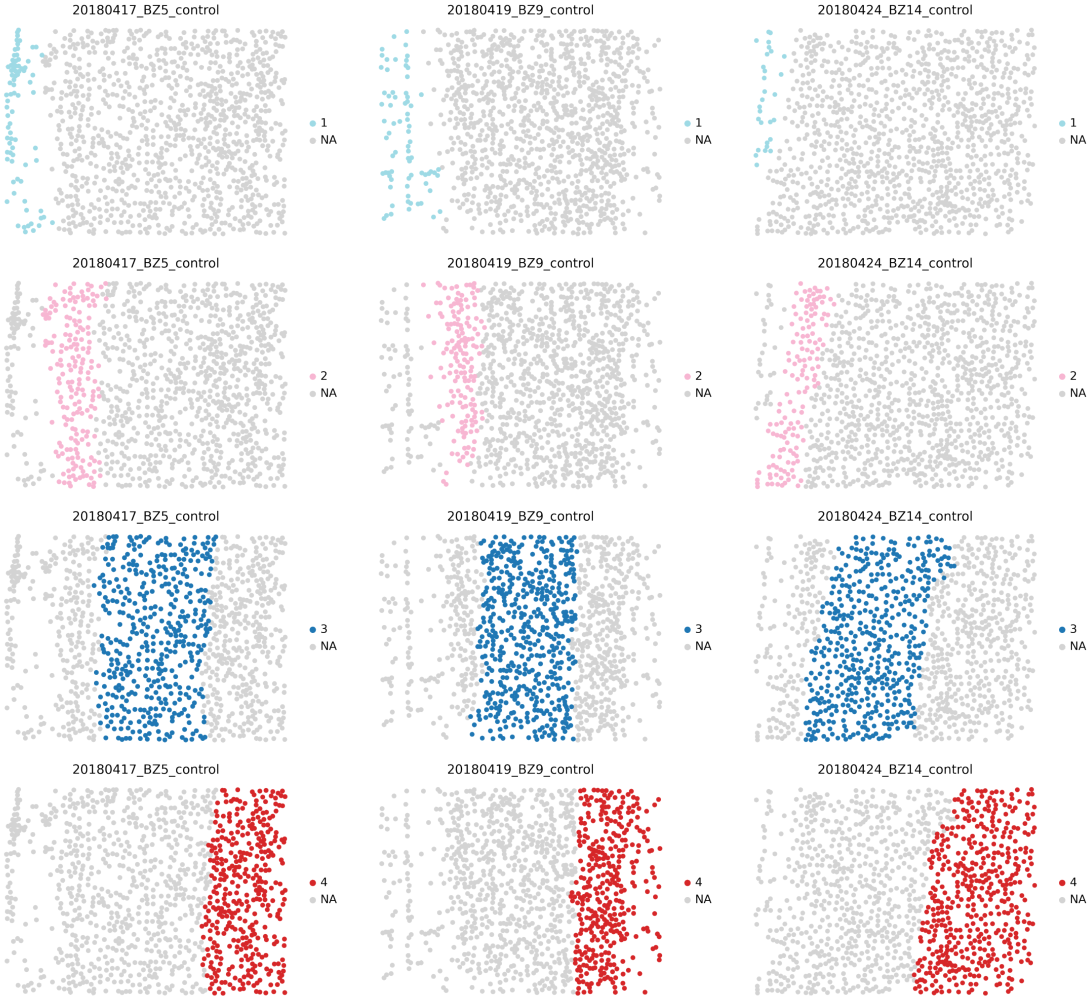
20180417_BZ5_control
1
NA
20180419_BZ9_control
1
NA
20180424_BZ14_control
1
NA
20180417_BZ5_control
2
NA
20180419_BZ9_control
2
NA
20180424_BZ14_control
2
NA
20180417_BZ5_control
3
NA
20180419_BZ9_control
3
NA
20180424_BZ14_control
3
NA
20180417_BZ5_control
4
NA
20180419_BZ9_control
4
NA
20180424_BZ14_control
4
NA

**Fig. S25 Spatial domains identified across sections (BZ5, BZ9, BZ14) of Mouse medial prefrontal cortex by STARmap after being integrated by STEP.** Each row represents one spatial domain across sections.

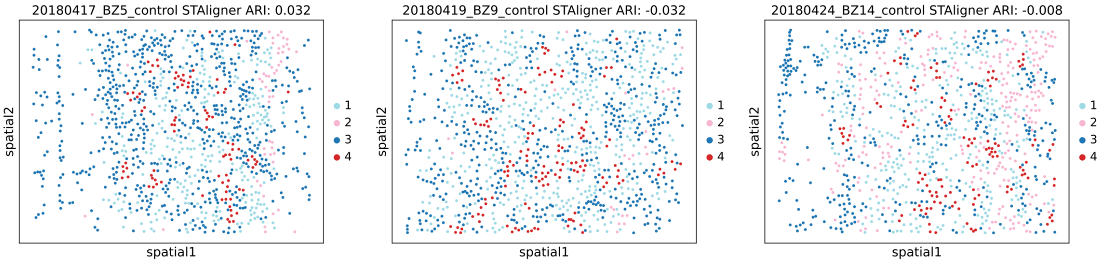
20180417_BZ5_control STAligner ARI: 0.032
spatial2
spatial1
1
2
3
4
20180419_BZ9_control STAligner ARI: -0.032
spatial2
spatial1
1
2
3
4
20180424_BZ14_control STAligner ARI: -0.008
spatial2
spatial1
1
2
3
4

**Fig. S26 Results of STAligner on Mouse medial prefrontal cortex by STARmap.**

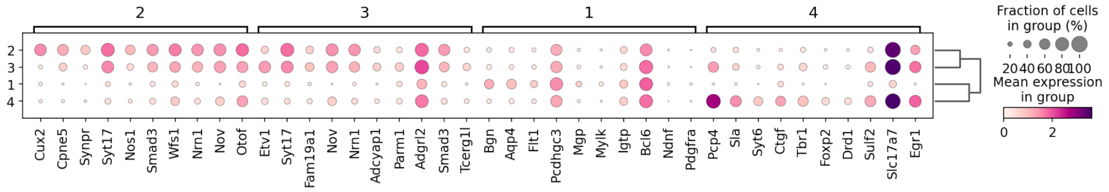
2
3
1
4
Cux2
Cpne5
Synpr
Syt17
Nos1
Smad3
Wfs1
Nrn1
Nov
Otof
Etv1
Syt17
Fam19a1
Nov
Nrn1
Adcyap1
Parm1
Adgrl2
Smad3
Tcerg1l
Bgn
Aqp4
Flt1
Pcdhgc3
Mgp
Mylk
Igtp
Bcl6
Ndnf
Pdgfra
Pcp4
Sla
Syt6
Ctgf
Tbr1
Foxp2
Drd1
Sulf2
Slc17a7
Egr1
Fraction of cells in group (%)
20 40 60 80 100
Mean expression in group
0
2

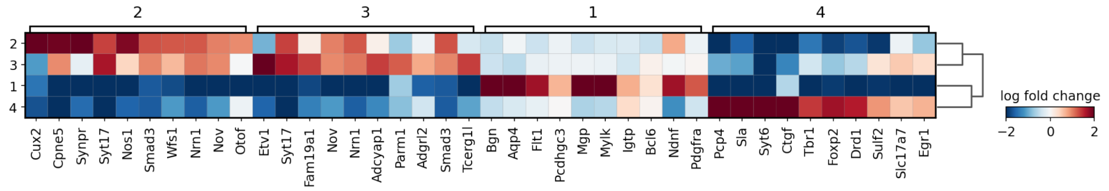
2
3
1
4
Cux2
Cpne5
Synpr
Syt17
Nos1
Smad3
Wfs1
Nrn1
Nov
Otof
Etv1
Syt17
Fam19a1
Nov
Nrn1
Adcyap1
Parm1
Adgrl2
Smad3
Tcerg1l
Bgn
Aqp4
Flt1
Pcdhgc3
Mgp
Mylk
Igtp
Bcl6
Ndnf
Pdgfra
Pcp4
Sla
Syt6
Ctgf
Tbr1
Foxp2
Drd1
Sulf2
Slc17a7
Egr1
log fold change
−2
0
2

**Fig. S27 Results of spatially differentially expressed genes analysis.** Top panel: dot plot of the average gene expression levels and proportions of expressed genes in each domain; Bottom panel: matrix plot of the average log fold changes of the gene expression levels in each domain when referring to the rest domains.

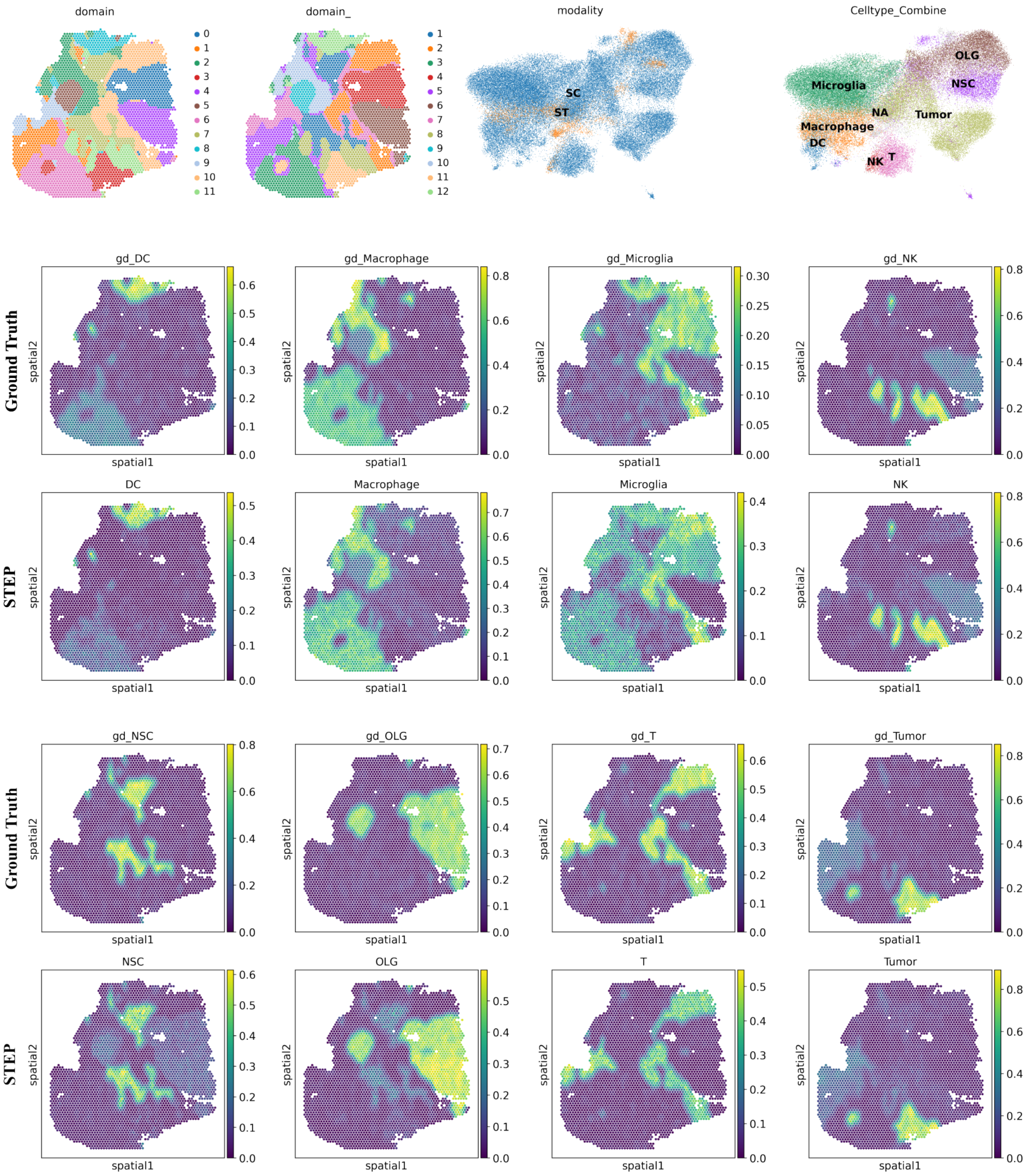
domain
0
1
2
3
4
5
6
7
8
9
10
11
domain_
1
2
3
4
5
6
7
8
9
10
11
12
modality
SC
ST
Celltype_Combine
OLG
Microglia
NSC
NA
Tumor
Macrophage
DC
NK
T
Ground Truth
STEP
Ground Truth
STEP
gd_DC
gd_Macrophage
gd_Microglia
gd_NK
DC
Macrophage
Microglia
NK
gd_NSC
gd_OLG
gd_T
gd_Tumor
NSC
OLG
T
Tumor
spatial1
spatial2

**Fig. S28 Simulation study of the cell-type deconvolution together with spatial domain identification.** Top panel: Ground truth of spatial domains and resembled spatial domains by STEP (first two figures). UMAPs of the co-embedding result by STEP, colored by modalities and cell-types, respectively; Bottom panel: Ground truth cell-type composition vs estimated cell-type composition by STEP in simulated SRT data.

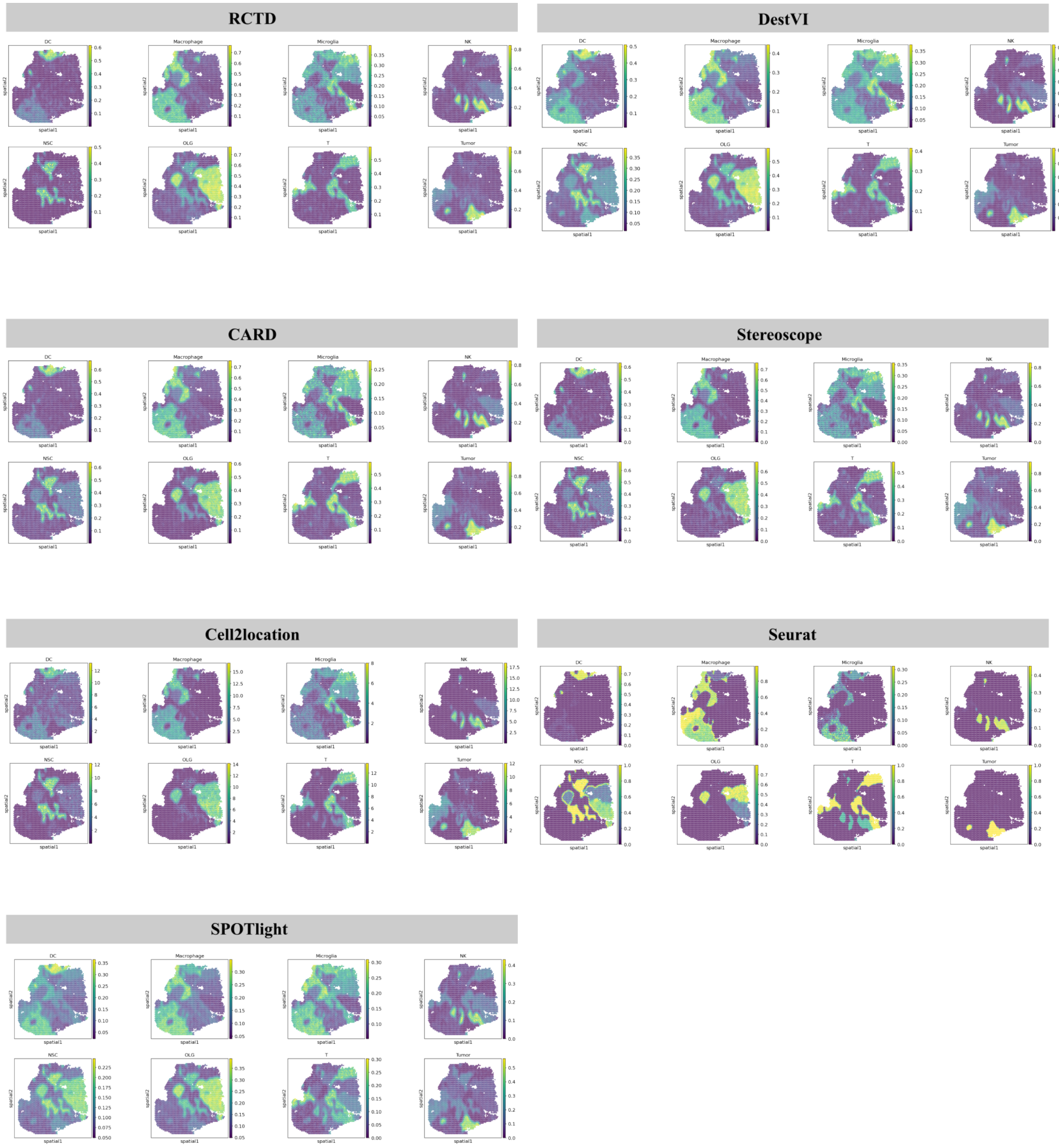
RCTD
DestVI
CARD
Stereoscope
Cell2location
Seurat
SPOTlight
DC
Macrophage
Microglia
NK
NSC
OLG
T
Tumor
spatial1
spatial2

**Fig. S29 Results of compared methods (RCTD, CARD, Cell2location, SPOTlight, DestVI, Stereoscope and Seurat) on the simulation study.**

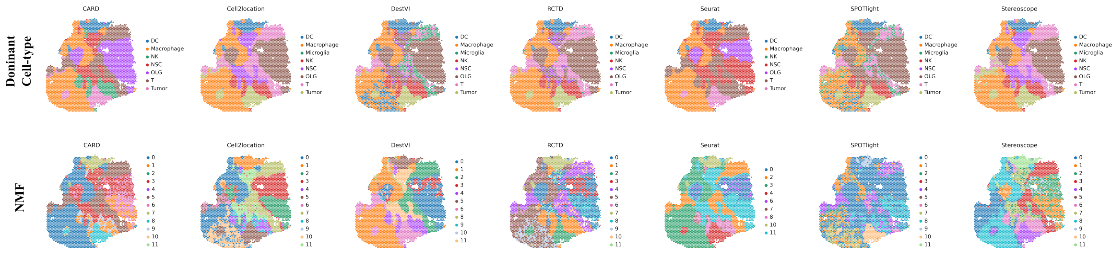
Dominant Cell-type
NMF
CARD
Cell2location
DestVI
RCTD
Seurat
SPOTlight
Stereoscope
DC
Macrophage
Microglia
NK
NSC
OLG
T
Tumor

**Fig. S30 Spatial domains inferred from estimated cell-type composition obtained by each compared methods through two ways: 1) Dominant cell type; 2) Non-negative Matrix Factorization (NMF).** Top panel: Dominant cell type in each spot; Bottom: Argmax of NMF coefficients of the components in each spot.

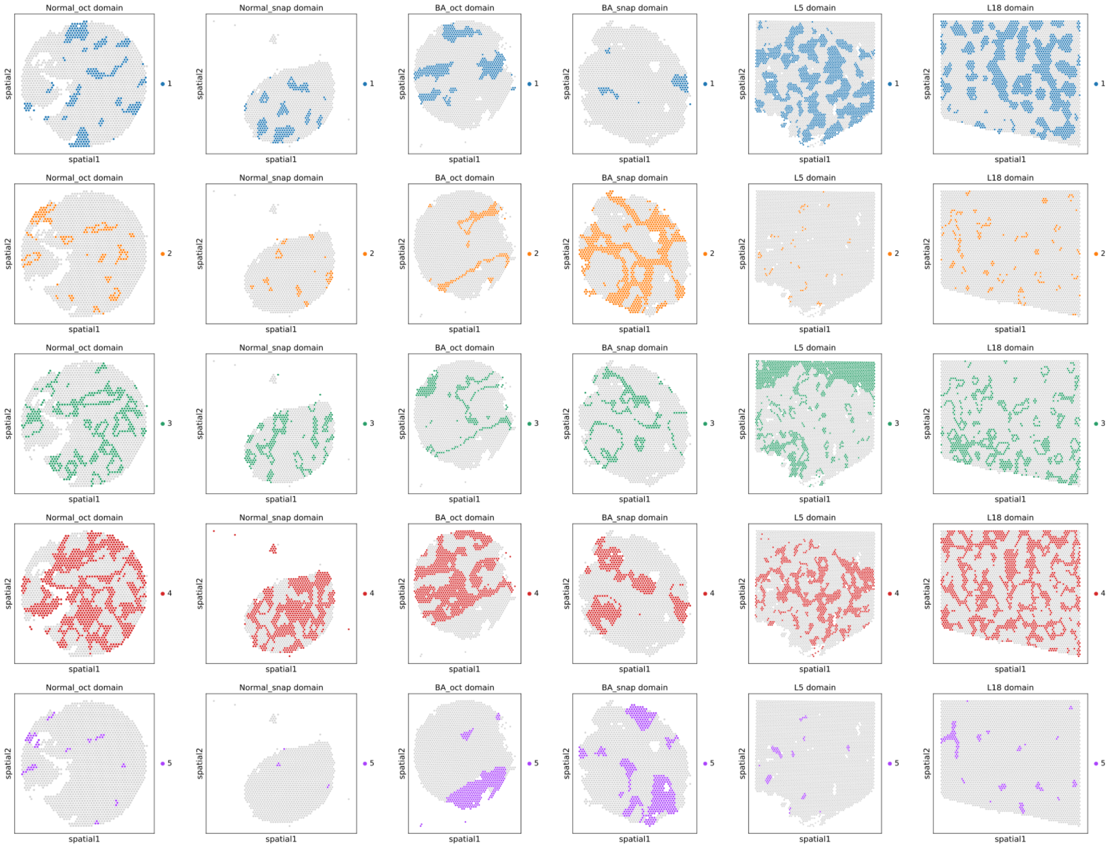
Normal_oct domain
Normal_snap domain
BA_oct domain
BA_snap domain
L5 domain
L18 domain
spatial1
spatial2
1
2
3
4
5

**Fig. S31 Spatial plots of each domain obtained from STEP in each section.** Each row represents the results across sections of a method; each column represents all identified spatial domains of a liver section.

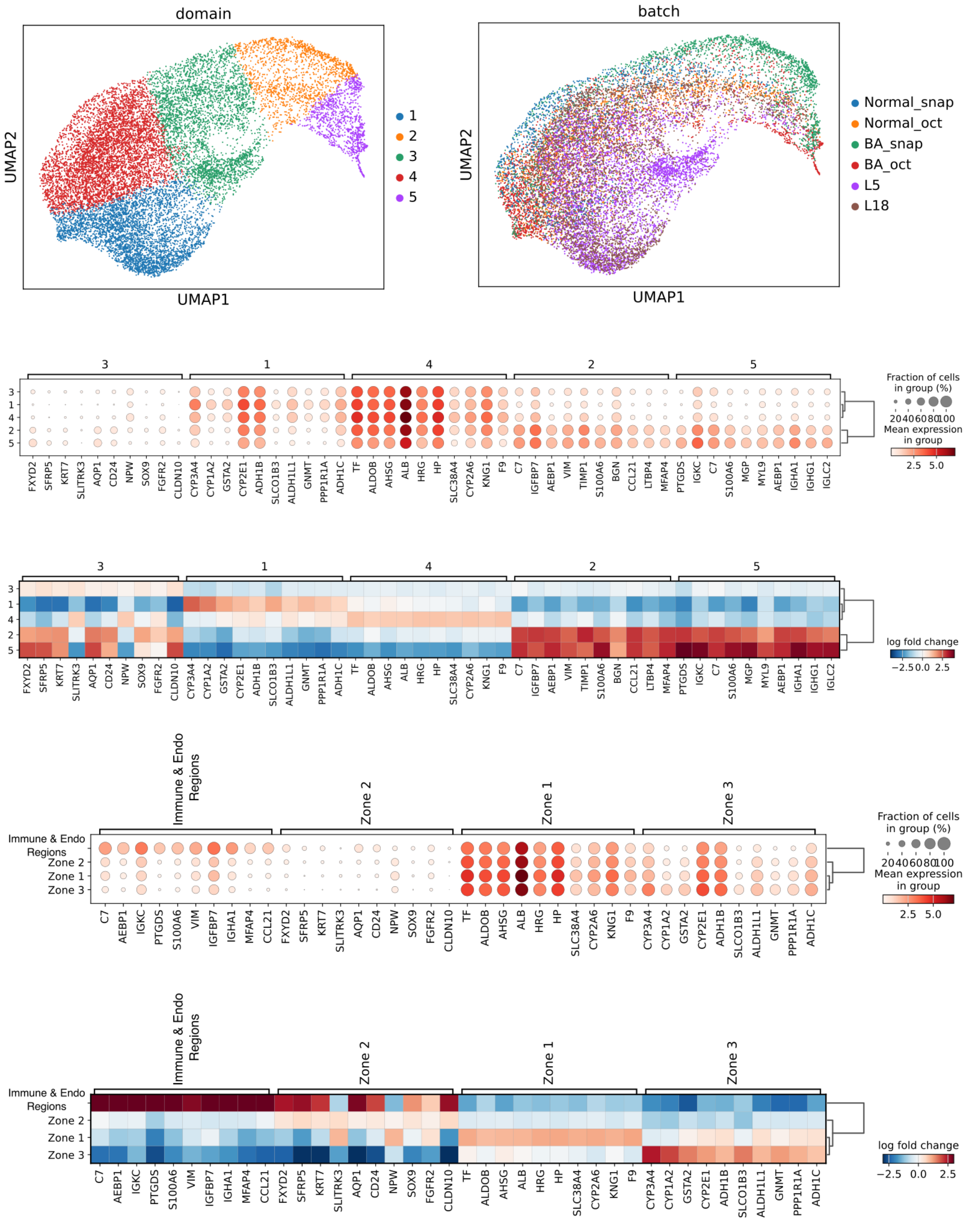
domain
UMAP2
UMAP1
1
2
3
4
5
batch
Normal_snap
Normal_oct
BA_snap
BA_oct
L5
L18
Fraction of cells in group (%)
20 40 60 80 100
Mean expression in group
2.5 5.0
log fold change
−2.5 0.0 2.5
FXYD2 SFRP5 KRT7 SLITRK3 AQP1 CD24 NPW SOX9 FGFR2 CLDN10
CYP3A4 CYP1A2 GSTA2 CYP2E1 ADH1B SLCO1B3 ALDH1L1 GNMT PPP1R1A ADH1C
TF ALDOB AHSG ALB HRG HP SLC38A4 CYP2A6 KNG1 F9
C7 IGFBP7 AEBP1 VIM TIMP1 S100A6 BGN CCL21 LTBP4 MFAP4
PTGDS IGKC C7 S100A6 MGP MYL9 AEBP1 IGHA1 IGHG1 IGLC2
Immune & Endo Regions
Zone 2
Zone 1
Zone 3
C7 AEBP1 IGKC PTGDS S100A6 VIM IGFBP7 IGHA1 MFAP4 CCL21

**Fig. S32 Results of differentially expressed genes analyses on the outputs of STEP.** The first row consists of two UMAP plots where the UMAP is generated by embeddings of STEP and plots are colored by spatial domains identified by STEP and section indicators, respectively; the remnant of the figure consists of two dotplots and two matrixplots, displaying the expression levels of the DEGs which are calculated based on spatial domains and zonation, respectively.

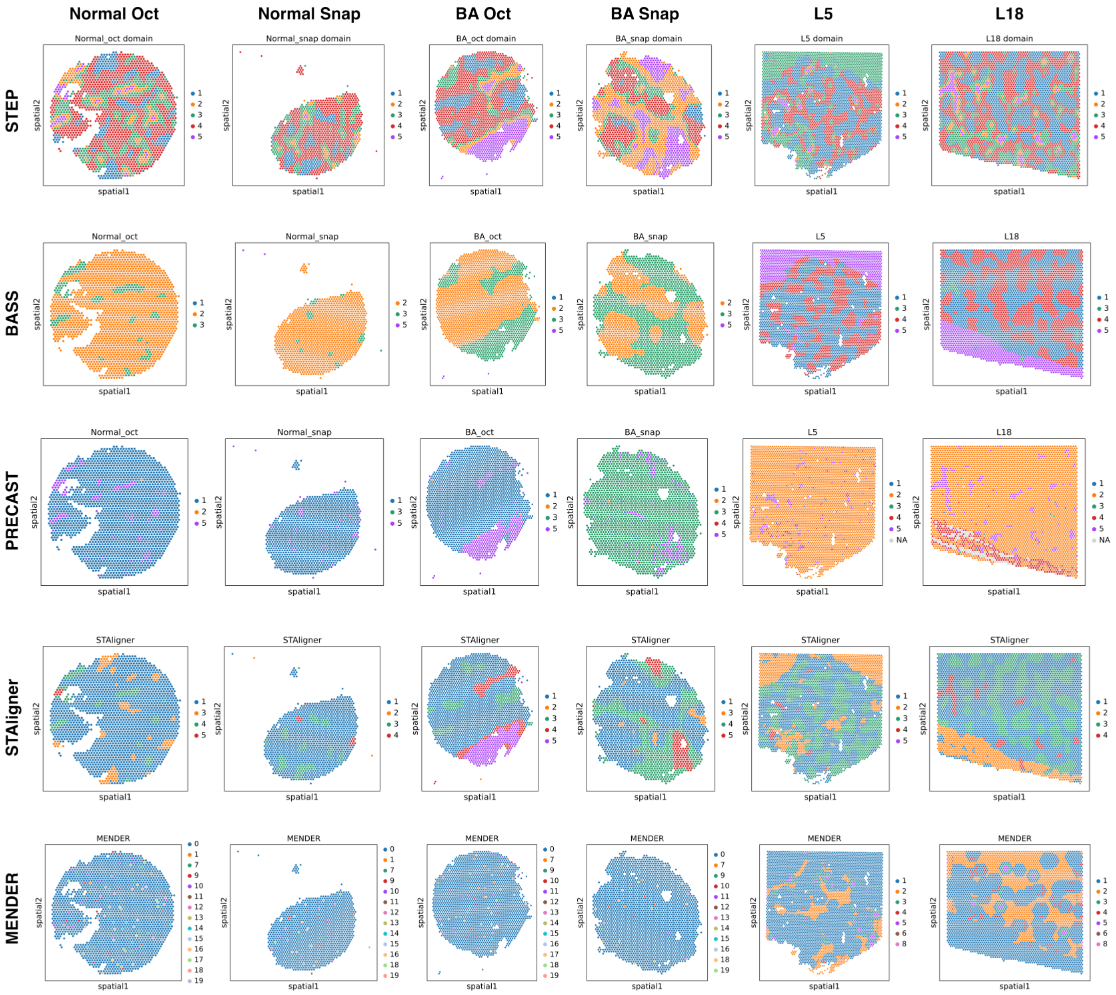
Normal Oct
Normal Snap
BA Oct
BA Snap
L5
L18
STEP
BASS
PRECAST
STAligner
MENDER
Normal_oct domain
Normal_snap domain
BA_oct domain
BA_snap domain
L5 domain
L18 domain
Normal_oct
Normal_snap
BA_oct
BA_snap
STAligner
spatial1
spatial2

**Fig. S33 Unified spatial domains generated by each compared method in zonation task of multiple non-contiguous liver sections.** Each row represents the results across sections of a method; each column represents the results of a liver section across methods.

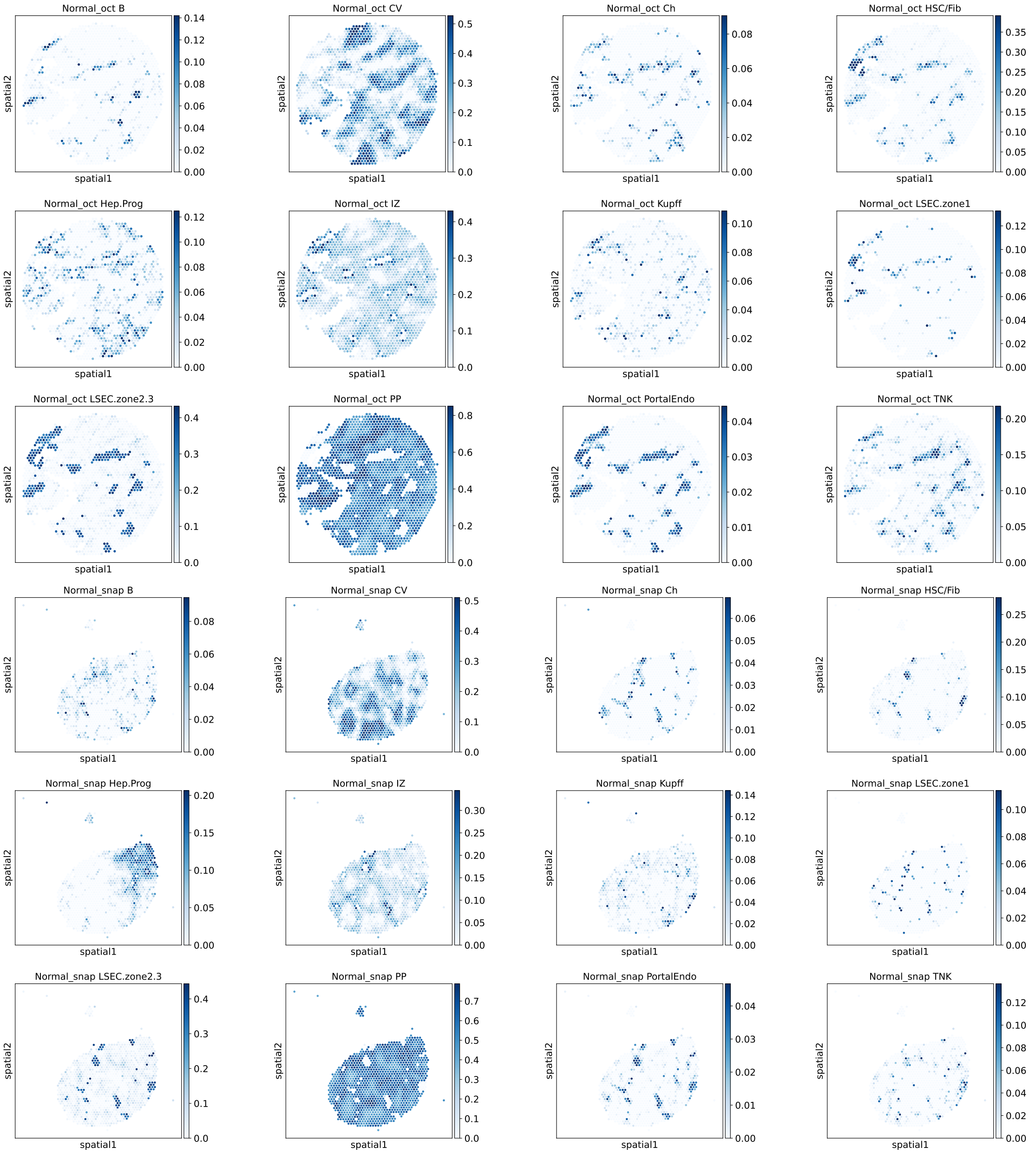
Normal_oct B
Normal_oct CV
Normal_oct Ch
Normal_oct HSC/Fib
Normal_oct Hep.Prog
Normal_oct IZ
Normal_oct Kupff
Normal_oct LSEC.zone1
Normal_oct LSEC.zone2.3
Normal_oct PP
Normal_oct PortalEndo
Normal_oct TNK
Normal_snap B
Normal_snap CV
Normal_snap Ch
Normal_snap HSC/Fib
Normal_snap Hep.Prog
Normal_snap IZ
Normal_snap Kupff
Normal_snap LSEC.zone1
Normal_snap LSEC.zone2.3
Normal_snap PP
Normal_snap PortalEndo
Normal_snap TNK
spatial1
spatial2

**Fig. S34 Spatial plots of multi-section cell-type deconvolution results of Normal Oct and Snap sections.** Colorbar represents the corresponding cell-type abundance. All values are under the same scale and be directly compared with each other, including the following two figures.

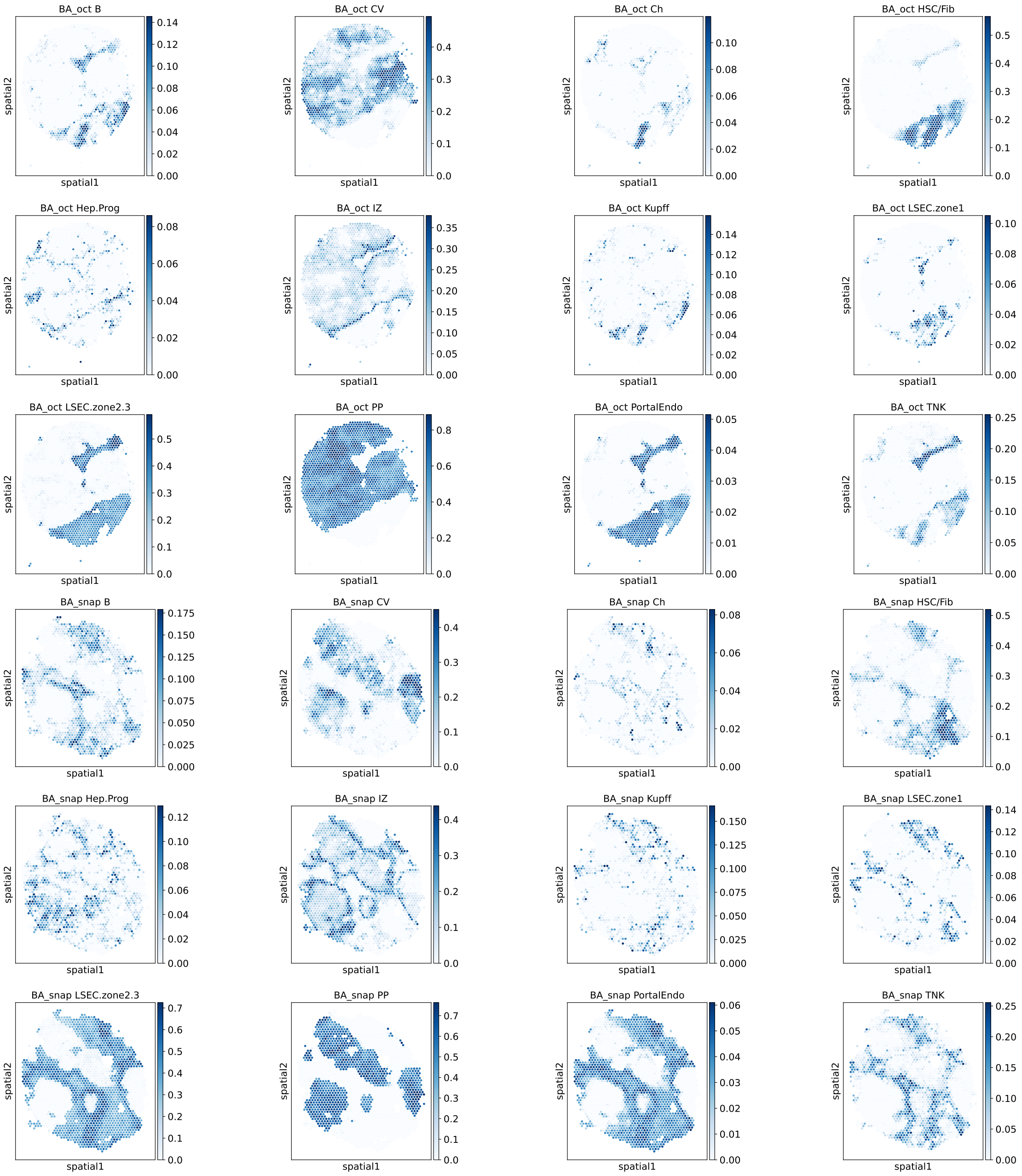

BA_oct B
BA_oct CV
BA_oct Ch
BA_oct HSC/Fib
BA_oct Hep.Prog
BA_oct IZ
BA_oct Kupff
BA_oct LSEC.zone1
BA_oct LSEC.zone2.3
BA_oct PP
BA_oct PortalEndo
BA_oct TNK
BA_snap B
BA_snap CV
BA_snap Ch
BA_snap HSC/Fib
BA_snap Hep.Prog
BA_snap IZ
BA_snap Kupff
BA_snap LSEC.zone1
BA_snap LSEC.zone2.3
BA_snap PP
BA_snap PortalEndo
BA_snap TNK
spatial1
spatial2

**Fig. S35 Spatial plots of multi-section cell-type deconvolution results of BA Oct and Snap sections.** Colorbar represents the corresponding cell-type abundance. All values are under the same scale and be directly compared with each other.

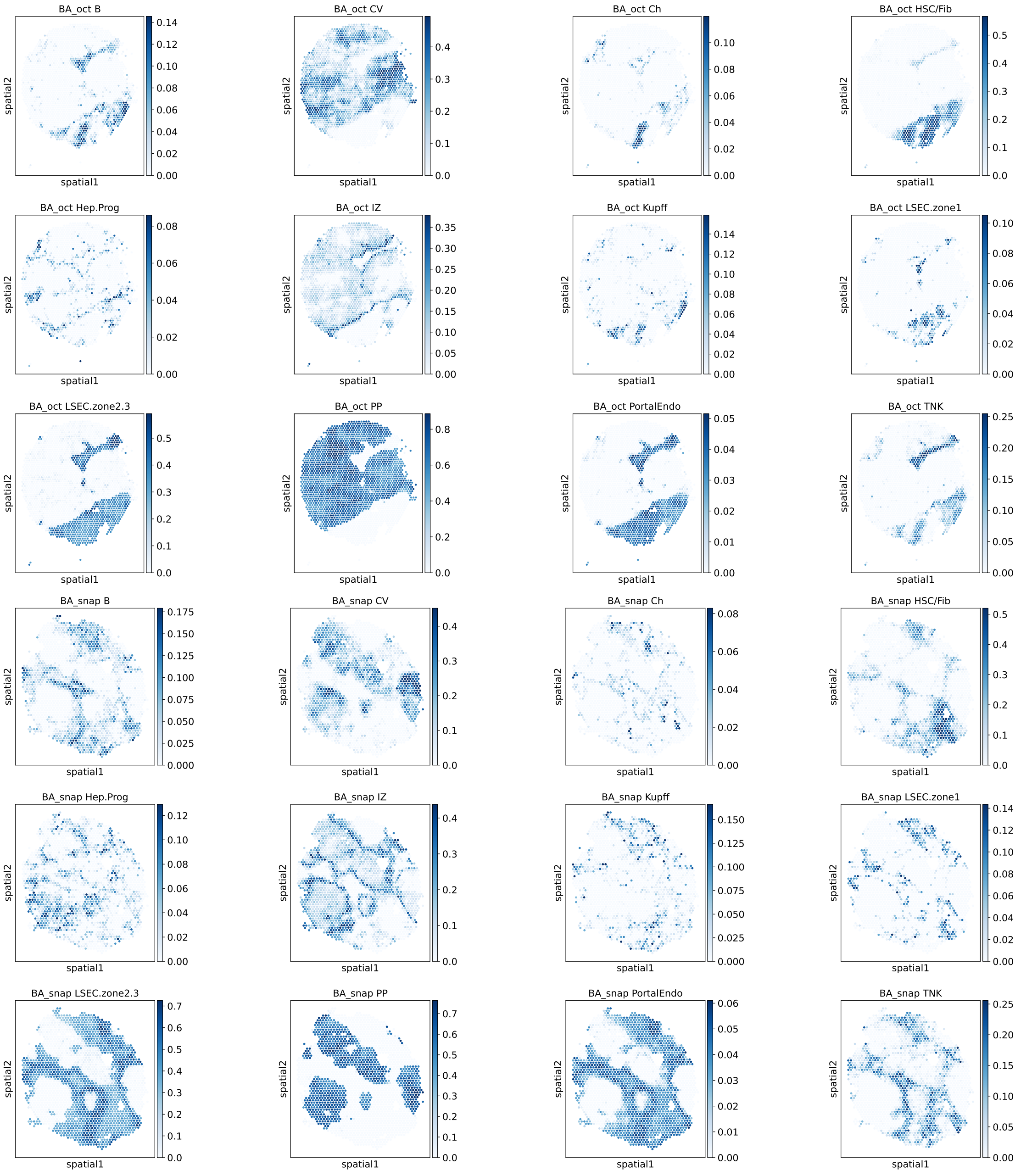
BA_oct B
BA_oct CV
BA_oct Ch
BA_oct HSC/Fib
BA_oct Hep.Prog
BA_oct IZ
BA_oct Kupff
BA_oct LSEC.zone1
BA_oct LSEC.zone2.3
BA_oct PP
BA_oct PortalEndo
BA_oct TNK
BA_snap B
BA_snap CV
BA_snap Ch
BA_snap HSC/Fib
BA_snap Hep.Prog
BA_snap IZ
BA_snap Kupff
BA_snap LSEC.zone1
BA_snap LSEC.zone2.3
BA_snap PP
BA_snap PortalEndo
BA_snap TNK
spatial1
spatial2

**Fig. S36 Spatial plots of multi-section cell-type deconvolution results of BA Oct and Snap sections.** Colorbar represents the corresponding cell-type abundance. All values are under the same scale and be directly compared with each other.

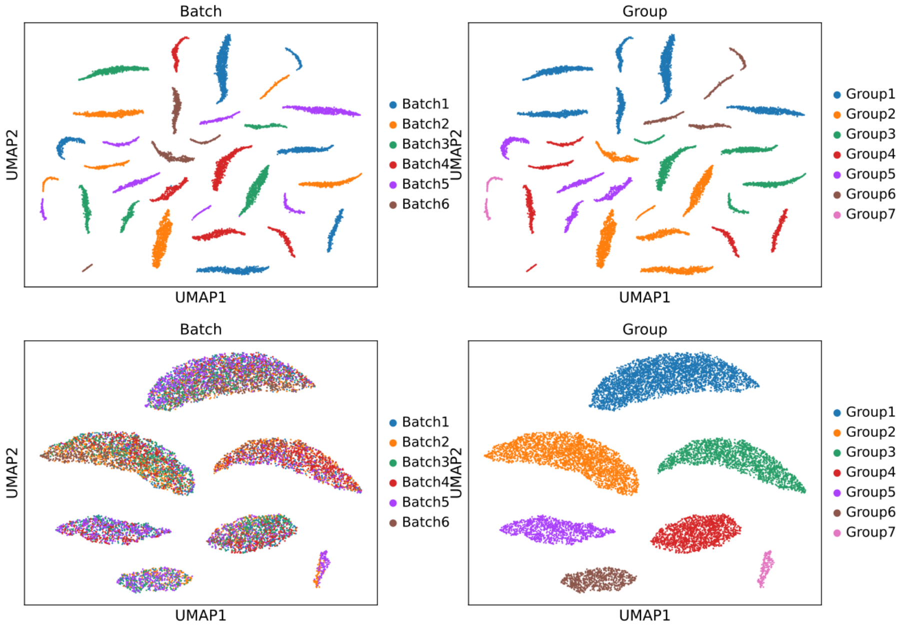

| | Bio conservation | | | | | | | Batch correction | | | | | Aggregate score | | |
|---|---|---|---|---|---|---|---|---|---|---|---|---|---|---|---|
| **Method** | Isolated labels | Leiden NMI | Leiden ARI | KMeans NMI | KMeans ARI | Silhouette label | cLISI | Silhouette batch | iLISI | KBET | Graph connectivity | PCR comparison | Batch correction | Bio conservation | Total |
| **X_pca** | 0.57 | 0.77 | 0.49 | 0.96 | 0.96 | 0.58 | 1.00 | 0.86 | 0.00 | 0.00 | 0.85 | 0.00 | 0.34 | 0.76 | 0.59 |
| **X_pca_uncorrected** | 0.67 | 0.69 | 0.30 | 0.19 | 0.04 | 0.45 | 1.00 | 0.75 | 0.00 | 0.00 | 0.47 | 0.00 | 0.25 | 0.48 | 0.38 |

**Fig. S37 Benchmarking embeddings generated by PCA on raw gene expression data and corrected gene expression data from Simulation Data 1.** Top row: UMAP of PCs from raw gene expression data; bottom row: UMAP of PCs from corrected gene expression data; left column: colored by batch ids; right column: colored by cell types.

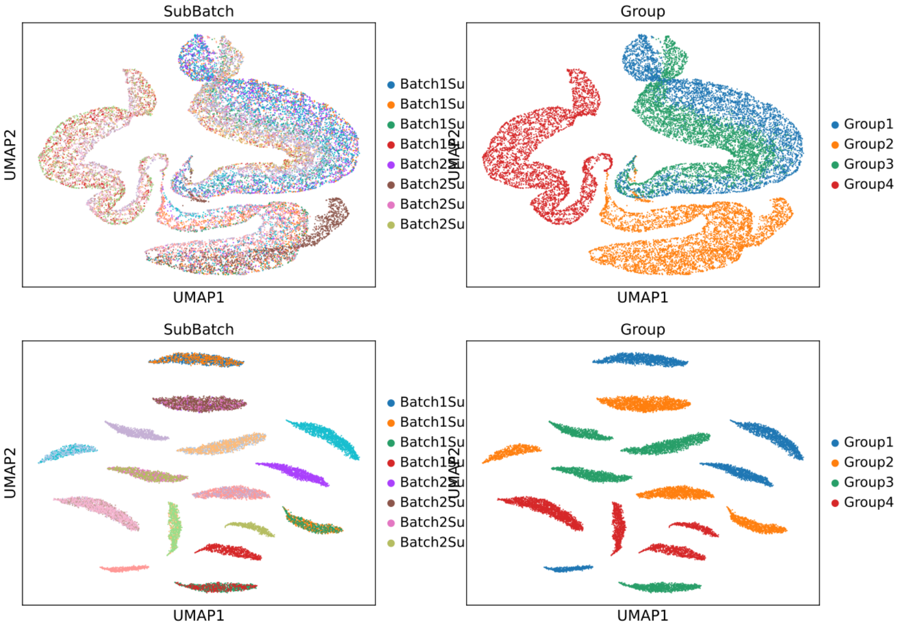

| Method | Bio conservation | | | | | | | Batch correction | | | | | Aggregate score | | |
|---|---|---|---|---|---|---|---|---|---|---|---|---|---|---|---|
| | Isolated labels | Leiden NMI | Leiden ARI | KMeans NMI | KMeans ARI | Silhouette label | cLISI | Silhouette batch | iLISI | KBET | Graph connectivity | PCR comparison | Batch correction | Bio conservation | Total |
| **X_pca_uncorrected** | 0.55 | 0.68 | 0.37 | 0.14 | 0.07 | 0.56 | 1.00 | 0.85 | 0.04 | 0.00 | 0.37 | 0.02 | 0.26 | 0.48 | 0.39 |
| **X_pca** | 0.57 | 0.60 | 0.25 | 0.47 | 0.32 | 0.56 | 1.00 | 0.51 | 0.00 | 0.00 | 0.28 | 0.00 | 0.16 | 0.54 | 0.39 |

**Fig. S38 Benchmarking embeddings generated by PCA on raw gene expression data and corrected gene expression data from Simulation Data 2.** Top row: UMAP of PCs from raw gene expression data; bottom row: UMAP of PCs from corrected gene expression data; left column: colored by batch ids; right column: colored by cell types.

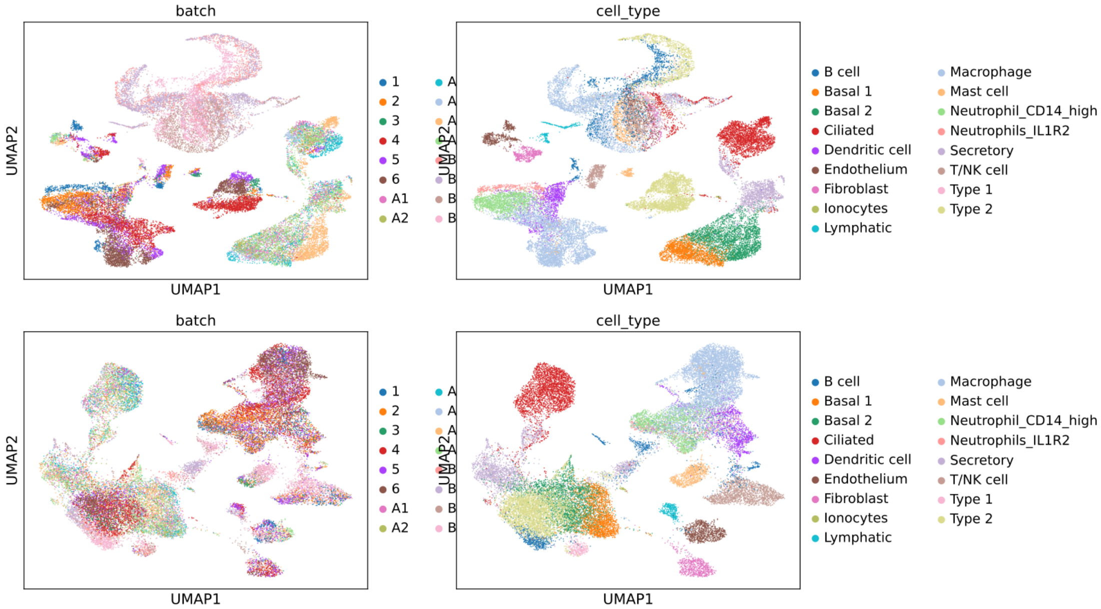

| | Bio conservation | | | | | | | Batch correction | | | | | Aggregate score | | |
|---|---|---|---|---|---|---|---|---|---|---|---|---|---|---|---|
| **Method** | Isolated labels | Leiden NMI | Leiden ARI | KMeans NMI | KMeans ARI | Silhouette label | cLISI | Silhouette batch | iLISI | KBET | Graph connectivity | PCR comparison | Batch correction | Bio conservation | Total |
| **X_pca_uncorrected** | 0.38 | 0.69 | 0.50 | 0.46 | 0.26 | 0.42 | 1.00 | 0.81 | 0.03 | 0.24 | 0.77 | 0.81 | 0.53 | 0.53 | 0.53 |
| **X_pca** | 0.66 | 0.73 | 0.41 | 0.66 | 0.43 | 0.55 | 1.00 | 0.88 | 0.00 | 0.22 | 0.74 | 0.00 | 0.37 | 0.63 | 0.53 |

**Fig. S39 Benchmarking embeddings generated by PCA on raw gene expression data and corrected gene expression data from Human Lung atlas.** Top row: UMAP of PCs from raw gene expression data; bottom row: UMAP of PCs from corrected gene expression data; left column: colored by batch ids; right column: colored by cell types.

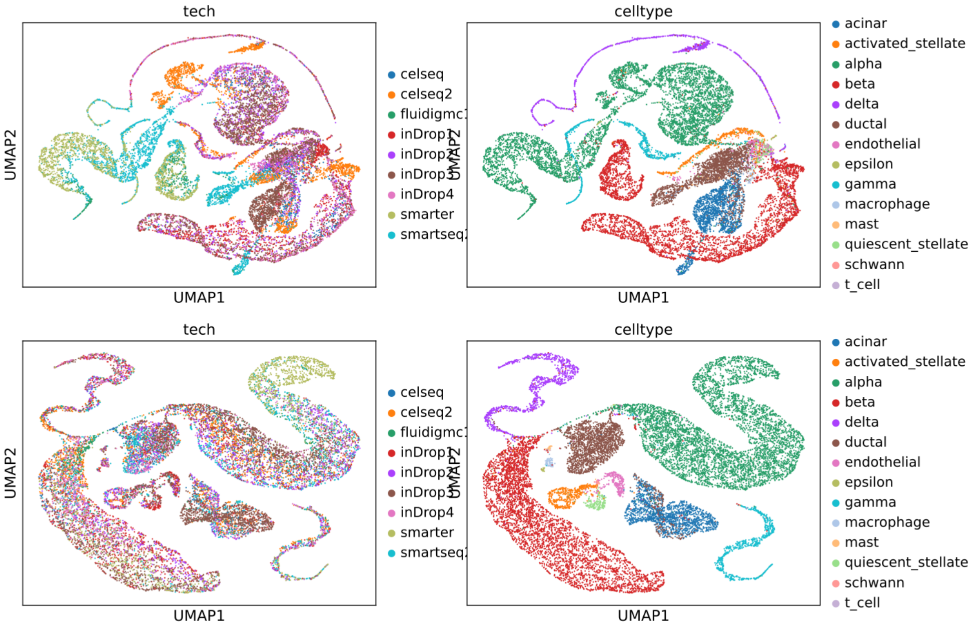

| Method | Bio conservation | | | | | | | Batch correction | | | | | Aggregate score | | |
|---|---|---|---|---|---|---|---|---|---|---|---|---|---|---|---|
| | Isolated labels | Leiden NMI | Leiden ARI | KMeans NMI | KMeans ARI | Silhouette label | cLISI | Silhouette batch | iLISI | KBET | Graph connectivity | PCR comparison | Batch correction | Bio conservation | Total |
| **X_pca** | 0.67 | 0.66 | 0.41 | 0.56 | 0.36 | 0.53 | 1.00 | 0.85 | 0.00 | 0.19 | 0.81 | 0.00 | 0.37 | 0.60 | 0.51 |
| **X_pca_uncorrected** | 0.99 | 0.63 | 0.34 | 0.17 | -0.04 | 0.14 | 1.00 | 0.67 | 0.11 | 0.16 | 0.63 | 0.19 | 0.35 | 0.46 | 0.42 |

**Fig. S40 Benchmarking embeddings generated by PCA on raw gene expression data and corrected gene expression data from Human Pancreas data.** Top row: UMAP of PCs from raw gene expression data; bottom row: UMAP of PCs from corrected gene expression data; left column: colored by batch ids; right column: colored by cell types.

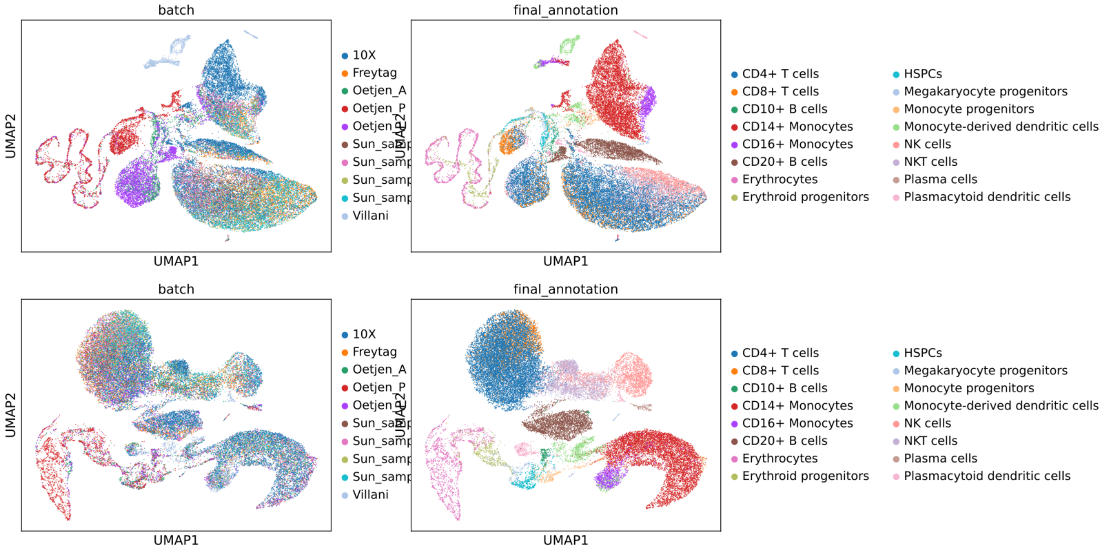

| | Bio conservation | | | | | | | Batch correction | | | | | Aggregate score | | |
|---|---|---|---|---|---|---|---|---|---|---|---|---|---|---|---|
| **Method** | Isolated labels | Leiden NMI | Leiden ARI | KMeans NMI | KMeans ARI | Silhouette label | cLISI | Silhouette batch | iLISI | KBET | Graph connectivity | PCR comparison | Batch correction | Bio conservation | Total |
| **X_pca** | 0.62 | 0.68 | 0.42 | 0.62 | 0.41 | 0.56 | 1.00 | 0.84 | 0.02 | 0.12 | 0.64 | 0.00 | 0.32 | 0.62 | 0.50 |
| **X_pca_uncorrected** | 0.42 | 0.57 | 0.32 | 0.22 | 0.06 | 0.41 | 0.98 | 0.79 | 0.08 | 0.14 | 0.73 | 0.00 | 0.35 | 0.43 | 0.39 |

**Fig. S41 Benchmarking embeddings generated by PCA on raw gene expression data and corrected gene expression data from Human Immune cell atlas.** Top row: UMAP of PCs from raw gene expression data; bottom row: UMAP of PCs from corrected gene expression data; left column: colored by batch ids; right column: colored by cell types.

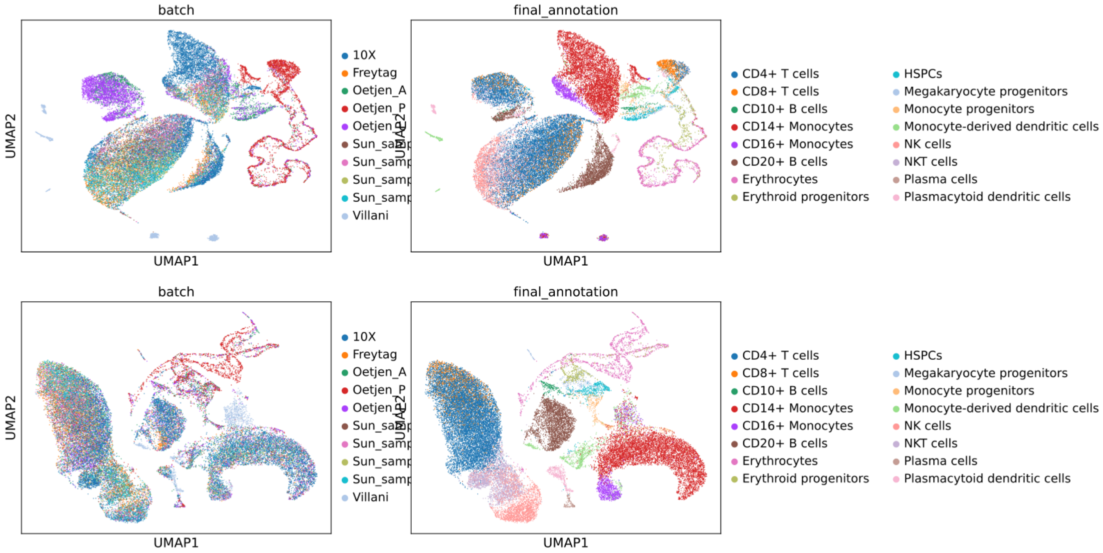

| | Bio conservation | | | | | | | Batch correction | | | | | Aggregate score | | |
|---|---|---|---|---|---|---|---|---|---|---|---|---|---|---|---|
| **Method** | Isolated labels | Leiden NMI | Leiden ARI | KMeans NMI | KMeans ARI | Silhouette label | cLISI | Silhouette batch | iLISI | KBET | Graph connectivity | PCR comparison | Batch correction | Bio conservation | Total |
| **X_anchord** | 0.74 | 0.98 | 0.99 | 0.82 | 0.50 | 0.64 | 1.00 | 0.95 | 0.30 | 0.83 | 0.89 | 0.84 | 0.76 | 0.81 | 0.79 |
| **X_rep** | 0.62 | 0.78 | 0.72 | 0.69 | 0.42 | 0.57 | 1.00 | 0.89 | 0.14 | 0.18 | 0.88 | 0.85 | 0.59 | 0.69 | 0.65 |
| **X_pca** | 0.62 | 0.68 | 0.42 | 0.62 | 0.41 | 0.56 | 1.00 | 0.84 | 0.02 | 0.13 | 0.64 | 0.00 | 0.32 | 0.62 | 0.50 |
| **X_pca_uncorrected** | 0.42 | 0.58 | 0.44 | 0.19 | 0.05 | 0.48 | 0.98 | 0.78 | 0.07 | 0.14 | 0.75 | 0.46 | 0.44 | 0.45 | 0.45 |

**Fig. S42 Benchmarking embeddings generated by PCA on raw gene expression data and corrected gene expression data from Human & Mouse Immune cell atlas.** Top row: UMAP of PCs from raw gene expression data; bottom row: UMAP of PCs from corrected gene expression data; left column: colored by batch ids; right column: colored by cell types.

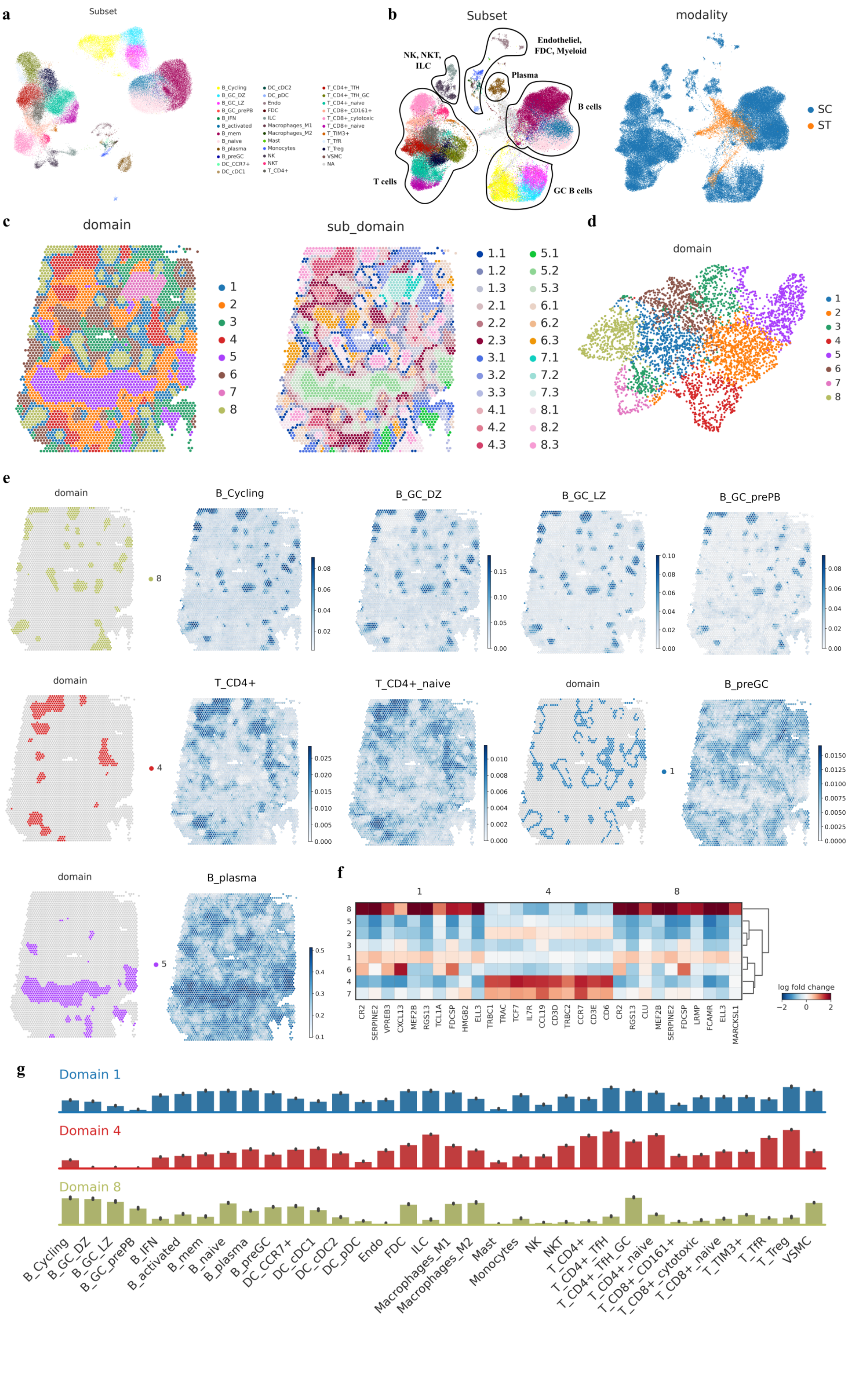
a
Subset
b
Subset
modality
Endotheliel, FDC, Myeloid
NK, NKT, ILC
Plasma
B cells
T cells
GC B cells
SC
ST
c
domain
sub_domain
d
domain
e
f
log fold change
g
Domain 1
Domain 4
Domain 8

**Fig. S43 Analysis of the spatial domain identification and deconvolution on Human Lymph Node data.** a. UMAP colored by cell-type for the reference scRNA-seq data obtained by the original study; b. left: UMAP colored by cell-type for the co-embedded two modalities obtained by STEP, right: UMAP colored by modality for the co-embedded two modalities obtained by STEP; c. Identified 8 spatial domains and 24 sub-domains by STEP; d. UMAP colored by spatial domain for the sole SRT data obtained by STEP; e. Comparison for the identified spatial domains and deconvolution results; f. Matrix plot for the result of the differential genes analysis based on 3 focused spatial domains; g. Bar plots for the scaled cell-type composition in the 3 focused spatial domains.

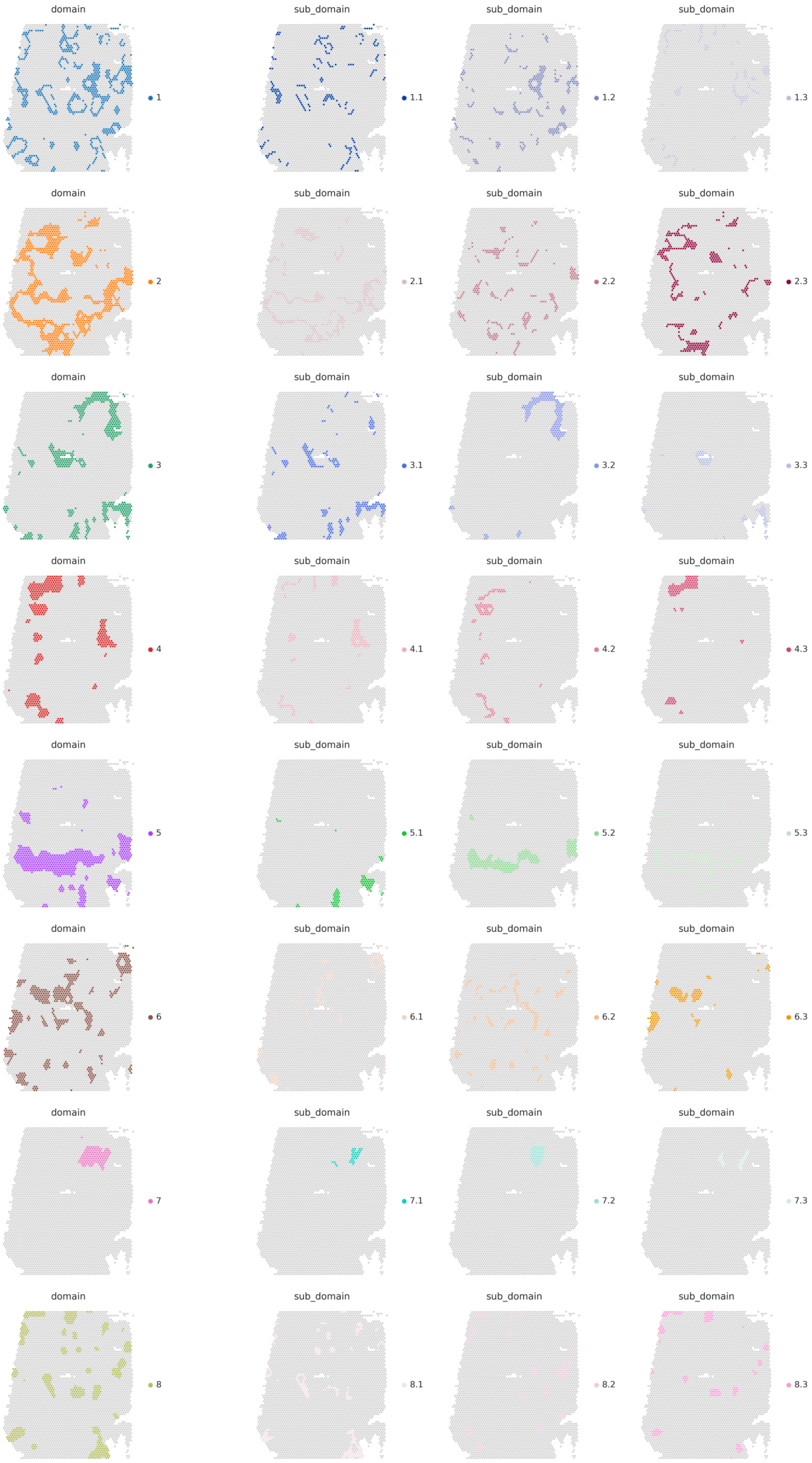
domain
sub_domain
1
1.1
1.2
1.3
2
2.1
2.2
2.3
3
3.1
3.2
3.3
4
4.1
4.2
4.3
5
5.1
5.2
5.3
6
6.1
6.2
6.3
7
7.1
7.2
7.3
8
8.1
8.2
8.3

**Fig. S44 Spatial domains and corresponding sub-domains identified by STEP on Human Lymph Node data by 10x Visium.** Each row represents one domain and the sub-domains identified based on it.

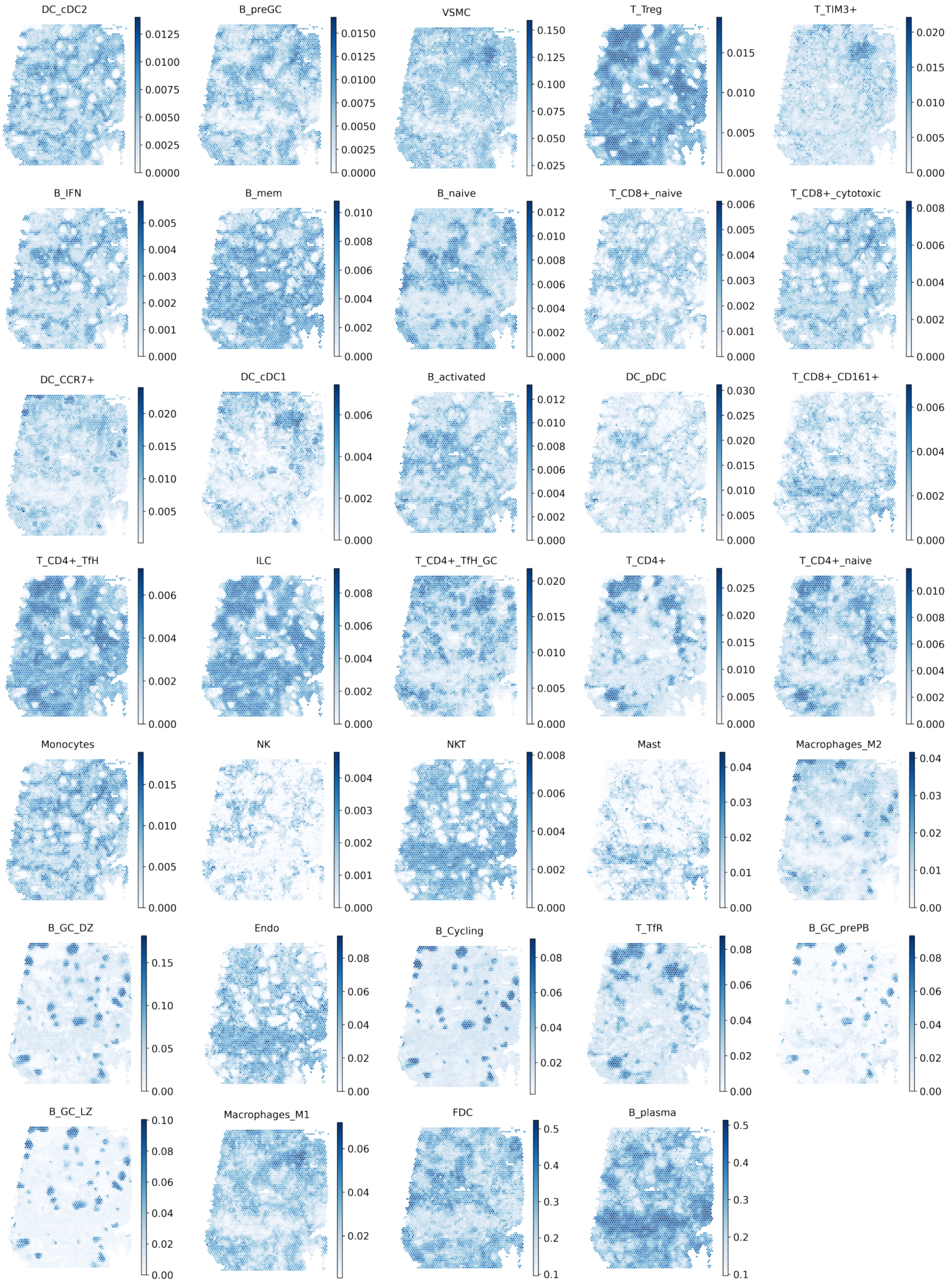
DC_cDC2
0.0125 0.0100 0.0075 0.0050 0.0025 0.0000
B_preGC
0.0150 0.0125 0.0100 0.0075 0.0050 0.0025 0.0000
VSMC
0.150 0.125 0.100 0.075 0.050 0.025
T_Treg
0.015 0.010 0.005 0.000
T_TIM3+
0.020 0.015 0.010 0.005 0.000
B_IFN
0.005 0.004 0.003 0.002 0.001 0.000
B_mem
0.010 0.008 0.006 0.004 0.002 0.000
B_naive
0.012 0.010 0.008 0.006 0.004 0.002 0.000
T_CD8+_naive
0.006 0.005 0.004 0.003 0.002 0.001 0.000
T_CD8+_cytotoxic
0.008 0.006 0.004 0.002 0.000
DC_CCR7+
0.020 0.015 0.010 0.005
DC_cDC1
0.006 0.004 0.002 0.000
B_activated
0.012 0.010 0.008 0.006 0.004 0.002 0.000
DC_pDC
0.030 0.025 0.020 0.015 0.010 0.005 0.000
T_CD8+_CD161+
0.006 0.004 0.002 0.000
T_CD4+_TfH
0.006 0.004 0.002 0.000
ILC
0.008 0.006 0.004 0.002 0.000
T_CD4+_TfH_GC
0.020 0.015 0.010 0.005 0.000
T_CD4+
0.025 0.020 0.015 0.010 0.005 0.000
T_CD4+_naive
0.010 0.008 0.006 0.004 0.002 0.000
Monocytes
0.015 0.010 0.005 0.000
NK
0.004 0.003 0.002 0.001 0.000
NKT
0.008 0.006 0.004 0.002 0.000
Mast
0.04 0.03 0.02 0.01 0.00
Macrophages_M2
0.04 0.03 0.02 0.01 0.00
B_GC_DZ
0.15 0.10 0.05 0.00
Endo
0.08 0.06 0.04 0.02 0.00
B_Cycling
0.08 0.06 0.04 0.02
T_TfR
0.08 0.06 0.04 0.02 0.00
B_GC_prePB
0.08 0.06 0.04 0.02 0.00
B_GC_LZ
0.10 0.08 0.06 0.04 0.02 0.00
Macrophages_M1
0.06 0.04 0.02
FDC
0.5 0.4 0.3 0.2 0.1
B_plasma
0.5 0.4 0.3 0.2 0.1

**Fig. S45 Results of cell-type deconvolution, obtained by STEP, on Human Lymph Node by 10x Visium.**

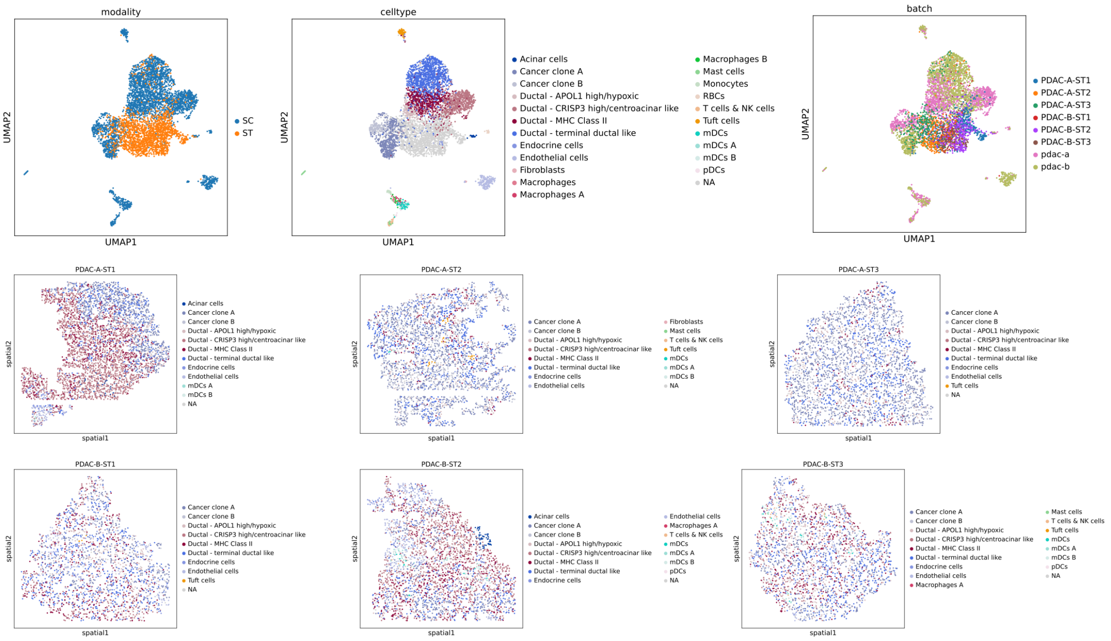
modality
UMAP2
UMAP1
SC
ST
celltype
UMAP2
UMAP1
Acinar cells
Cancer clone A
Cancer clone B
Ductal - APOL1 high/hypoxic
Ductal - CRISP3 high/centroacinar like
Ductal - MHC Class II
Ductal - terminal ductal like
Endocrine cells
Endothelial cells
Fibroblasts
Macrophages
Macrophages A
Macrophages B
Mast cells
Monocytes
RBCs
T cells & NK cells
Tuft cells
mDCs
mDCs A
mDCs B
pDCs
NA
batch
UMAP2
UMAP1
PDAC-A-ST1
PDAC-A-ST2
PDAC-A-ST3
PDAC-B-ST1
PDAC-B-ST2
PDAC-B-ST3
pdac-a
pdac-b
PDAC-A-ST1
spatial2
spatial1
Acinar cells
Cancer clone A
Cancer clone B
Ductal - APOL1 high/hypoxic
Ductal - CRISP3 high/centroacinar like
Ductal - MHC Class II
Ductal - terminal ductal like
Endocrine cells
Endothelial cells
mDCs A
mDCs B
NA
PDAC-A-ST2
spatial2
spatial1
Cancer clone A
Cancer clone B
Ductal - APOL1 high/hypoxic
Ductal - CRISP3 high/centroacinar like
Ductal - MHC Class II
Ductal - terminal ductal like
Endocrine cells
Endothelial cells
Fibroblasts
Mast cells
T cells & NK cells
Tuft cells
mDCs
mDCs A
mDCs B
NA
PDAC-A-ST3
spatial2
spatial1
Cancer clone A
Cancer clone B
Ductal - APOL1 high/hypoxic
Ductal - CRISP3 high/centroacinar like
Ductal - MHC Class II
Ductal - terminal ductal like
Endocrine cells
Endothelial cells
Tuft cells
NA
PDAC-B-ST1
spatial2
spatial1
Cancer clone A
Cancer clone B
Ductal - APOL1 high/hypoxic
Ductal - CRISP3 high/centroacinar like
Ductal - MHC Class II
Ductal - terminal ductal like
Endocrine cells
Endothelial cells
Tuft cells
NA
PDAC-B-ST2
spatial2
spatial1
Acinar cells
Cancer clone A
Cancer clone B
Ductal - APOL1 high/hypoxic
Ductal - CRISP3 high/centroacinar like
Ductal - MHC Class II
Ductal - terminal ductal like
Endocrine cells
Endothelial cells
Macrophages A
T cells & NK cells
mDCs
mDCs A
mDCs B
pDCs
NA
PDAC-B-ST3
spatial2
spatial1
Cancer clone A
Cancer clone B
Ductal - APOL1 high/hypoxic
Ductal - CRISP3 high/centroacinar like
Ductal - MHC Class II
Ductal - terminal ductal like
Endocrine cells
Endothelial cells
Macrophages A
Mast cells
T cells & NK cells
Tuft cells
mDCs
mDCs A
mDCs B
pDCs
NA

**Fig. S46 STEP co-embeds 2 scRNA-seq samples and 6 ST samples and uniformly maps scRNA-seq data to ST samples.** scRNA-seq samples: pdac-a and pdac-b; ST samples: PDAC-A-ST1, PDAC-A-ST2, PDAC-A-ST3, PDAC-B-ST1, PDAC-B-ST2, PDAC-B-ST3. Top panel: UMAP of co-embedding colored by modality, cell-type, sample id, respectively.

#### ST Raw

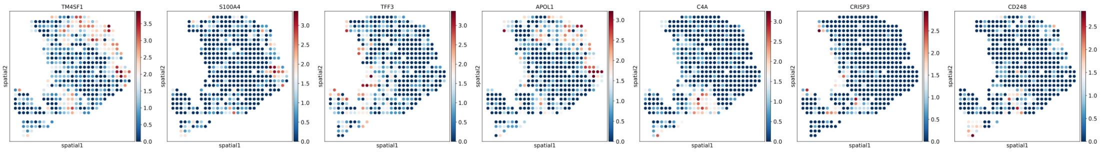

#### ST Raw + SC Mapped

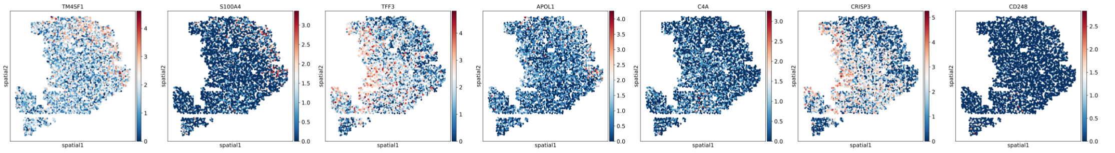

#### SC Mapped

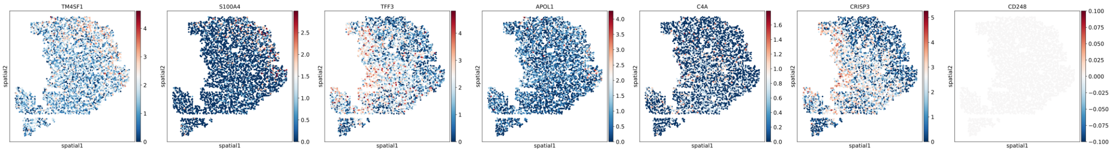

**Fig. S47 Spatial plot of marker genes' expression after scRNA-seq reference mapping for PDAC-A-ST1.** "ST Raw" represents the raw gene expression data of spots in ST sample; "ST Raw + SC Mapped" represents the gene expression of cells mapped on the tissue as well as the original spots; "SC Mapped" represents only the gene expression of cells mapped on the tissue.

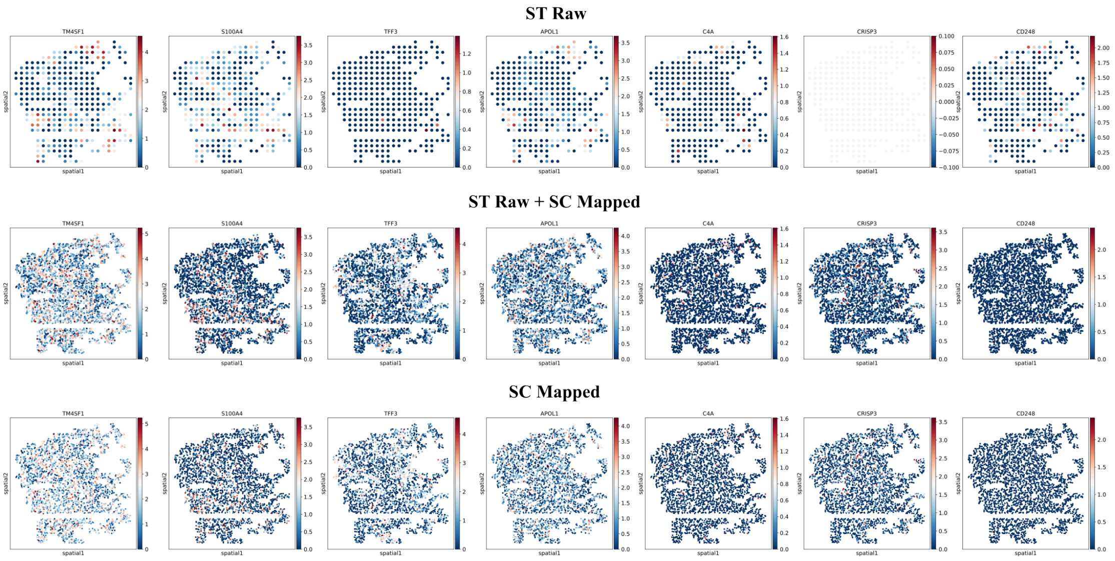

ST Raw
ST Raw + SC Mapped
SC Mapped
TM4SF1
S100A4
TFF3
APOL1
C4A
CRISP3
CD248
spatial1
spatial2

**Fig. S48 Spatial plot of marker genes' expression after scRNA-seq reference mapping for PDAC-A-ST2.** "ST Raw" represents the raw gene expression data of spots in ST sample; "ST Raw + SC Mapped" represents the gene expression of cells mapped on the tissue as well as the original spots; "SC Mapped" represents only the gene expression of cells mapped on the tissue.

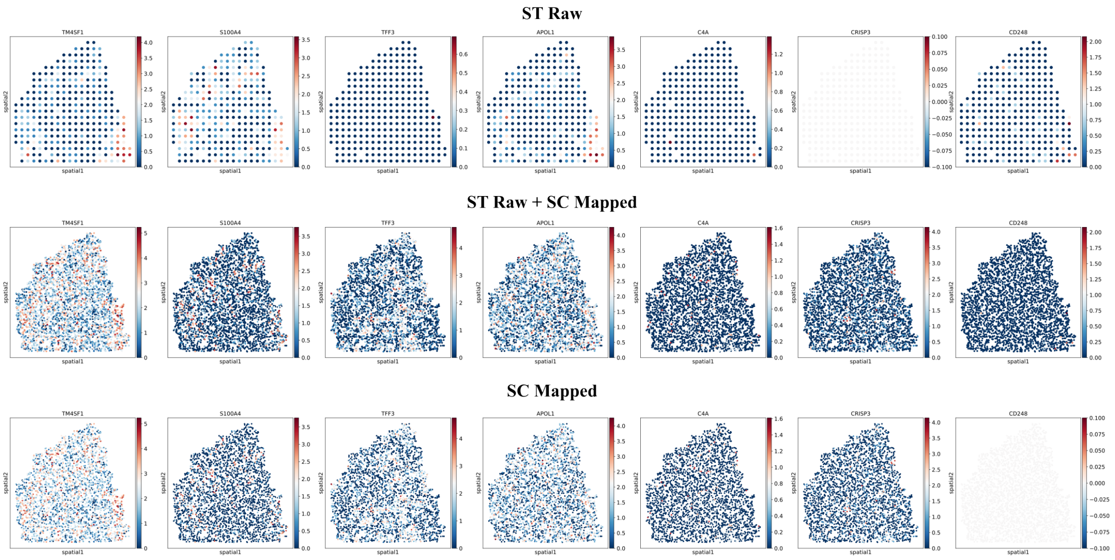

ST Raw
ST Raw + SC Mapped
SC Mapped
TM4SF1
S100A4
TFF3
APOL1
C4A
CRISP3
CD248
spatial1
spatial2

**Fig. S49 Spatial plot of marker genes' expression after scRNA-seq reference mapping for PDAC-A-ST3.** "ST Raw" represents the raw gene expression data of spots in ST sample; "ST Raw + SC Mapped" represents the gene expression of cells mapped on the tissue as well as the original spots; "SC Mapped" represents only the gene expression of cells mapped on the tissue.

#### ST Raw

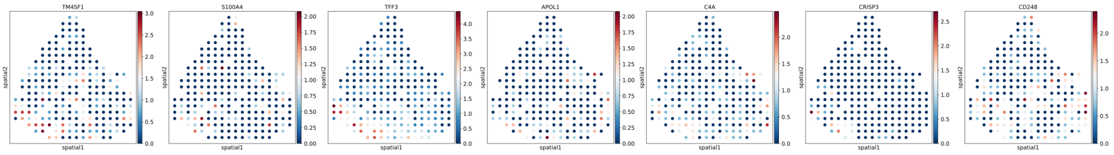

#### ST Raw + SC Mapped

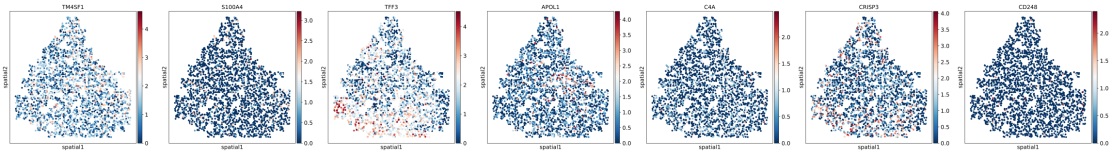

#### SC Mapped

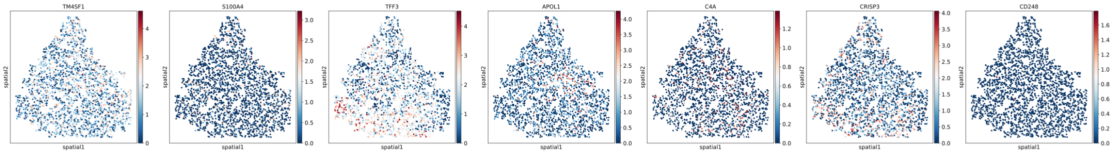

**Fig. S50 Spatial plot of marker genes' expression after scRNA-seq reference mapping for PDAC-B-ST1.** "ST Raw" represents the raw gene expression data of spots in ST sample; "ST Raw + SC Mapped" represents the gene expression of cells mapped on the tissue as well as the original spots; "SC Mapped" represents only the gene expression of cells mapped on the tissue.

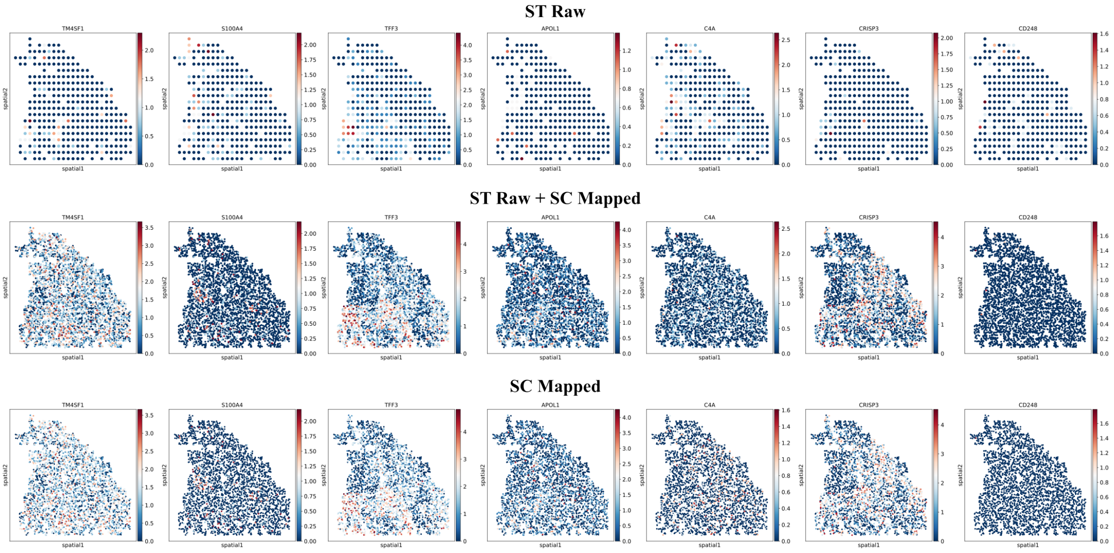
ST Raw
ST Raw + SC Mapped
SC Mapped
TM4SF1
S100A4
TFF3
APOL1
C4A
CRISP3
CD248
spatial1
spatial2

**Fig. S51 Spatial plot of marker genes' expression after scRNA-seq reference mapping for PDAC-B-ST2.** "ST Raw" represents the raw gene expression data of spots in ST sample; "ST Raw + SC Mapped" represents the gene expression of cells mapped on the tissue as well as the original spots; "SC Mapped" represents only the gene expression of cells mapped on the tissue.

#### ST Raw

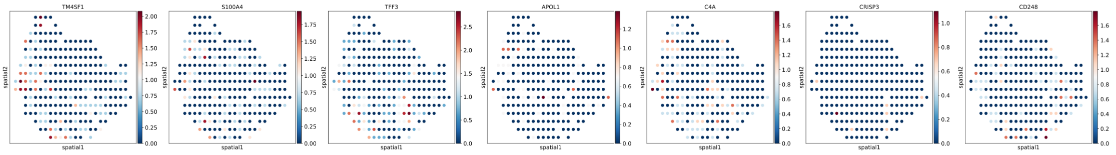

#### ST Raw + SC Mapped

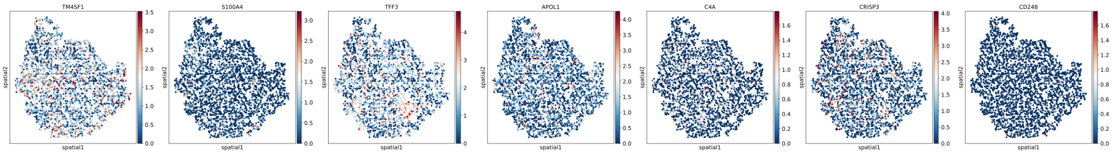

#### SC Mapped

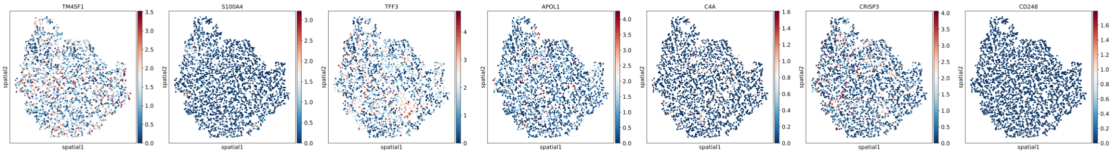

**Fig. S52 Spatial plot of marker genes' expression after scRNA-seq reference mapping for PDAC-B-ST3.** "ST Raw" represents the raw gene expression data of spots in ST sample; "ST Raw + SC Mapped" represents the gene expression of cells mapped on the tissue as well as the original spots; "SC Mapped" represents only the gene expression of cells mapped on the tissue.

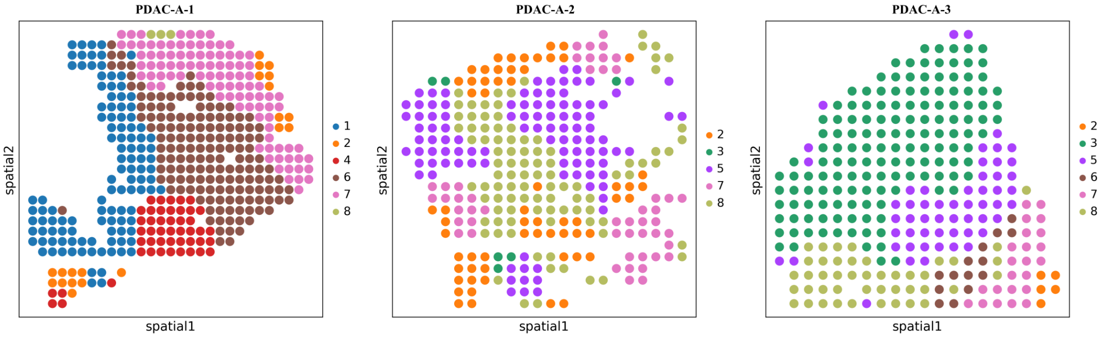
PDAC-A-1
spatial2
spatial1
1
2
4
6
7
8
PDAC-A-2
spatial2
spatial1
2
3
5
7
8
PDAC-A-3
spatial2
spatial1
2
3
5
6
7
8

**Fig. S53 Identified spatial domains across 3 sections in PDAC-A dataset.** Distinct spatial domains across sections were successfully identified.

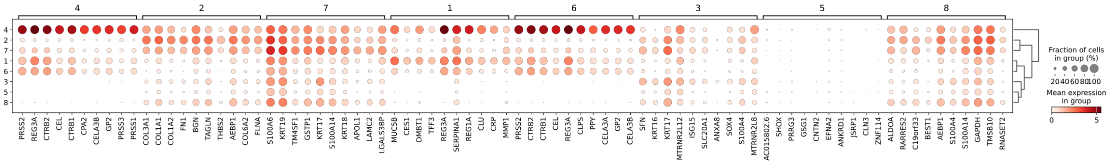
4 2 7 1 6 3 5 8
4 2 7 1 6 3 5 8
PRSS2 REG3A CTRB2 CEL CTRB1 CPA2 CELA3B GP2 PRSS3 PRSS1
COL3A1 COL1A1 COL1A2 FN1 BGN TAGLN THBS2 AEBP1 COL6A2 FLNA
S100A6 KRT19 TM4SF1 GSTP1 KRT17 S100A14 KRT18 APOL1 LAMC2 LGALS3BP
MUC5B CES1 DMBT1 TFF3 REG3A SERPINA1 REG1A CLU CRP MMP1
PRSS2 CTRB2 CTRB1 CEL REG3A CLPS PPY CELA3A GP2 CELA3B
SFN KRT16 KRT17 MTRNR2L12 ISG15 SLC20A1 ANXA8 SOX4 S100A4 MTRNR2L8
AC015802.6 SHOX PRRG3 GSG1 CNTN2 EFNA2 ANKRD1 JSRP1 CLN3 ZNF114
ALDOA RARRES2 C19orf33 BEST1 AEBP1 S100A4 S100A14 GAPDH TMSB10 RNASET2
Fraction of cells in group (%)
20 40 60 80 100
Mean expression in group
0 5

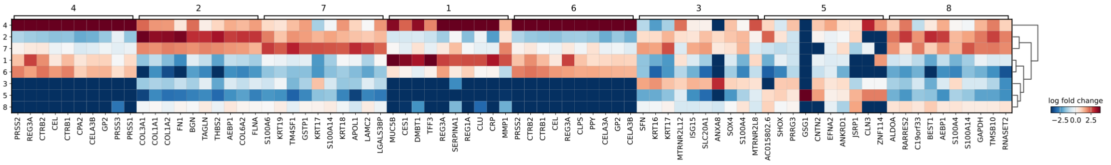
4 2 7 1 6 3 5 8
4 2 7 1 6 3 5 8
PRSS2 REG3A CTRB2 CEL CTRB1 CPA2 CELA3B GP2 PRSS3 PRSS1
COL3A1 COL1A1 COL1A2 FN1 BGN TAGLN THBS2 AEBP1 COL6A2 FLNA
S100A6 KRT19 TM4SF1 GSTP1 KRT17 S100A14 KRT18 APOL1 LAMC2 LGALS3BP
MUC5B CES1 DMBT1 TFF3 REG3A SERPINA1 REG1A CLU CRP MMP1
PRSS2 CTRB2 CTRB1 CEL REG3A CLPS PPY CELA3A GP2 CELA3B
SFN KRT16 KRT17 MTRNR2L12 ISG15 SLC20A1 ANXA8 SOX4 S100A4 MTRNR2L8
AC015802.6 SHOX PRRG3 GSG1 CNTN2 EFNA2 ANKRD1 JSRP1 CLN3 ZNF114
ALDOA RARRES2 C19orf33 BEST1 AEBP1 S100A4 S100A14 GAPDH TMSB10 RNASET2
log fold change
−2.5 0.0 2.5

**Fig. S54 Results of spatially differentially expressed genes analysis.** Top panel: dot plot of the average gene expression level and expressed proportion in each domain; Bottom panel: matrix plot of the average log fold changes of the gene expression in each domain when compared to the rest all domains.

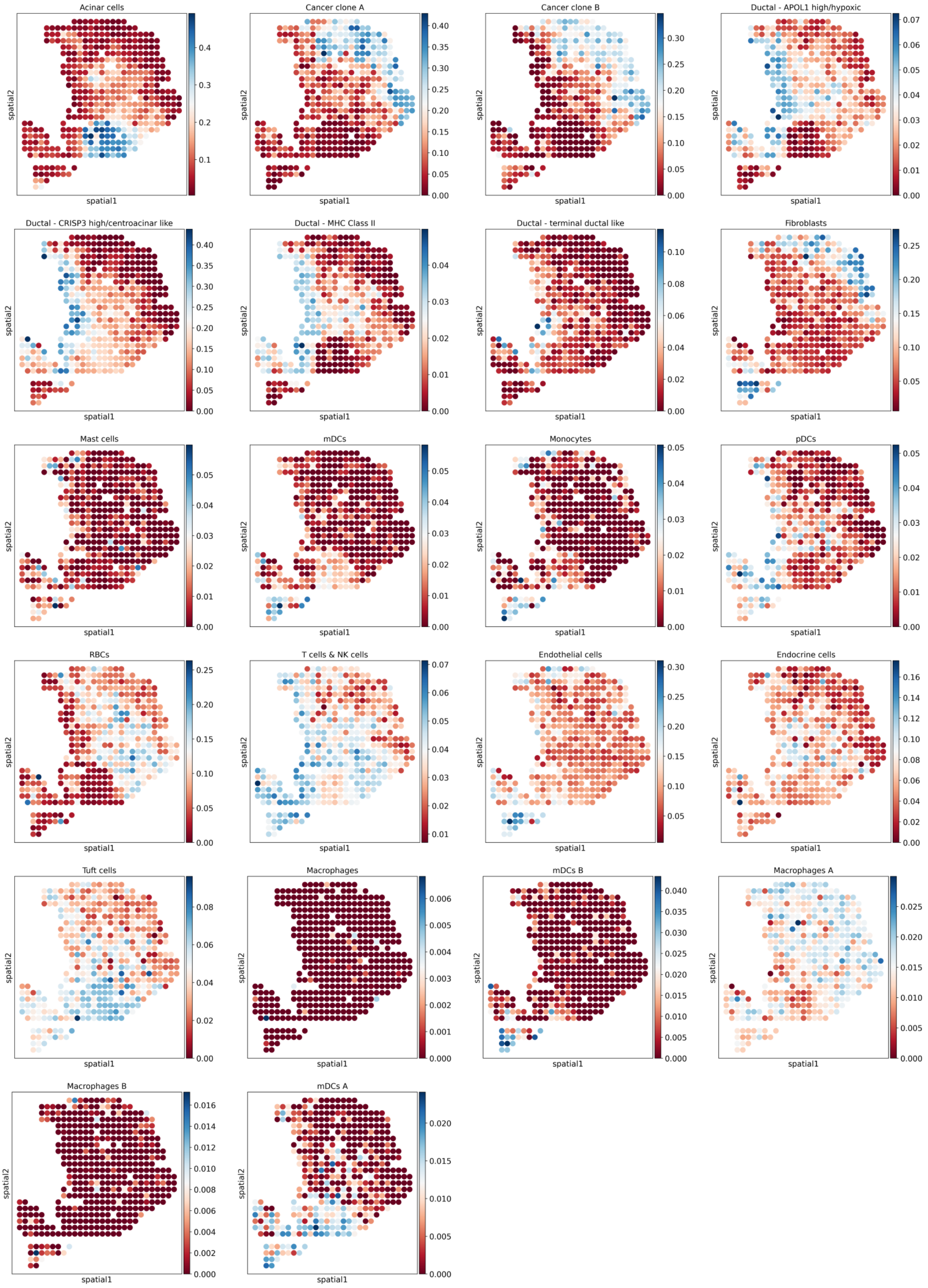
Acinar cells
Cancer clone A
Cancer clone B
Ductal - APOL1 high/hypoxic
Ductal - CRISP3 high/centroacinar like
Ductal - MHC Class II
Ductal - terminal ductal like
Fibroblasts
Mast cells
mDCs
Monocytes
pDCs
RBCs
T cells & NK cells
Endothelial cells
Endocrine cells
Tuft cells
Macrophages
mDCs B
Macrophages A
Macrophages B
mDCs A
spatial1
spatial2

**Fig. S55 Cell-type deconvolution results of PDAC-A-1.**

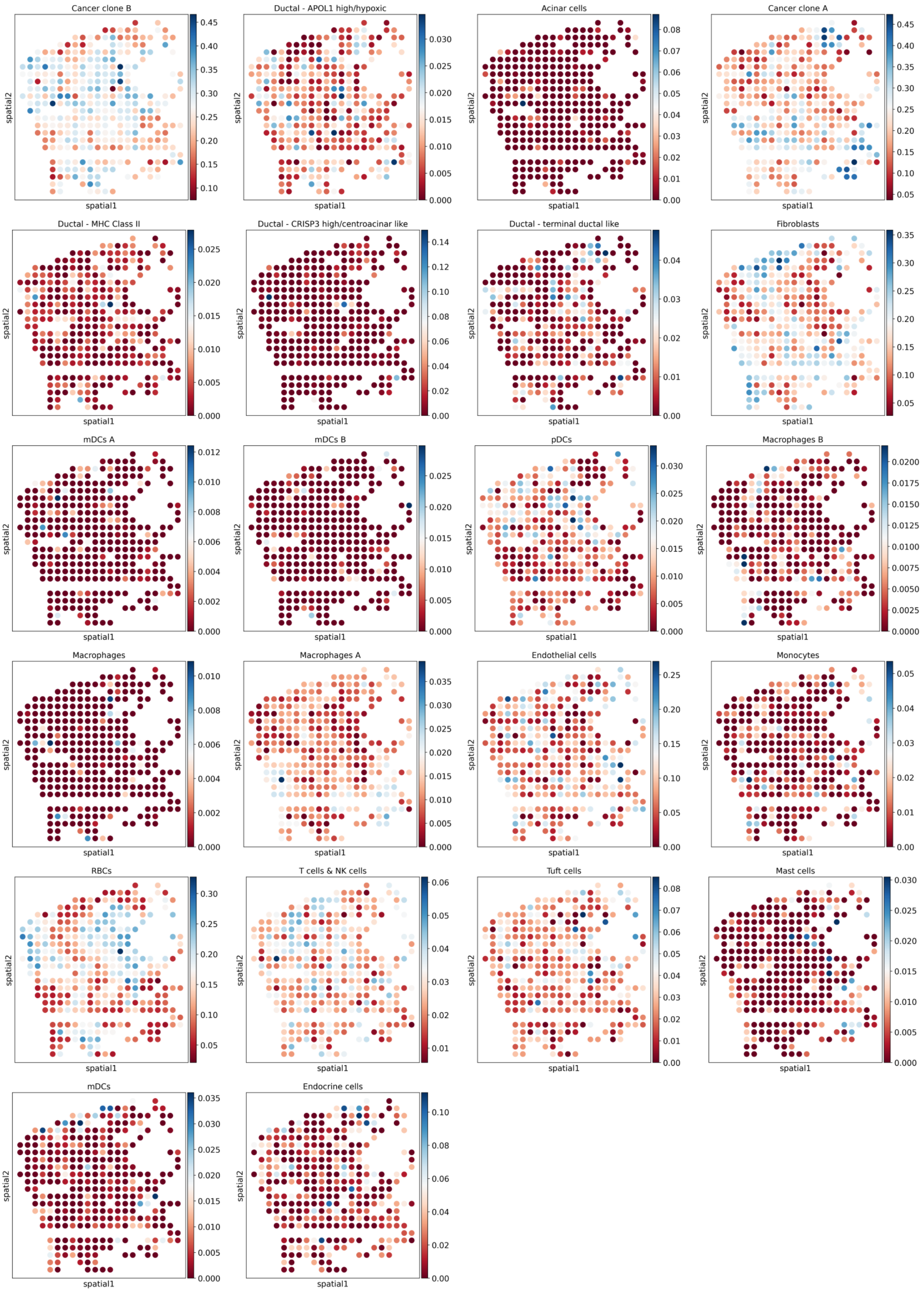
Cancer clone B
0.45 0.40 0.35 0.30 0.25 0.20 0.15 0.10
Ductal - APOL1 high/hypoxic
0.030 0.025 0.020 0.015 0.010 0.005 0.000
Acinar cells
0.08 0.07 0.06 0.05 0.04 0.03 0.02 0.01 0.00
Cancer clone A
0.45 0.40 0.35 0.30 0.25 0.20 0.15 0.10 0.05
Ductal - MHC Class II
0.025 0.020 0.015 0.010 0.005 0.000
Ductal - CRISP3 high/centroacinar like
0.14 0.12 0.10 0.08 0.06 0.04 0.02 0.00
Ductal - terminal ductal like
0.04 0.03 0.02 0.01 0.00
Fibroblasts
0.35 0.30 0.25 0.20 0.15 0.10 0.05
mDCs A
0.012 0.010 0.008 0.006 0.004 0.002 0.000
mDCs B
0.025 0.020 0.015 0.010 0.005 0.000
pDCs
0.030 0.025 0.020 0.015 0.010 0.005 0.000
Macrophages B
0.0200 0.0175 0.0150 0.0125 0.0100 0.0075 0.0050 0.0025 0.0000
Macrophages
0.010 0.008 0.006 0.004 0.002 0.000
Macrophages A
0.035 0.030 0.025 0.020 0.015 0.010 0.005 0.000
Endothelial cells
0.25 0.20 0.15 0.10 0.05 0.00
Monocytes
0.05 0.04 0.03 0.02 0.01 0.00
RBCs
0.30 0.25 0.20 0.15 0.10 0.05
T cells & NK cells
0.06 0.05 0.04 0.03 0.02 0.01
Tuft cells
0.08 0.07 0.06 0.05 0.04 0.03 0.02 0.01 0.00
Mast cells
0.030 0.025 0.020 0.015 0.010 0.005 0.000
mDCs
0.035 0.030 0.025 0.020 0.015 0.010 0.005 0.000
Endocrine cells
0.10 0.08 0.06 0.04 0.02 0.00
spatial1
spatial2

**Fig. S56 Cell-type deconvolution results of PDAC-A-2.**

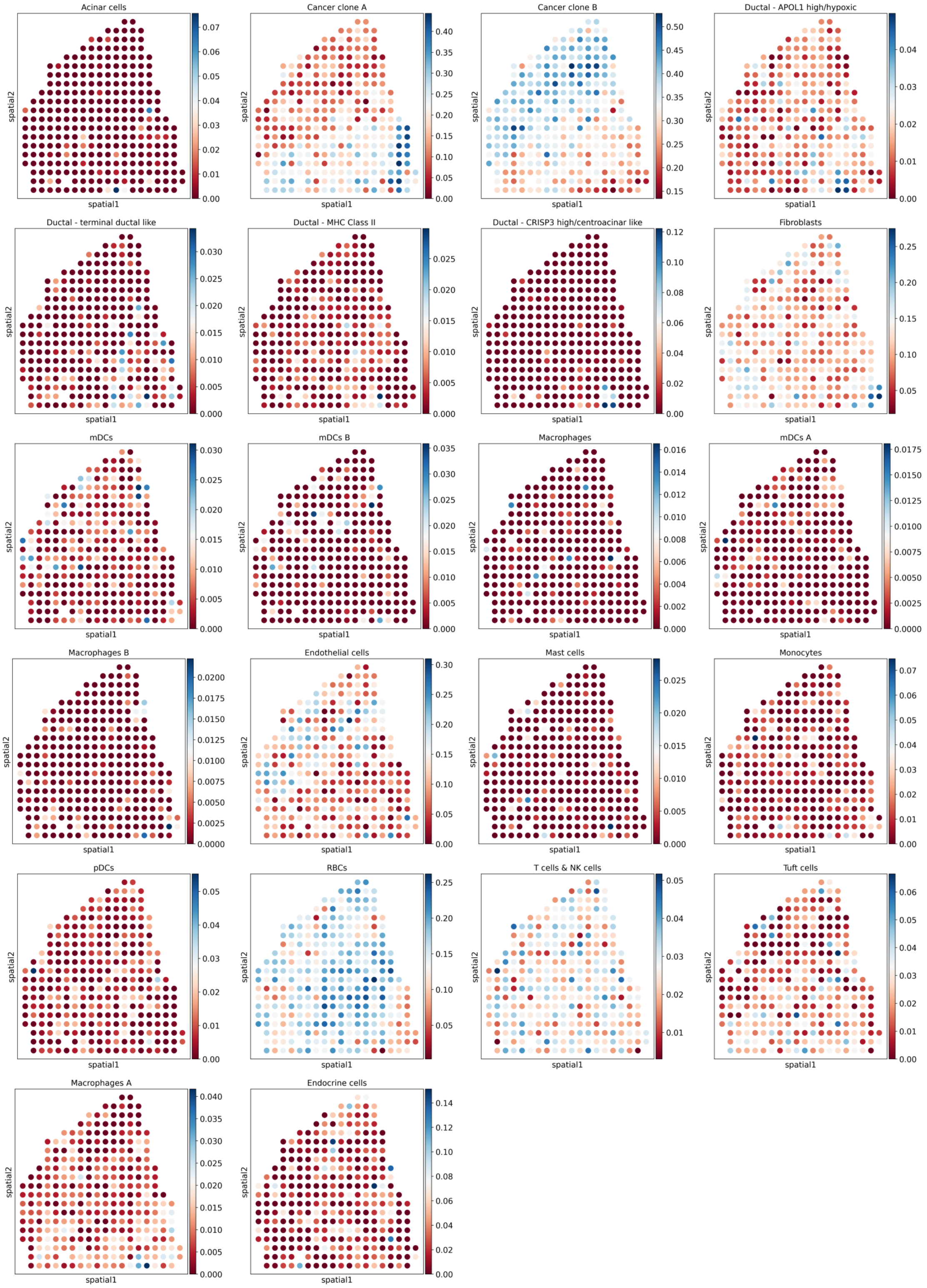

Acinar cells
Cancer clone A
Cancer clone B
Ductal - APOL1 high/hypoxic
Ductal - terminal ductal like
Ductal - MHC Class II
Ductal - CRISP3 high/centroacinar like
Fibroblasts
mDCs
mDCs B
Macrophages
mDCs A
Macrophages B
Endothelial cells
Mast cells
Monocytes
pDCs
RBCs
T cells & NK cells
Tuft cells
Macrophages A
Endocrine cells
spatial1
spatial2

**Fig. S57 Cell-type deconvolution results of PDAC-A-3.**

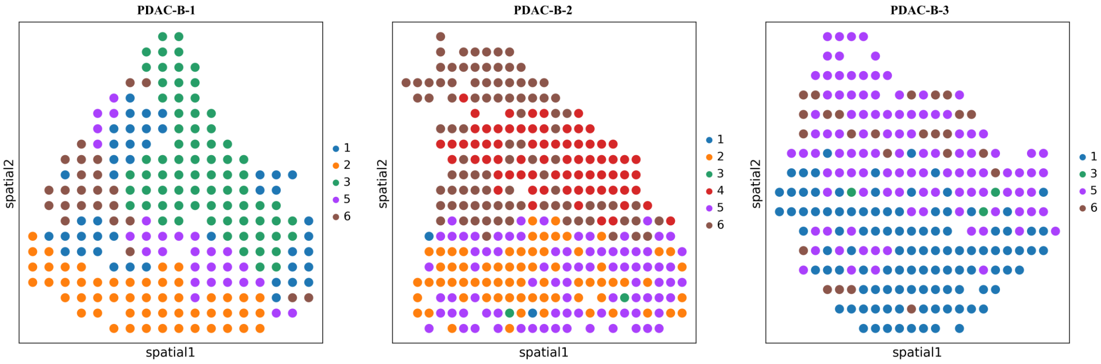
PDAC-B-1
PDAC-B-2
PDAC-B-3
spatial1
spatial2
1
2
3
4
5
6

**Fig. S58 Identified spatial domains across 3 sections in PDAC-B dataset.** Distinct spatial domains across sections were successfully identified.

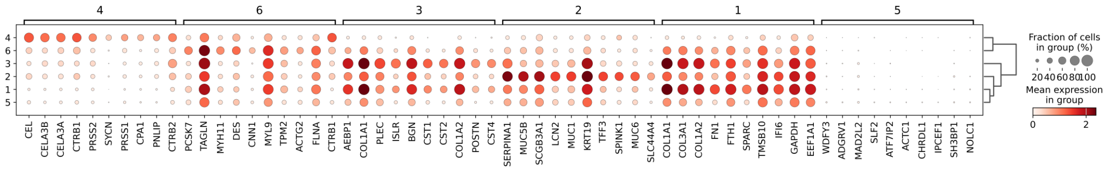
4
6
3
2
1
5
CEL
CELA3B
CELA3A
CTRB1
PRSS2
SYCN
PRSS1
CPA1
PNLIP
CTRB2
PCSK7
TAGLN
MYH11
DES
CNN1
MYL9
TPM2
ACTG2
FLNA
CTRB1
AEBP1
COL1A1
PLEC
ISLR
BGN
CST1
CST2
COL1A2
POSTN
CST4
SERPINA1
MUC5B
SCGB3A1
LCN2
MUC1
KRT19
TFF3
SPINK1
MUC6
SLC44A4
COL1A1
COL3A1
COL1A2
FN1
FTH1
SPARC
TMSB10
IFI6
GAPDH
EEF1A1
WDFY3
ADGRV1
MAD2L2
SLF2
ATF7IP2
ACTC1
CHRDL1
IPCEF1
SH3BP1
NOLC1
Fraction of cells
in group (%)
20406080100
Mean expression
in group
0
2

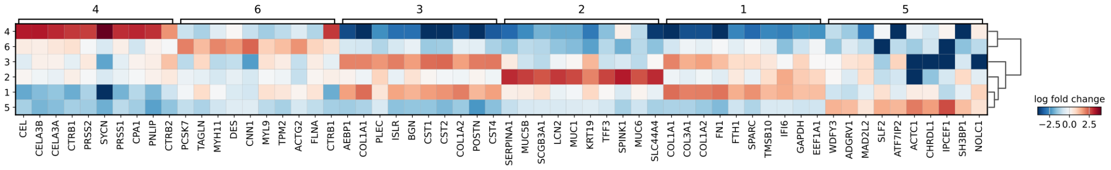
4
6
3
2
1
5
CEL
CELA3B
CELA3A
CTRB1
PRSS2
SYCN
PRSS1
CPA1
PNLIP
CTRB2
PCSK7
TAGLN
MYH11
DES
CNN1
MYL9
TPM2
ACTG2
FLNA
CTRB1
AEBP1
COL1A1
PLEC
ISLR
BGN
CST1
CST2
COL1A2
POSTN
CST4
SERPINA1
MUC5B
SCGB3A1
LCN2
MUC1
KRT19
TFF3
SPINK1
MUC6
SLC44A4
COL1A1
COL3A1
COL1A2
FN1
FTH1
SPARC
TMSB10
IFI6
GAPDH
EEF1A1
WDFY3
ADGRV1
MAD2L2
SLF2
ATF7IP2
ACTC1
CHRDL1
IPCEF1
SH3BP1
NOLC1
log fold change
−2.5 0.0 2.5

**Fig. S59 Results of spatially differentially expressed genes analysis.** Top panel: dot plot of the average gene expression level and expressed proportion in each domain; Bottom panel: matrix plot of the average log fold changes of the gene expression in each domain when compared to the rest all domains.

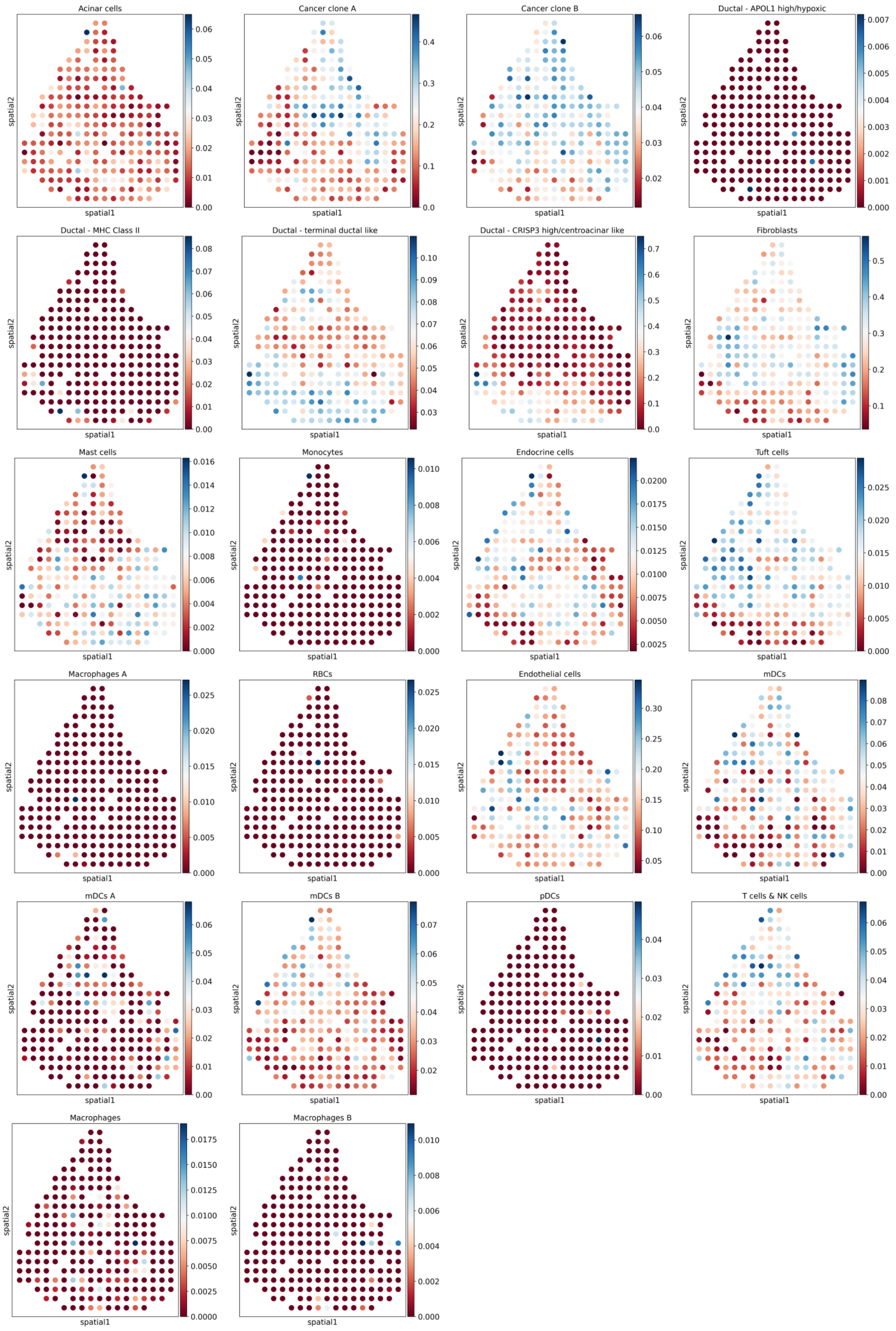
Acinar cells
spatial2
spatial1
0.00 0.01 0.02 0.03 0.04 0.05 0.06
Cancer clone A
0.0 0.1 0.2 0.3 0.4
Cancer clone B
0.02 0.03 0.04 0.05 0.06
Ductal - APOL1 high/hypoxic
0.000 0.001 0.002 0.003 0.004 0.005 0.006 0.007
Ductal - MHC Class II
0.00 0.01 0.02 0.03 0.04 0.05 0.06 0.07 0.08
Ductal - terminal ductal like
0.03 0.04 0.05 0.06 0.07 0.08 0.09 0.10
Ductal - CRISP3 high/centroacinar like
0.0 0.1 0.2 0.3 0.4 0.5 0.6 0.7
Fibroblasts
0.1 0.2 0.3 0.4 0.5
Mast cells
0.000 0.002 0.004 0.006 0.008 0.010 0.012 0.014 0.016
Monocytes
0.000 0.002 0.004 0.006 0.008 0.010
Endocrine cells
0.0025 0.0050 0.0075 0.0100 0.0125 0.0150 0.0175 0.0200
Tuft cells
0.000 0.005 0.010 0.015 0.020 0.025
Macrophages A
0.000 0.005 0.010 0.015 0.020 0.025
RBCs
0.000 0.005 0.010 0.015 0.020 0.025
Endothelial cells
0.05 0.10 0.15 0.20 0.25 0.30
mDCs
0.00 0.01 0.02 0.03 0.04 0.05 0.06 0.07 0.08
mDCs A
0.00 0.01 0.02 0.03 0.04 0.05 0.06
mDCs B
0.02 0.03 0.04 0.05 0.06 0.07
pDCs
0.00 0.01 0.02 0.03 0.04
T cells & NK cells
0.00 0.01 0.02 0.03 0.04 0.05 0.06
Macrophages
0.0000 0.0025 0.0050 0.0075 0.0100 0.0125 0.0150 0.0175
Macrophages B
0.000 0.002 0.004 0.006 0.008 0.010

**Fig. S60 Cell-type deconvolution results of PDAC-B-1.**

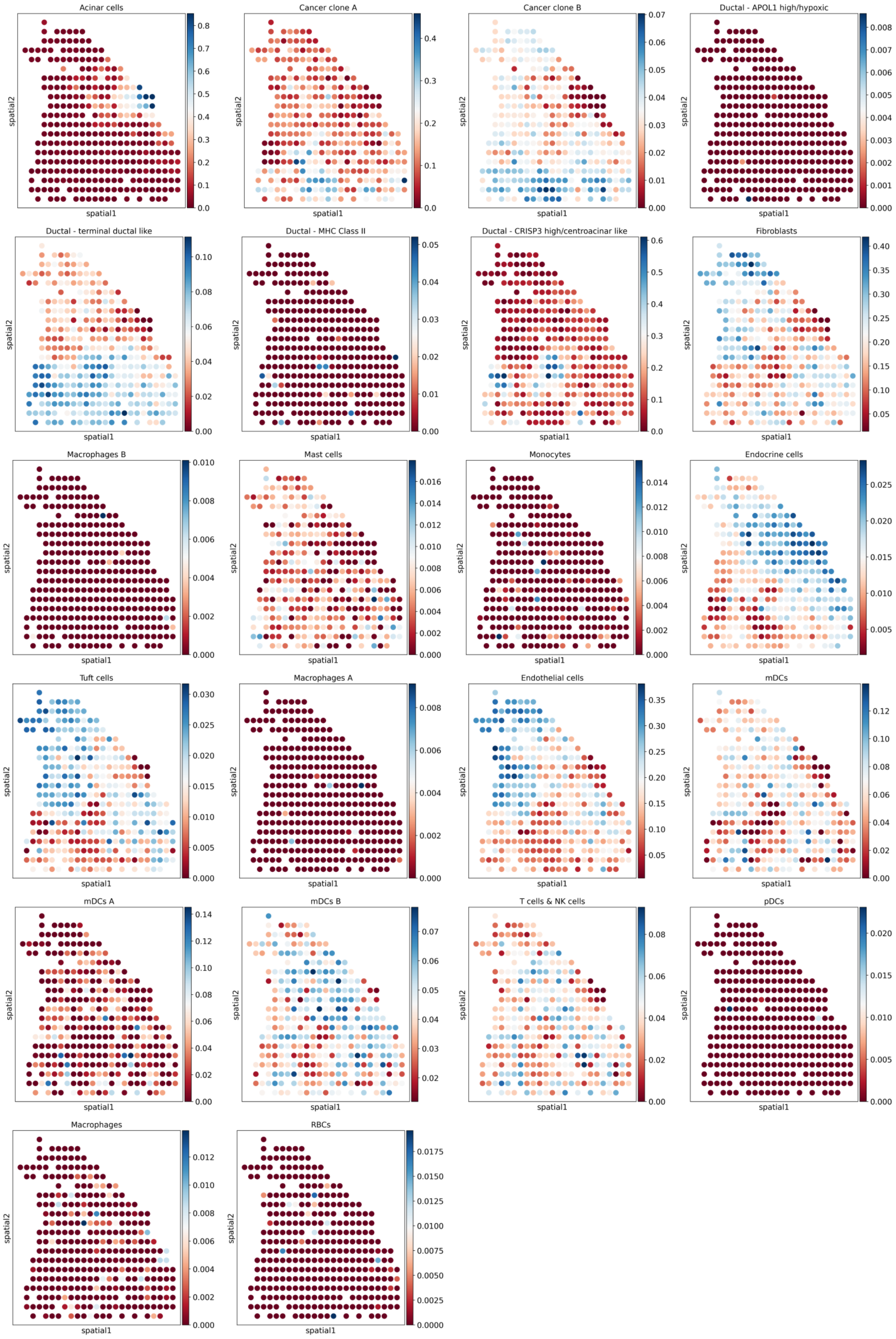
Acinar cells
Cancer clone A
Cancer clone B
Ductal - APOL1 high/hypoxic
Ductal - terminal ductal like
Ductal - MHC Class II
Ductal - CRISP3 high/centroacinar like
Fibroblasts
Macrophages B
Mast cells
Monocytes
Endocrine cells
Tuft cells
Macrophages A
Endothelial cells
mDCs
mDCs A
mDCs B
T cells & NK cells
pDCs
Macrophages
RBCs
spatial1
spatial2

**Fig. S61 Cell-type deconvolution results of PDAC-B-2.**

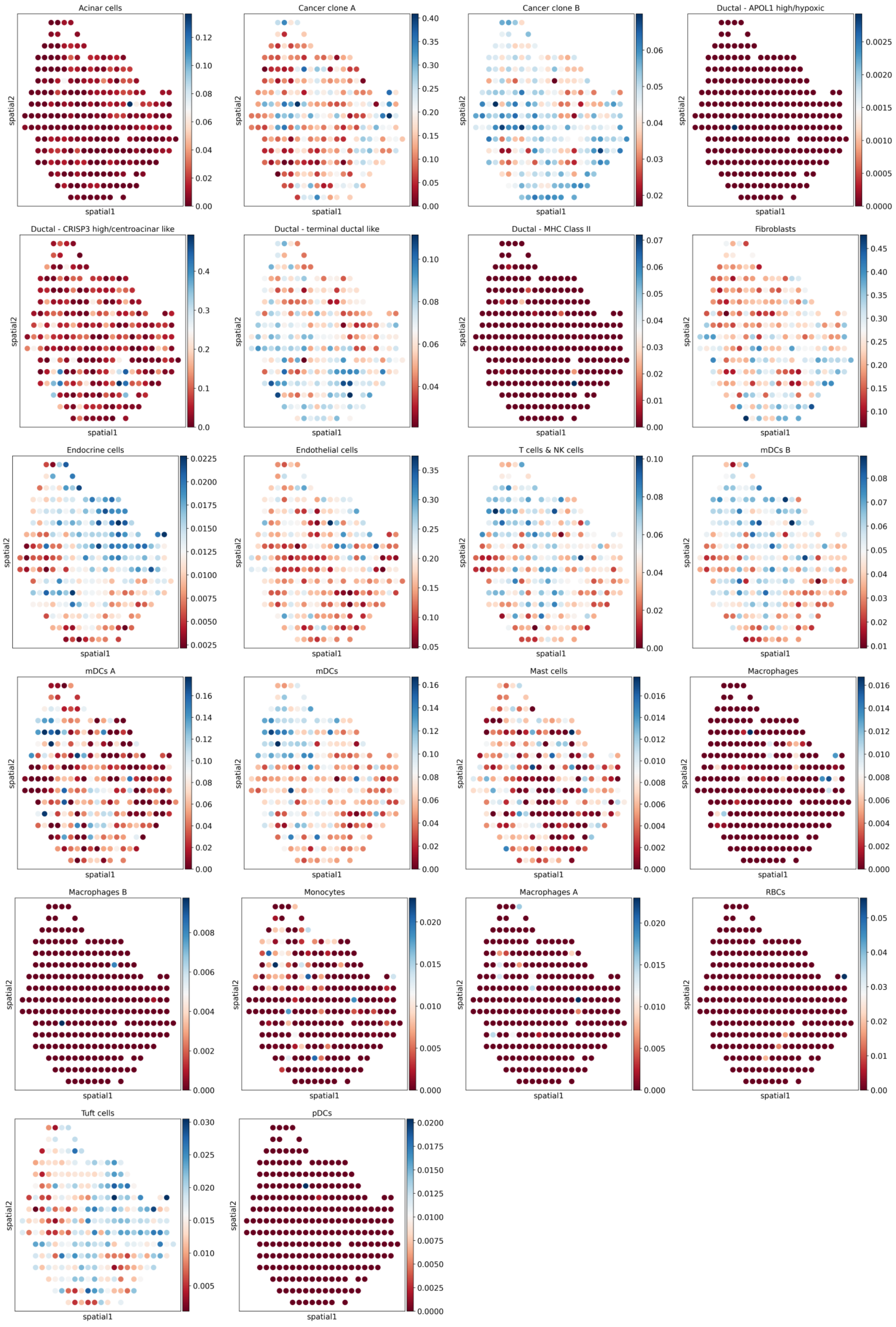
Acinar cells
Cancer clone A
Cancer clone B
Ductal - APOL1 high/hypoxic
Ductal - CRISP3 high/centroacinar like
Ductal - terminal ductal like
Ductal - MHC Class II
Fibroblasts
Endocrine cells
Endothelial cells
T cells & NK cells
mDCs B
mDCs A
mDCs
Mast cells
Macrophages
Macrophages B
Monocytes
Macrophages A
RBCs
Tuft cells
pDCs
spatial1
spatial2

**Fig. S62 Cell-type deconvolution results of PDAC-B-3.**

### 4 Supplementary Tables

| | NMI_cluster/label | ARI_cluster/label | ASW_label | ASW_label/batch | PCR_batch | isolated_label_F1 | isolated_label_silhouette | graph_conn | kBET | iLISI | cLISI |
|---|---|---|---|---|---|---|---|---|---|---|---|
| **Human & Mouse Immune cell STEP-anchord** | 0.863 | 0.874 | 0.518 | 0.974 | 0.942 | 0.788 | 0.544 | 0.872 | 0.804 | 0.286 | 0.991 |
| **Human & Mouse Immune cell STEP** | 0.802 | 0.772 | 0.564 | 0.891 | 0.871 | 0.835 | 0.624 | 0.952 | 0.206 | 0.181 | 0.997 |
| **Human Immune cell STEP** | 0.802 | 0.763 | 0.565 | 0.889 | 0.836 | 0.828 | 0.639 | 0.912 | 0.186 | 0.180 | 0.997 |
| **Human Immune cell STEP-anchord** | 0.994 | 0.998 | 0.594 | 0.968 | 0.893 | 0.979 | 0.675 | 0.915 | 0.800 | 0.306 | 1.000 |
| **Human Lung Atlas STEP-anchord** | 0.999 | 0.999 | 0.596 | 0.909 | 0.663 | 1.000 | 0.665 | 0.912 | 0.384 | 0.162 | 1.000 |
| **Human Lung Atlas STEP** | 0.732 | 0.613 | 0.545 | 0.901 | 0.731 | 0.654 | 0.587 | 0.887 | 0.384 | 0.104 | 0.996 |
| **Human Pancreas STEP-anchord** | 0.921 | 0.956 | 0.515 | 0.939 | 0.967 | 0.240 | 0.553 | 0.885 | 0.794 | 0.443 | 0.992 |
| **Human Pancreas STEP** | 0.884 | 0.930 | 0.603 | 0.873 | 0.664 | 0.182 | 0.662 | 0.916 | 0.347 | 0.219 | 1.000 |
| **Simulation 1 STEP** | 1.000 | 1.000 | 0.648 | 0.958 | 0.929 | 1.000 | 0.729 | 0.972 | 0.654 | 0.498 | 1.000 |
| **Simulation 1 STEP-anchord** | 1.000 | 1.000 | 0.633 | 0.989 | 0.927 | 1.000 | 0.728 | 0.914 | 0.965 | 0.531 | 1.000 |
| **Simulation 2 STEP-anchord** | 0.977 | 0.989 | 0.529 | 0.995 | 0.981 | 0.996 | 0.530 | 0.881 | 0.956 | 0.686 | 0.941 |
| **Simulation 2 STEP** | 0.792 | 0.639 | 0.683 | 0.786 | 0.962 | 0.858 | 0.683 | 0.760 | 0.017 | 0.183 | 0.934 |

**Table S1 Metrics values of STEP in each scRNA-seq dataset.**

| Datasets | # Batches/Sections | # cells | # hvgs | # gene modules | Hidden dimensions | Module dimensions | Epochs | Batch size | Beta | Execution time |
|---|---|---|---|---|---|---|---|---|---|---|
| **Human & Mouse Immune cell** | 33 | 97952 | 2000 | 32 | 64 | 30 | 400 | 1024 | 0.01 | 17:29 |
| **Human Immune cell** | 10 | 33056 | 2000 | 32 | 64 | 30 | 262 | 1024 | 0.01 | 10:24 |
| **Human Lung Atlas** | 16 | 32472 | 2000 | 32 | 64 | 30 | 400 | 1024 | 0.001 | 16:59 |
| **Human Pancreas** | 9 | 16382 | 2000 | 32 | 64 | 30 | 300 | 1024 | 0.03 | 05:37 |
| **Simulation 1** | 16 | 19318 | 2000 | 32 | 64 | 30 | 400 | 1024 | 0.01 | 06:26 |
| **Simulation 2** | 6 | 12097 | 2000 | 32 | 64 | 30 | 300 | 1024 | 0.001 | 07:26 |
| **MERFISH Animal 1** | 5 | 28317 | All genes used (166) | 20 | 64 | 30 | 400 | 2048 | 0.01 | 08:51 |
| **Human Colorectal Cancer (16um) HD** | 1 | 136984 | 3000 | 32 | 64 | 30 | 202 | 2048 | 0.001 | 18:05 |
| **Mouse Intestine (8um) HD** | 1 | 350107 | 2000 | 32 | 64 | 30 | 205 | 4096 | 0.01 | 52:39 |

**Table S2 Hyper parameter settings and execution time in the tasks of scRNA-seq data integration. The experiments ran on the RTX 3060 device.** The execution time is formatted as minutes: seconds.

| Datasets | # Batches/ Sections | # spots/cells | # hvgs | # gene modules | Hidden dimensions | Module dimensions | Batch Embedding Dimensions | Iterations | Node Smapling amount/rate | Graph batch-size | Beta | # GC layers | Execution time |
|---|---|---|---|---|---|---|---|---|---|---|---|---|---|
| **Human Colorectal Cancer (16um) HD** | 1 | 136984 | 3000 | 32 | 64 | 30 | 30 | 4000 | 2048 | 1 | 0.0001 | 3 | 06:14 |
| **Mouse Intestine (8um) HD** | 1 | 350107 | 2000 | 32 | 64 | 30 | 30 | 2048 | 1024 | 1 | 0.01 | 3 | 02:27 |
| **DLPFC P1 (151507 to 151510)** | 4 | 18033 | 2000 | 20 | 64 | 30 | 64 | 800 | 1 | 1 | 0.001 | 6 | 02:20 |
| **DLPFC P2 (151669 to 151672)** | 4 | 15284 | 2000 | 10 | 64 | 30 | 30 | 800 | 1 | 1 | 0.01 | 6 | 02:19 |
| **DLPFC P3 (151673 to 151676)** | 4 | 14364 | 2000 | 10 | 64 | 30 | 30 | 2000 | 1 | 2 | 0.01 | 6 | 04:03 |
| **Mouse Hypothalamus Animal 1** | 5 | 28317 | 161 | 20 | 64 | 30 | 30 | 2000 | 1 | 1 | 0.01 | 4 | 04:22 |
| **Mouse Hypothalamus Animal 7** | 12 | 70948 | 161 | 12 | 64 | 30 | 30 | 1600 | 1 | 4 | 0.001 | 4 | 07:35 |
| **Mouse Medial Prefrontal Cortex** | 3 | 3188 | 166 | 8 | 40 | 20 | 30 | 1600 | 1 | 1 | 0.01 | 4 | 00:37 |
| **Mouse Brain Sagittal Anterior & Posterior** | 2 | 6050 | 3000 | 16 | 40 | 30 | 30 | 800 | 1 | 2 | 0.01 | 3 | 04:09 |
| **Human Normal and BA Liver** | 6 | 15120 | 2000 | 20 | 64 | 30 | 30 | 2200 | 0.7 | 3 | 0.01 | 2 | 06:13 |

**Table S3 Hyper parameter settings and execution time in the tasks of SRT data integration. The experiments ran on the RTX A6000 device except the experiment of Mouse Brain Sagittal Anterior & Posterior, which ran on RTX 3060.** The execution time is formatted as minutes: seconds.

| Datasets | | # Batches/Sections | # spots/cells | # cell types | # hvgs | # gene modules | Hidden dimensions | Module dimensions | Batch Embedding Dimensions | Epochs | Batch size | Beta | # GC layers | Execution time |
|---|---|---|---|---|---|---|---|---|---|---|---|---|---|---|
| **Simulation Study** | scRNA-seq reference | 10 | 60337 | 8 | 2000 | 32 | 64 | 30 | 64 | 400 | 1024 | 0.01 | / | 26:44 |
| | SRT | 1 | 3592 | | | | 64 | 30 | 30 | 800; 1000; 2400 | / | | 2 | 01:18; 00:47; 03:50 |
| **Human Lymph Node** | scRNA-seq reference | 4 | 73260 | 34 | 2000 | 32 | 64 | 30 | 30 | 334 | 2048 | 0.01 | / | 27:23 |
| | SRT | 1 | 4035 | | | | 64 | 30 | 30 | 800; 800; 2000 | / | | 1 | 00:58; 00:47; 05:14 |
| **Human Normal and BA Liver** | snRNA-seq reference | 2 | 20324 | 12 | 2000 | 32 | 64 | 30 | 30 | 400 | 1024 | 0.01 | / | 07:00 |
| | SRT | 4 | 14431 | | | | 64 | 30 | 30 | /; /; 2000 | / | | / | /; /; 12:11 |
| **PDAC-A** | scRNA-seq reference | 2 | 3659 | 22 | 2000 | 20 | 64 | 30 | 30 | 300 | 512 | 0.01 | / | 14:50 |
| | SRT | 3 | 1032 | | | | 64 | 30 | 30 | 400; 400; 1500 | 3 | | 3 | 00:16; 00:12; 01:28 |
| **PDAC-B** | scRNA-seq reference | 2 | 3659 | 22 | 2000 | 20 | 64 | 30 | 30 | 300 | 512 | 0.01 | / | 01:15 |
| | SRT | 3 | 787 | | | | 64 | 30 | 30 | 800; 800; 1500 | 3 | | 3 | 00:09; 00:14; 01:05 |

**Table S4 Hyper parameter settings and execution time in the tasks of cross modality integration. The first two tests (simulation study and human lymph node) ran on the RTX A6000 device, while the last two tests ran on RTX 3090 device.** The execution time is formatted as minutes: seconds.
